# A mixture of endocrine disrupting chemicals linked to lower birth weight induces adipogenesis and transcriptional changes related to birth weight alterations and diabetes

**DOI:** 10.1101/2025.02.13.638050

**Authors:** Polina Lizunkova, Nicolò Caporale, Elin Engdahl, Cristina Cheroni, Pierre-Luc Germain, Gábor Borbély, Christian Lindh, Chris Gennings, Carl-Gustaf Bornehag, Giuseppe Testa, Joëlle Rüegg

## Abstract

There is increasing evidence that endocrine disrupting chemicals (EDCs) are contributing to the rise in metabolic disorders and obesity. Humans are constantly exposed to numerous EDCs, thus human exposure entails complex EDC mixtures. In this study, we examined the effects of an EDC mixture, mixture G, composed of four phthalate esters, triclosan, and three poly- och perfluorinated alkyl substances. Mixture G had previously been defined based on its association with lower birth weight in a pregnancy cohort, where low birth weight is an early risk factor for metabolic morbidities later in life. Here, we studied its effects on adipogenesis and uncovered their underlying transcriptional changes. Human mesenchymal stem cells (hMSCs) were exposed to mixture G in concentrations and mixing ratios that reflect those measured in human serum. Mixture G induced adipogenesis in hMSCs, as evidenced by a dose-dependent increase in lipid droplet accumulation after 14-21 days. Notably, significant adipogenic effects were observed at concentrations comparable to those detected in humans. RNA-sequencing upon exposure for 48 h revealed dose-dependent transcriptional changes in over 1000 genes. Mixture G-induced differentially expressed genes (DEGs) showed significant overlap with genes involved in osteogenesis, with glucocorticoid-regulated genes, and with genes associated with birth weight alterations and diabetes type II. These results indicate that exposure to an environmentally relevant EDC mixture induces adipogenesis and leads to transcriptional alterations that might change the balance between adipogenic and osteogenic differentiation as well as the functionality of MSCs, possibly via interference with glucocorticoid signalling. Thus our findings underscore the role of EDCs as metabolic disruptors and shed light on the molecular mechanisms underlying their potential contribution to the development of metabolic disorders.

**Highlights:**

- An endocrine disruptor mixture linked to lower birth weight increases adipogenesis
- The mixture induced transcriptomic changes at low doses
- Affected genes are associated with birth weight and diabetes type II

**Graphical abstract:** 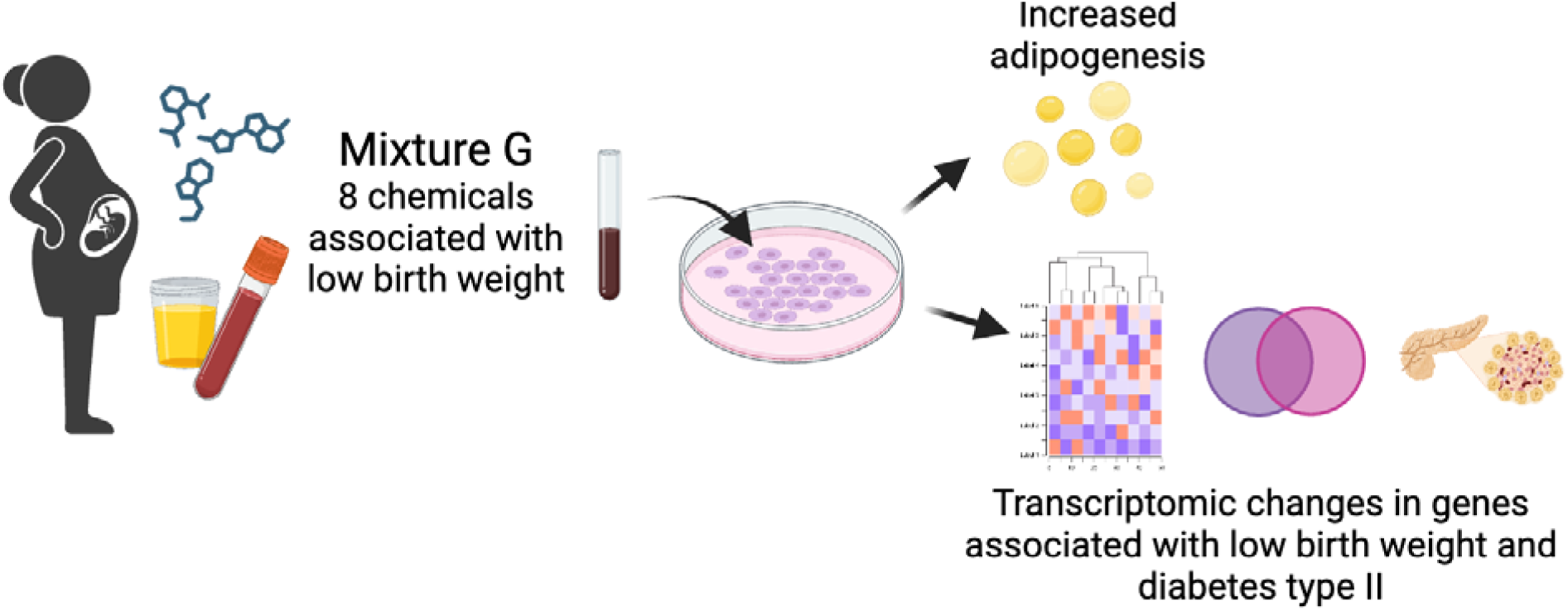

## 1. Introduction

Endocrine-disrupting chemicals (EDCs) interfere with the endocrine system, thereby leading to disruptions of physiological functions. EDCs are omnipresent in the environment and are used in a multitude of common consumer goods, such as toys, food, personal care products, and clothing (Ribeiro et al., 2017). Examples are phthalate esters found in a wide range of products such as paints, toys, cosmetics, and food packaging, or poly- and perfluoroalkyl substances (PFAS) found in a products like clothes, carpets, building materials, and food contact papers due to their water repellent properties. These and other chemicals can leach from their materials into the environment. Thus, EDCs from different chemical classes are detectable in groundwater (Banzhaf et al., 2017; Dueñas-Moreno et al., 2022; Hu et al., 2016; Kotowska et al., 2020; Norin and Strömvall, 2004; Sackaria and Elango, 2020) as well as in human blood, breast milk, amniotic liquid, and urine (Authority, 2013; Fromme et al., 2007; Wang et al., 2019). EDCs have the potential to interfere with essential and carefully regulated hormone-driven developmental processes throughout gestation, infancy, and childhood, resulting in long-lasting health consequences (Amato et al., 2021; Haverinen et al., 2021; Ko et al., 2019; Mariana and Cairrao, 2023).

In recent years several epidemiological studies have found associations between prenatal exposure to different types of EDCs and lower birth weight (LBW) (Ashley-Martin et al., 2017; Birks et al., 2016; Branda et al., 2022; Guo et al., 2014), a risk factor for adult-onset of metabolic diseases such as diabetes and obesity (Bianchi and Restrepo, 2022; Branda et al., 2022; Govarts et al., 2018; Hack et al., 1995; Nakano, 2020; Tanis et al., 2005), and related health outcomes. For example, Govarts and colleagues (2018) found associations between prenatal exposure to organochlorine compounds and PFASs and increased prevalence of small for gestational age (SGA) birth outcome using the data from seven European birth cohorts, totalling 5,446 mother-child pairs (Govarts et al., 2018). Furthermore, data from the Swedish Environmental Longitudinal, Mother and Child, Asthma and allergy (SELMA) study has shown that prenatal exposure to a combination of EDCs including phthalates, bisphenols, PFASs, organochlorine pesticides, was associated with LBW (Svensson et al., 2021). Furthermore, prenatal exposure to a combination of EDCs has been associated with alterations in infant weight gain trajectories characterized by a slower infant growth spurt rate (Svensson et al., 2021). In other studies, maternal phthalate urine concentrations, such as mono(2- ethylhexyl)phthalate (MEHP) and monobenzyl phthalate (MBzP), have been negatively associated with weight-for-length z-scores at birth, but positively with body mass increase during 3 months after birth and with Body Mass Index (BMI) z-scores at 3–4 years of age, and positively with total cholesterol and triglyceride in cord serum (Ferguson et al., 2022; Kim et al., 2016; Sol et al., 2020).

These epidemiological studies have been corroborated by experimental studies showing evidence for metabolic disruption induced by EDC exposure (Braun, 2017). For example, studies using rodent models showed that developmental exposure to phthalates like diethyl phthalate (DEP), di-(2-ethylhexyl) phthalate (DEHP) and its metabolite MEHP, resulted in metabolic changes in the offspring, such as increased body weight, insulin insensitivity, increased adipose tissue deposition, and increase in serum glucose (Campioli et al., 2014; Ding et al., 2021, 2019; Hao et al., 2013; Lee et al., 2016; Mondal and Mukherjee, 2020; Rajagopal et al., 2019a, 2019b; Rajesh and Balasubramanian, 2015). Furthermore, in *in vitro* studies EDCs such as mono isononyl cyclohexane-1,2-dicarboxylic acid ester (MINCH), MBzP, MEHP, and DEHP, have been shown to activate peroxisome proliferator-activated receptor-γ (PPARγ), the master regulator of adipogenesis, and promote adipogenesis (Biemann et al., 2012; Ellero-Simatos et al., 2011; Feige et al., 2007; Hao et al., 2012; Huang et al., 2016; Hurst and Waxman, 2003a; Schaffert et al., 2022; Useini et al., 2023).

Adipose tissue plays a central role in energy homeostasis, hence disrupting its development or function can have effects on metabolic health. Thus, studying the effects of EDCs on adipose tissue development can provide information on mechanisms underlying effects of EDCs on metabolic outcomes. Adipose tissue development is a tightly regulated process that, in mammals, starts *in utero* and relies on precise transcriptional and epigenetic coordination (Desoye and Herrera, 2021; Orsso et al., 2020). Adipose tissue originates from the mesodermal lineage. Therefore, a commonly used *in vitro* model to study adipose tissue development are mesenchymal stem cells (MSC). Human MSC (hMSCs) differentiation to adipocytes captures an essential part of human adipose tissue development occurring during the gestational period (Robert et al., 2020a). Several EDCs have been shown to induce adipogenesis in MSCs, partly via interaction with PPARγ. For example, phthalates are known to be at least partial PPARγ agonists, and MEHP was shown to induce adipogenesis in mouse bone-marrow derived MSCs (mBMMSCs) and to increase mRNA expression of Pparγ (Chiu et al., 2018; Watt and Schlezinger, 2015). Furthermore, MEHP inhibited osteoblast differentiation and mRNA expression Runx2, one of the osteogenesis inducers, in mBMMSCs (Chiu et al., 2018). Similarly, PFOA was shown to interact with PPARγ, induce adipogenesis, and inhibit osteogenesis in hMSCs (Kirk et al., 2021; Qin et al., 2022). Furthermore, benzyl butyl phthalate (BBP) has been shown to induce adipogenesis and epigenetic changes in MSCs (Sonkar et al., 2016).

Real-life exposures for humans and the environment entail simultaneous exposure to a multitude of EDCs from a variety of sources, leading to exposure to non-intentional mixtures. In such complex mixtures, chemical interactions may occur that cannot be fully captured when studying effects of single chemicals isolation. In comparison to other studies conducted solely on individual chemicals, in a previous study we have demonstrated that an EDC mixture relevant for human exposure, both in terms of concentrations and proportions, induces epigenetic changes in human MSCs (Lizunkova et al., 2022). In this study, we aimed at exploring early transcriptional responses in hMSCs induced by an EDC mixture called mixture G, which has been associated with LBW in the SELMA cohort (Svensson et al., 2021), and shown to increase adipogenesis in zebrafish (Mentor et al., 2020), in order to delineate potential mechanisms underlying long-term effects of real-life exposures.

## 2. Methods

### 2.1 Mixture G

The chemical mixture used in this study, mixture G, originated from the EU-funded Horizon 2020 project EDC-MixRisk where a whole mixture approach was used to determine the contribution of 20 analytes/compounds (measured in maternal urine and serum during first trimester), as a mixture, on birth weight in the offspring in the SELMA study. The procedure to establish chemical mixtures associated with fetal development has been described in detail elsewhere (Bornehag et al., 2019; Caporale et al., 2022). The EDCs included in mixture G, and their mixing proportions, are shown in Supplementary table 1 and consisted of monoethyl phthalate (MEP), Monobutyl Phthalate (MBP), MBzP, MEHP, Di-isononyl phthalate (DINP), Triclosan, Perfluorooctanoic acid (PFOA), Perfluorooctane sulfonate (PFOS), and Perfluorohexanesulphonic acid (PFHxS) (Mentor et al., 2020). This mixture was tested using concentrations and mixing ratios corresponding to human exposure, where 1X denotes the geometric mean of exposure levels in Swedish pregnant women included in the SELMA study (Bornehag et al., 2019).

### 2.2 Human mesenchymal stem cell culture

The molecular impact of mixture G on two growth and metabolism-relevant human cell models: adult bone marrow-derived mesenchymal stem cells (bmMSCs) and iPSC-derived mesenchymal stem cells (hiPSC-MSCs).

bmMSCs obtained from 2 donors were a kind gift of Dr. Katarina Leblanc (Center of Hematology and Regenerative Medicine, Department of Medicine, Karolinska Institutet, Sweden). They were cultured at 37°C, 5% CO2, in growth media (GM): Dulbecco’s Modified Eagle’s medium (DMEM, Gibco ®) supplemented with 10% heat-inactivated FBS (Gibco®), 1% penicillin/streptomycin (Gibco®) and 2% L-glutamine (Gibco®). The hiPSC-MSCs were obtained and cultured from neural crest stem cells derived from iPSCs according to the protocol described by (Menendez et al., 2013) and previously used in (Adamo et al., 2015). hiPSC-MSCs were cultured under the same conditions but in DMEM/F12 (Gibco®) medium supplemented with 10% heat inactivated FBS, 1% penicillin/streptomycin and 1% L-glutamine. All cultures were routinely tested for mycoplasma. During experiments, 10% FBS was exchanged to 10% charcoal stripped FBS (Gibco®), hereafter referred to as treatment media.

### 2.3 Lipid droplet accumulation assay

For adipogenic induction and lipid accumulation assessment, cells were seeded in 96 black-walled µCLEAR well plates and treatments were performed in treatment media containing mixture G (0.1-1000X human exposure) or DMSO as solvent control for 14-21 days where the medium was changed twice a week. Lipid accumulation in adult hMSCs was assessed by imaging as previously described by Vemuri et al (Vemuri et al., 2011) with small modifications to the procedure. Briefly, cells were exposed to DMSO (1:1000) or mixture G in six replicate wells for 14-21 days. The lipid droplets and cell nuclei were stained by either directly replacing the treatment media with 100µl media containing 10µg/mL BODIPY 493/503 (Gibco®) and 2 µg/mL of Hoechst 33342 (Gibco®) or by first fixating the cells in 4% PFA followed by the staining. After 1 hour of incubation at 37°C, 5% CO2, or at room temperature for PFA-fixed cells, the wells were washed with PBS and images were acquired immediately using the Image Xpress Micro High-Content Analysis System. Images were taken in FITC and DAPI channels at 10x magnification, at 16 sites per well. Images were further analyzed with the MetaXpress High-Content Image Acquisition and Analysis software (Molecular Devices, Sunnyvale California USA). Using the Transfluor HT analysis module, lipid droplets were quantified by measuring the integrated granule intensity and this value was normalized to nuclei count. For each treatment condition, the lipid accumulation per cell is presented as a ratio compared to the lipid accumulation per cell of the DMSO control on the same plate.

### 2.4 RNA seq

Cells were seeded in 6 well plates and expanded until they reached 70-80% confluence. Growth media was replaced by treatment media two days before experiment start, and treatments were performed for 48h in this media before the cells were lysed. Mixture G concentrations tested were 0.1-1000X human exposure.

Total RNA was isolated with the RNeasy Micro kit (Qiagen) according to the manufacturer’s instructions. TruSeq Stranded Total RNA LT Sample Prep Kit (Illumina) was used to run the library for each sample. Sequencing was performed with the Illumina HiSeq 2000 platform, sequencing on average 10 million 50bp paired-end reads per sample.

### 2.5 Analyses of the RNA - Seq data

All transcriptomic data will be made publicly available on a standard repository for omics data, and all bioinformatic analyses, and the code will be released on GitHub upon acceptance to ensure reproducibility. In brief, RNA-seq quantification was performed directly from FastQ files using Salmon 0.6.1, using the hg38 Refseq annotation. Only genes with at least 20 reads in at least 2 samples were included in downstream analyses; small (<200 nt) genes, ribosomal RNA genes, and fusion genes were excluded.

### 2.6 Differentially expressed genes (DEGs)

Differential expression analysis was performed on the estimated counts after TMM normalization with edgeR (Robinson et al., 2010) using a likelihood ratio test on the coefficients of a negative binomial model including the genetic background and the mixture concentration (Robinson et al., 2010). Concentration of mixture G was treated as a categorical variable (converted to factor), and tested for any non-zero coefficient. Genes identified through this method were then kmeans-clustered on the basis of their smoothed fold change upon each concentration. The mean smoothed fold change patterns for the main cluster(s) were plotted for dose-response patterns.

### 2.7 Gene Ontology enrichment analysis

Gene Ontology (GO) over-representation analyses were performed with topGO R package (Alexa and Rahnenfuhrer, 2023) on the Biological Process domain of gene ontology, relying on Fisher test and Weight01 method; an enrichment cut-off of 2 was imposed. The analysis was performed on differentially expressed genes (DEGs), split in up- or down-regulated; a pool of tested genes (as selected for differential expression analysis) was used as background.

GO enriched terms were further examined on the complete list of DEGs (both up- and down regulated), with the goseq R package, including correction for eventual RNA-seq transcript-length bias and excluding genes without annotation. Terms with at least 10 but no more than 1,000 associated genes were considered, and Fisher’s exact test was used. Parent terms with significantly enriched children terms were filtered out to improve the specificity of the enrichments. Unless stated otherwise, other enrichment tests were performed using the hypergeometric test.

## 3. Results

### 3.1 Mixture G alters gene expression in MSCs at human relevant concentrations

We set out to confirm the ability of mixture G to induce adipogenesis in a human cell model and further investigated the early transcriptional changes induced by mixture G. In a first step, we confirmed mixture G’s ability to induce adipogenesis in bmMSCs. To this end, bmMSCs were exposed to five concentrations of mixture G (0.1X,1X,10X,100X, and 1000X the concentration corresponding to the mean of the levels in the SELMA mothers) for 14-21 days, after which lipid droplets accumulated in pre-adipocytes were stained and quantified by high content fluorescence imaging. Mixture G significantly enhances lipid droplet accumulation already at 1X exposure (Figure 1A), showing that it can promote adipogenic differentiation of bmMSCs at levels found in serum of Swedish women.

**Figure 1:**
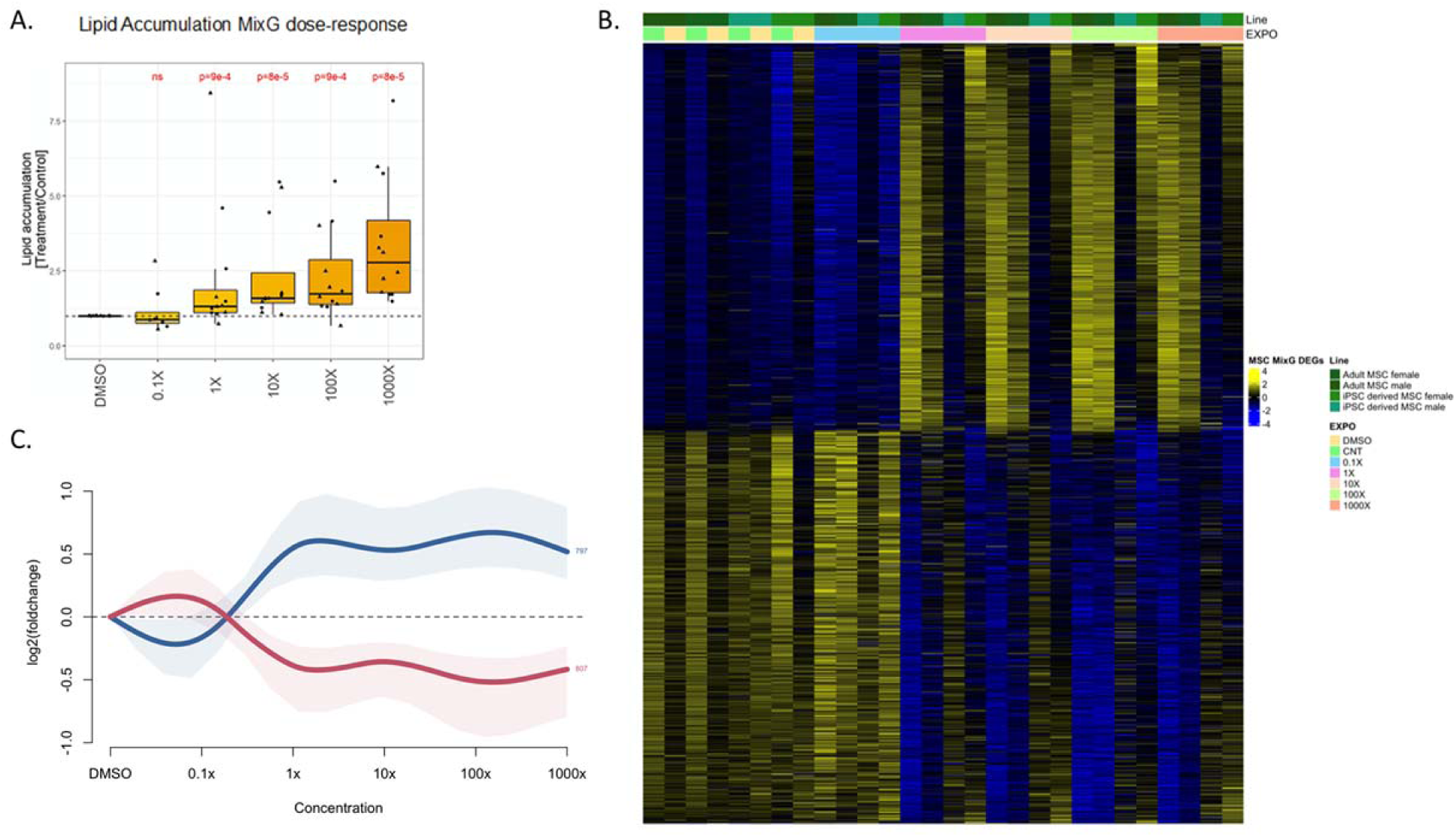
Effects of mixture G exposure on MSCs. A. Mixture G induced lipid droplet accumulation in bmMSCs. Mean and standard error per treatment is shown in the graph. * = p < 0.05 obtained by Kruskal–Wallis rank sum test followed by Dunns post hoc test adjusted using Benjamini-Hochberg when comparing with DMSO as a control. B. Heatmap of the core set of 1604 differentially expressed gene (DEGs; FDR < 0.05, logFC > 0.2, logCPM > 0; both MSC models combined) that are differentially expressed upon mixture G treatment in comparison to the control DMSO treatment. Values are log2FC computed for each cell line exposed to MIX G relative to the corresponding control DMSO treatment. C. Dose-response curves of the DEG core set over the range of mixture G concentrations showing clustering into two groups with dose-dependent increases or decreases.

Next, the short term impact of mixture G was characterized at the transcriptional level in bmMSCs as well as iPSC-derived mesenchymal stem cells (iPSC-MSC), which were included as proxies of fetal MSCs vis a vis adult-derived bmMSC. Both MSC models were exposed to the five mixture G concentrations (0.1X, 1X, 10X, 100X, 1000X) for 48 hrs and transcriptional profiling was conducted by RNA-Seq. Both MSC models, regardless of their origin, showed similar expression of hallmark MSC genes (Supplementary figure S1), grounding the rationale for a joint differential expression analysis across both cell types to identify the most robust targets of the mixture. Also, the different doses were treated as categorical (i.e. independent treatments, without assuming a dose-response) for the analysis. The identified set contained 1604 differentially expressed genes (DEGs) (Figure 1B, Supplementary figure S2, Supplementary table S2). A dose-response analysis of these DEGs identified two clusters showing a dose-dependent increase or decrease upon exposure (Figure 1C). There were slightly more down-regulated (n = 807), than upregulated (n = 797) genes, and gene expression changes in both directions followed a similar pattern whereby most of the expression changes reached their maximum already around the 1X concentration (Figure 1C). Similar dose-response curves for these core DEGs were observed when the MSC lines were analyzed individually (Supplementary Figure S3). Thus, mixture G changed expression of 1604 genes similarly in four different MSC lines of two different origins at concentrations corresponding to the mean measured in the SELMA women.

### 3.2 Mixture G induces cell proliferation and represses cell adhesion-associated pathways

Next, we performed GO enrichment analysis to identify biological processes associated with the mixture G-induced DEGs. Upregulated DEGs were enriched in 130 gene ontology (GO) terms while the downregulated DEGs were enriched in 104 terms (Supplementary table S3). The top eight GO terms of the upregulated and downregulated DEGs are shown in Figure 2. DEGs upregulated by mixture G showed significant enrichment for GO terms related to cell division and DNA repair, while enrichment of DEGs downregulated by mixture G were related to cell adhesion and extracellular matrix modulation terms.

**Figure 2:**
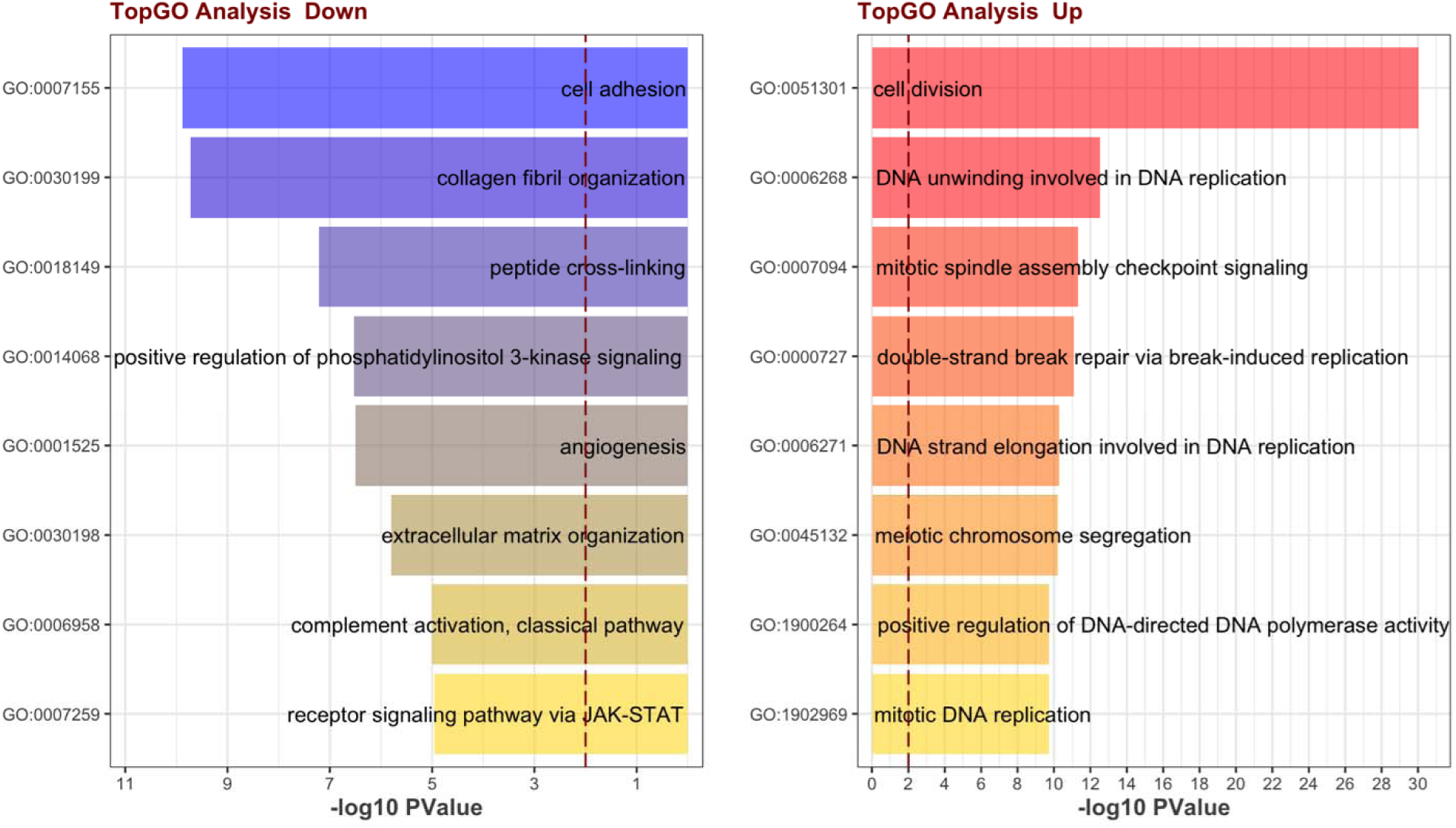
Gene Ontology over-representation analysis of the core DEGs. Top Gene Ontology (GO) terms associated with the downregulated (left) and the upregulated (right) DEGs.

While these processes are rather generic, cell adhesion (GO: 0007155) and extracellular matri organization (GO:0030198) have previously been shown to be upregulated in early osteogenesis (Assis-Ribas et al., 2018; Bosch et al., 2013; van de Peppel et al., 2017; Z et al., 2018). Thus, we investigated further if mixture G DEGs overlap with DEGs identified in early time points of adipogenesis or osteogenesis induction in MSCs from a previously published study on male human bmMSCs (Peppel et al., 2017). As shown in Figure 3, upregulated mixture G DEGs (MSC_DEGs_UP) significantly overlapped with downregulated genes upon induction of both adipogenesis and osteogenesis (all the xx_DOWN labels in the different timepoints in the x axis of Figure 3). When testing for Gene Ontology enrichment among the genes overlapping between MSC_DEGs_UP and h48_DOWN (282 genes for adipogenesis, 278 genes for osteogenesis, from (Peppel et al., 2017)), we found categories related to cell proliferation processes (Figure 4 A, B), showing that while mixture G induces proliferation related GO terms after 48 h treatment, induction of adipogenesis and osteogenesis leads to an immediate suppression of these processes. Similarly, downregulated mixture G DEGs (MSC_DEGs_DOWN) significantly overlapped with upregulated genes upon induction of both adipogenesis and osteogenesis (all the xx_UP labels in the different timepoints in the x axis of Figure 3). The overlapping genes between MSC_DEGs_DOWN and adipogenic induction (h48_UP, 149 genes from Figure 3A) include processes that are not related to adipogenesis (Figure 4 C). On the other hand, overlapping genes between MSC_DEGs_DOWN and osteogenic induction (h48_UP, 216 genes from Figure 3B) were related to the previously mentioned cell adhesion (GO: 0007155) and extracellular matrix organization (GO:0030198) processes (Figure 4 D), which are relevant for osteogenesis. This suggests that mixture G down-regulates genes that are important for early osteogenesis in MSCs.

**Figure 3:**
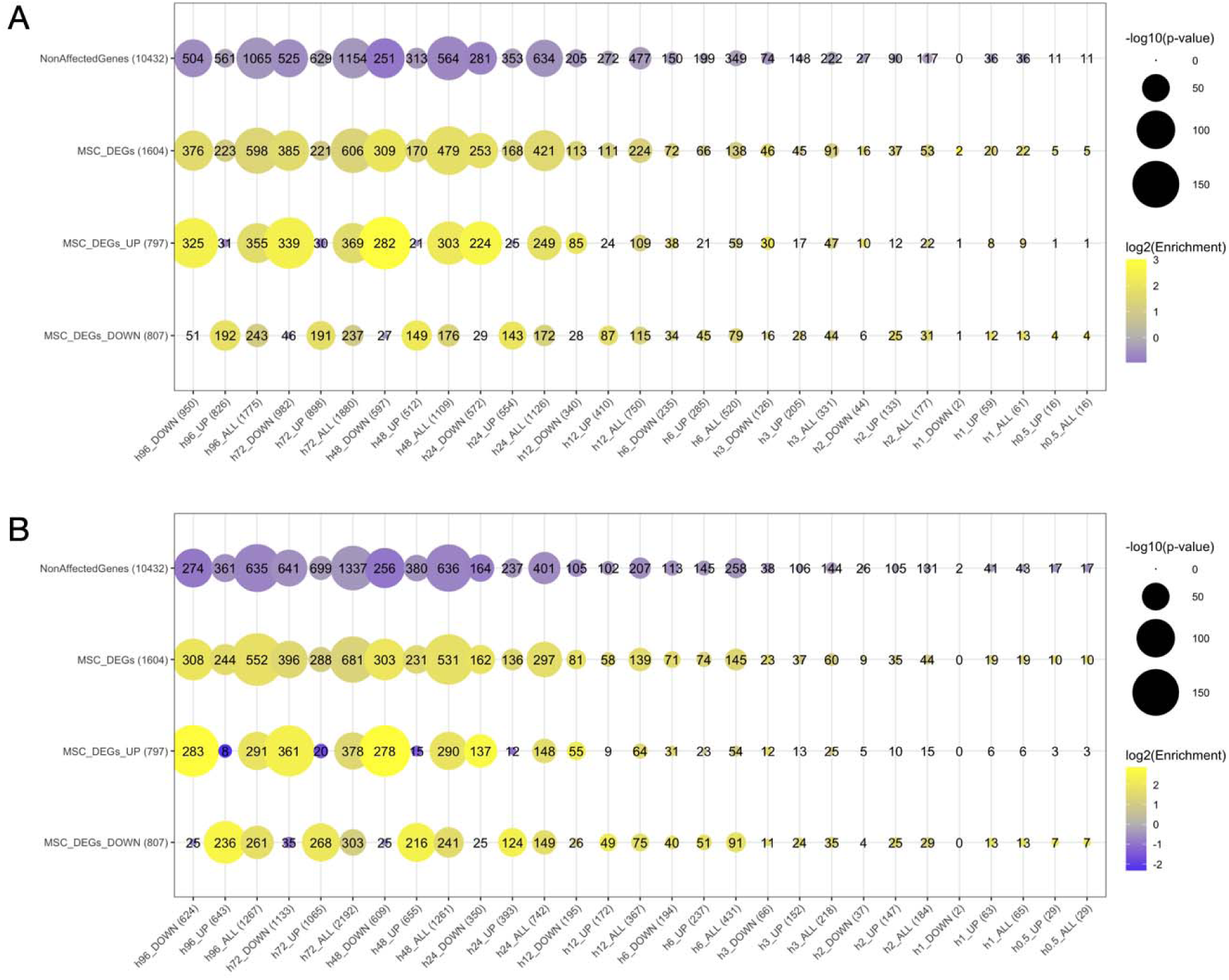
Overlap of mixture G DEGs with Early Adipogenesis/Osteogenesis DEGs. Overlaps, enrichment and significance between mixture G DEGs and DEGs identified at early time points of adipogenesis (A) or osteogenesis (B) induction in human bmMSCs from a previously published study by Peppel et al (2017). DEGs from this study at different time-points post induction of either adipogenesis or osteogenesis as well as direction of DEGs as either upregulated (“UP”) or downregulated (“DOWN”) are plotted on the x-axis while mixture G induced DEGs (all, UP or DOWN) as well as unaffected genes are shown on the y-axis. Ball size represents the p-value while the colour indicates log2-fold enrichment (positive, yellow indicates enrichment, negative, purple indicates depletion).

**Figure 4:**
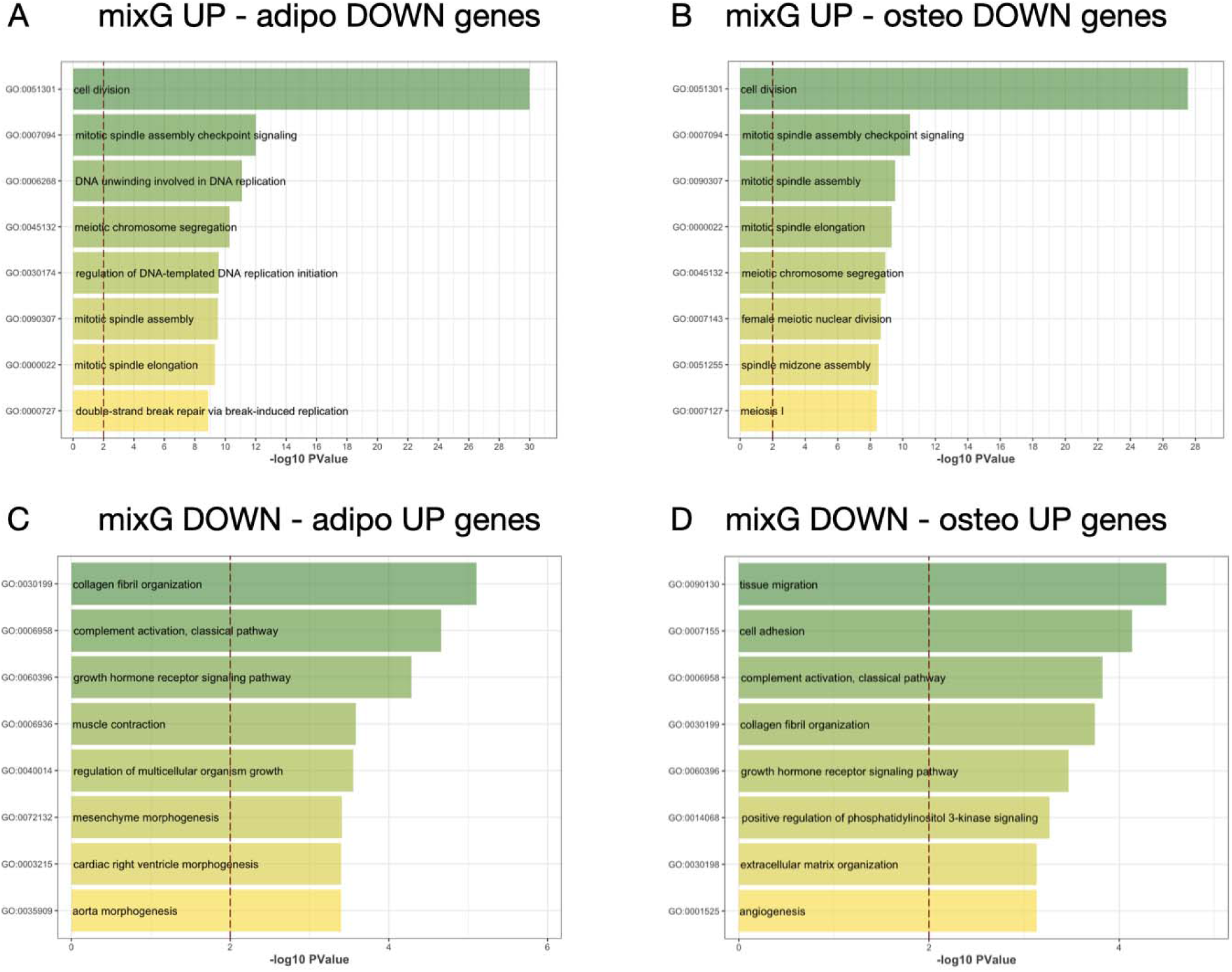
GO Enrichment Analysis for mixture G and Adipogenesis/Osteogenesis DEGs. Gene Ontology Enrichment analysis for genes overlapping between mixture G DEGs and DEGs identified at early time points of adipogenesis or osteogenesis induction in MSCs. A. For mixture G downregulated and adipogenesis upregulated DEGs. B. for mixture G upregulated and adipogenesis downregulated DEGs C. For mixture G downregulated and osteogenesis upregulated DEGs D. for mixture G upregulated and osteogenesis downregulated DEGs.

### 3.3 Mixture G dysregulates hormone response and disease-relevant genes

Considering that mixture G is composed of known and suspected EDCs, we investigated whether mixture G DEGs significantly overlap with genes that are known to be regulated b hormonal pathways of interest in the context of endocrine disruption and/or MSC differentiation. As shown in Figure 5, significant overlaps were observed for several of the selected pathways. Notably, however, mixture G DEGs did not overlap with genes regulated by PPARg, one of the master inducers of adipogenesis. On the other hand, they overlapped most prominently with corticoid-regulated genes, followed by estrogen-, progesterone-, and androgen-regulated genes, and to a lesser extent with thyroid-, and retinoic acid-regulated genes. While estrogen-regulated genes overlapped exclusively with upregulated DEGs, the others showed overlap with both up- and downregulated DEGs.

**Figure 5:**
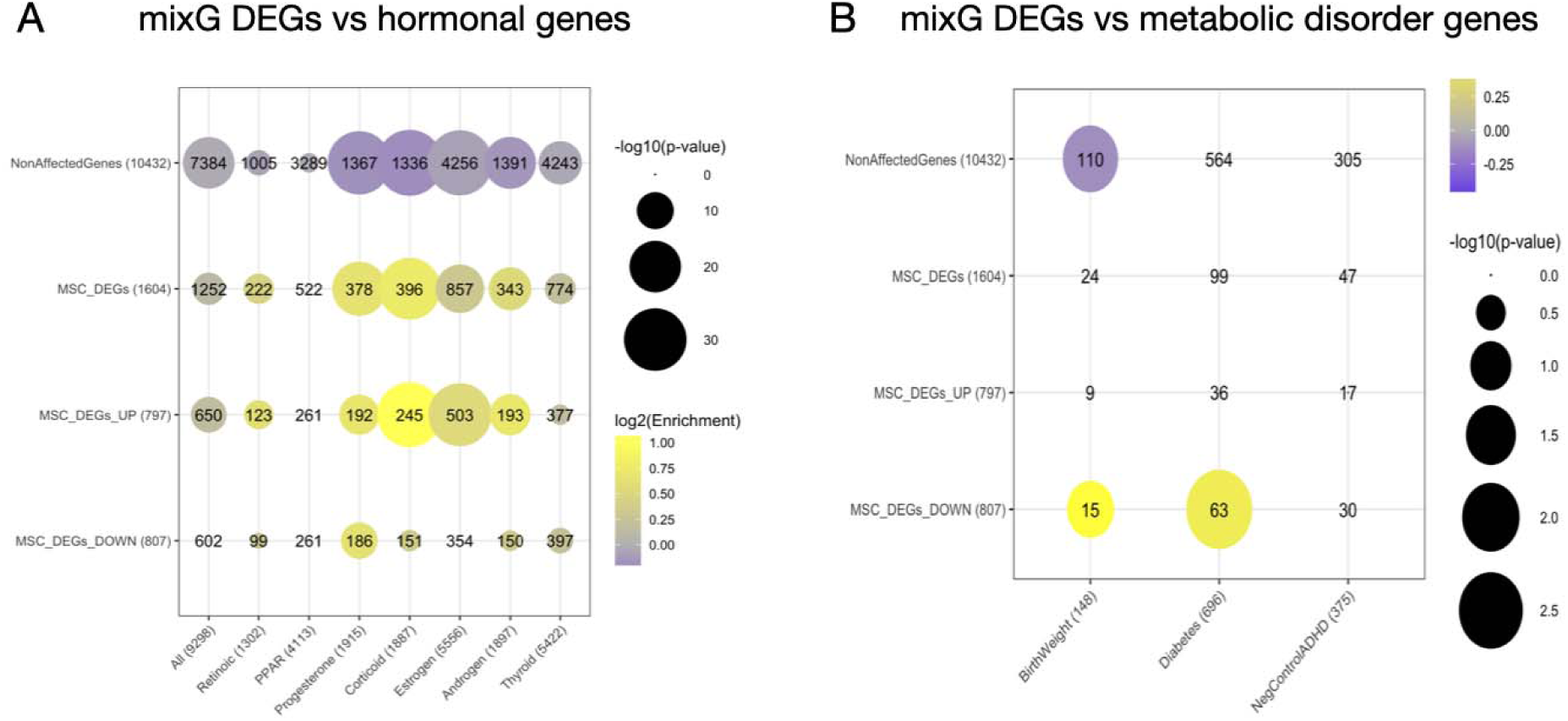
Overlap of the core set mixture G induced DEGs with genes regulated by major hormonal pathways and disease-relevant genes. Overlaps, enrichment and significance between mixture G DEGs and genes regulated by major hormonal pathways, retrieving the genes that are annotated in hormonal related pathways from the Human Molecular Signatures Database (MSigDB) (10.1016/j.cels.2015.12.004) (A) or disease-relevant genes, retrieving genes that are associated to birth weight and diabetes (and ADHD as negative control) by GWAS and present in the NHGRI-EBI GWAS Catalog ((Sollis et al., 2023); https://www.ebi.ac.uk/gwas/) (B).

Considering that mixture G was inferred by association with LBW, we assessed if Mix G-induced DEGs significantly overlap with genes carrying variants identified as risk factors for birth weight alterations and diabetes, due to the link between LBW and diabetes later in life. We found significant and specific overlaps between mixture G DEGs and genes associated with birth weight alterations and diabetes, harnessing the best-established database of genome wide association studies (GWAS) represented by the NHGRI-EBI GWAS Catalog ((Sollis et al., 2023); https://www.ebi.ac.uk/gwas/) (Fig. 5B). Notably, among the significant overlap between genes down-regulated by mixture G and genes associated with birth weight alterations, 14 out of 15 were associated with increased birth weight (Supplementary table S4). This is in line with the fact that higher mixture G exposure is associated with lower birth weight as it down-regulates genes that favor the opposite phenotype.

## 4. Discussion

In this study, we investigated the transcriptional effects underlying metabolic disruption by mixture G, an EDC mixture synthetized based on its association with LBW in epidemiological data. Numerous epidemiological studies have shown associations between LBW and childhood adiposity as well as other metabolic dysfunctions like insulin resistance in adulthood (Bianchi and Restrepo, 2022; Branda et al., 2022; de Mendonça et al., 2020; Hack et al., 1995; K. C. et al., 2020; LaFranchi, 2021; Nakano, 2020; Svensson et al., 2021; Tanis et al., 2005; Wikström et al., 2020). It has been hypothesized that these associations can, at least in part, be mediated by malfunctioning adipose tissue (Nakano, 2020). We found that mixture G, at human-relevant concentrations, induced significant transcriptional changes in hMSCs after 48 hrs of exposure and sustained adipogenesis after 21 days. Mixture G upregulated genes linked to cell proliferation, which is in contrast to what is observed when adipogenesis or osteogenesis in MSCs are induced by the respective adipogenesis or osteogenesis induction cell media. On the other hand, genes down-regulated by mixture G were associated with cell adhesion and extracellular matrix processes that are involved in early osteogenesis. Furthermore, mixture G-induced DEGs significantly overlapped with genes regulated by glucocorticoids and other endocrine pathways, but not by the master regulator of adipogenesis, PPARg. Finally, linking back the molecular signatures induced by mixture G to human phenotypes, we demonstrated a significant overlap of mixture G DEGs with genes associated with birth weight and diabetes.

Adipogenesis entails a highly regulated interplay of transcription factors, signaling pathways, and epigenetic modifications that drive differentiation from MSCs through preadipocyte progenitor cells and into adipocytes (de Sá et al., 2017). It is a critical process during the gestational period where adipose tissue development occurs (Orsso et al., 2020). However, MSCs can differentiate into a variety of cells of the mesodermal lineage besides, including osteoblasts and chondrocytes. This range of lineage commitments can be affected by chemical, physical, and biological factors (Breitfeld et al., 2020; de Sá et al., 2017; Han et al., 2019; Robert et al., 2020b; Sarjeant and Stephens, 2012), and a disturbed balance between, e.g., adipocytes and osteoblasts has been linked to several pathophysiologic conditions, such as obesity and osteoporosis (Armitage et al., 2008; Breitfeld et al., 2020; Cai et al., 2021; Nissen-Meyer et al., 2007; Teixeira et al., 2010; Z et al., 2018). Both bmMSCs and iPSC-derived MSCs are physiopathologically relevant *in vitro* models for studying the molecular regulation of adipogenesis (Augello and De Bari, 2010). In this setting, adipogenesis is induced by applying e.g. a mixture of insulin, dexamethasone (DEX) and 3-Isobutyl-1-methylxanthine (IBMX) (Scott et al., 2011). However, in the present study, all exposures with mixture G were conducted in hormone free media to avoid any forced induction of adipogenesis (or other lineages) and thereby capture direct exposure effects, including the most subtle. Consistently, while mixture G’s effects were not as strong as with the adipogenesis induction medium containing insulin, DEX and IBMX, Mixture G was by itself potent enough to significantly increase lipid droplet formation after 14-21 days exposure already at the 1X concentration. This is in accordance with our previous study where we reported significantly increased lipid accumulation in human bmMSCs upon exposure to a similar EDCs mixture, mixture G1, that consists of 14 compounds of which 8 were also present in mixture G (Lizunkova et al., 2022). Our findings are also consistent with those of Mentor et al. (2020), where developmental exposure to mixture G induced significant lipid accumulation in zebrafish (Mentor et al., 2020). A number of previously published studies have shown adipogenic properties of single EDCs included in mixture G, although at several fold higher concentrations than those in mixture G. For example, 100 µM of MBzP, 2.5 µM of PFOS, and 0.25 µM of PFHxS were found to induce adipogenesis in 3T3[L1 cell model (Hurst and Waxman, 2003b; Modaresi et al., 2022). In a bone marrow stromal cell model, 1 µM of MEHP induced adipogenesis and 10 µM of MEHP decreased osteoblast differentiation (Chiu et al., 2018). Notably, however, the effects of mixture G in this study were observed in the nano molar range, pointing to the enhanced potency of the mixture compared to the single chemicals.

For the transcriptomics analysis, we employed bmMSCs from two different human donors, as well as two iPSCs derived MSCs cell lines in order to enhance the robustness of our findings and reduce the potential biases associated with using a specific model or donor. Using these models, we identified a core set of genes that were differentially expressed in all cell models and followed the same dose-response curve. The core set served as a basis for identifying the early transcriptional mixture G-induced changes that could act as drivers for the observed adipogenic effect. The upregulated DEGs in the core set showed enrichment in GO terms related to cell division and DNA replication, suggesting that mixture G increases MSCs proliferation. Downregulated DEGs showed enrichment in the GO terms related to cell adhesion and extracellular matrix organization, the same GO terms enriched by genes upregulated upon induction of osteogenesis. Previous studies have explored the relationship between osteogenesis and cell adhesion indicating that changes in cytoskeletal tension can direct osteogenesis of MSCs and osteoblasts (Kaivosoja et al., 2013; Zhao et al., 2021). Interestingly, comparison of upregulated DEGs by mixture G with previously reported early transcriptional changes in MSCs (Peppel et al., 2017) did not significantly overlap with genes induced during adipogenesis. Additionally, no significant overlap was found between the mixture G DEGs and genes by the PPAR gamma, the master regulator of adipogenesis. It is important to note that in these studies adipogenesis was induced via supplementation with IBMX, DEX and indomethacin, while, as mentioned above, in the present study, MSCs were cultured with mixture G in hormone-free media. This difference suggests that mixture G does not initially act via induction of genes canonically recruited in adipogenic differentiation but rather by repressing genes that are physiologically involved in osteogenesis.

Previous studies have reported that during early MSCs differentiation, both toward the adipogenic and the osteogenic lineage, cell proliferation related processes are downregulated (de Sá et al., 2017; Peppel et al., 2017). Specifically during adipogenesis, some cell cycle genes were found to be downregulated as early as 24 h post induction of differentiation and the number of proliferating cells first increased up until 24 h and then decreased and stayed at almost zero from 48 h onwards (Marquez et al., 2017). However, when cellular proliferation was induced by addition of basic fibroblast growth factor during the first 48 h of adipogenic differentiation induction, the total number of adipocytes was enhanced (Marquez et al., 2017). Considering that mixture G induced cell-cycle- and proliferation associated genes, it is possible that it acts similarly, namely by increasing the total number of cells that become preadipocytes, which could result in an increased number of adipocytes after the 21 days exposure. Notably, treatment of human MSCs with 17β-estradiol induced proliferation in several studies (DiSilvio et al., 2006; Hong et al., 2011), which is interesting as mixture G upregulated DEGs significantly overlapped with estrogen-regulated genes.

In addition, mixture G DEGs significantly overlapped with genes regulated by glucocorticoids (GCs). In previous studies, GCs have been shown to affect the fate decision balance between adipogenesis and osteogenesis of MSCs, whereby at physiological concentrations, GCs were shown to promote osteogenesis, while excessive or pharmacological concentrations inhibit osteogenesis (Han et al., 2019). Several key pathways and genes were identified to be important for the fate decision of MSCs via the glucocorticoid receptor (GR) (Han et al., 2019). Notably, some of them were also identified in the overlap between mixture G DEGs and corticoid regulated genes, namely *FOXO1, SOX4, COL1A1*, and *TGFBR2*. The gene *FOXO1*, which was downregulated by mixture G, encodes FoxO1, a transcription factor that plays an important role in early osteogenesis (Teixeira et al., 2010) and has been shown to promote early osteogenic differentiation of MSCs (Ma et al., 2020). *SOX4* was also identified as a DEG downregulated by mixture G. SOX4 is highly expressed in osteoblast progenitor cells, and knockdown of *Sox4* in primary osteoblasts reduced proliferation of progenitor cells and delayed osteoblast differentiation (Nissen-Meyer et al., 2007, p. 4; Yi et al., 2021). *COL1A1,* the type 1 collagen gene, was also downregulated by mixture G. Collagen is accumulating during bone mineralization and mutations in this gene result in brittle bone disease Osteogenesis Imperfecta (Kaneto et al., 2014; Millington-Ward et al., 2004). Finally, *TGFBR2,* transforming growth factor beta receptor 2, was upregulated by mixture G. TGFBR2 is a transmembrane protein that binds TGF-β, suggesting that its upregulation leads to induction of TGF-β signaling (Nakamura et al., 2020). The TGFβ/bone morphogenic protein signaling pathway was shown to have a role in differentiation of MSCs both in the osteogenic lineage, by regulating the expression of the runt-related gene 2, and in the adipogenic lineage, by regulating the expression of PPARγ, (Han et al., 2019). GCs given during osteogenic differentiation were found to upregulate microRNAs (MiRs) targeting *FOXO1, SOX4* and *COL1A1* and downregulate MiRs targeting *TGFBR2* (Han et al., 2019), which suggests a similar effect of GC treatment on the expression of these genes as we have observed upon mixture G exposure.

Finally, we tested if genes altered by mixture G were significantly overlapping with genes associated with relevant phenotypes in humans by genome wide association studies, and found a significant overlap between genes down-regulated by mixture G and those associated with birth weight or diabetes. Notably, four genes were overlapping with both birth weight and diabetes GWAS, namely Insulin Receptor (INSR), Insulin Like Growth Factor 2 (IGF2), Insulin Like Growth Factor 1 Receptor (IGF1R), and Archaelysin Family Metallopeptidase 1 (AMZ1). IGF1 and insulin signalling play crucial roles in glucose response and function of adipose tissue, hence our findings suggest impaired functionality of adipocytes derived from mixture G-exposed MSCs. Interestingly, while both INSR and IGF1R are known to promote adipogenesis (Boucher, et al., 2016), we found that mixture G, while increasing differentiation of MSCs to pre-adipocytes, downregulates *INSR* and *IGF1R*. This may reflect the fact that, in human in vitro models of adipogenesis, insulin signalling appears involved only in the late phases rather than in the early stages we have investigated (Cignarelli et al., 2016). Additionally, IGF2 has also been shown to act as an inducer of adipogenesis in human models. Interestingly, previous study suggested that it promotes adipogenesis in subcutaneous pre-adipocytes while preventing it in visceral adipocytes, thus suggesting a protective role for IGF2 in body fat composition (Alfares et al., 2018). Whether the early down-regulation of *IGF2* by mixture G in our model would affect body fat composition will have to be addressed in future studies. The role of AMZ1 in regulating adipose tissue development or function is not established. Interestingly, however, it was found to be transcriptionally repressed in differentiating human primary pre-adipocytes by the EDC bisphenol A (BPA), which also induced adipogenesis via its estrogenic effect (Boucher et al., 2016, 2014). BPA was not included in mixture G, yet, a similar effect of mixture G could be at play, as it affected estrogen-regulated genes. Other mixture G-downregulated DEGs associated with increased birth weight have also been shown to be linked to metabolic disorders, albeit not found in the diabetes-related GWAS. One example is *RABGAP1* for which lower expression has been linked to obesity and diabetes type II (Zeng et al., 2021). Another one is Cdk5 and Abl enzyme substrate 1 (CABLES1), a cell cycle regulator, whose expression has been shown to be decreased in subcutaneous adipose tissue in persons with obesity and type 2 diabetes (Hetty et al., 2023).

## 5. Conclusion

In summary, here we show that an EDC mixture associated with lower birth weight biases MSC fate towards increased adipogenesis at human-relevant concentrations and alters transcription of genes related to osteogenesis, birth weight alterations and diabetes. This adds to the evidence that EDC mixtures are physiopathologically relevant as metabolic disruptors, providing novel insights into the underlying mechanisms.

## Supporting information

Supplementary Notebooks Bioinformatic Analysis

## Author contributions

CC: Formal analysis, Writing – Review and Editing; CG: Writing – Review and Editing; CGB: Writing – Review and Editing, Supervision, Project administration, Funding acquisition; CL: Resources; EE: Conceptualisation, Formal analysis, Investigation, Writing – Review and Editing, Visualisation; Supervision; JR: Conceptualisation, Writing – original draft, Supervision, Project administration, Funding acquisition; GB: Investigation; GT: Conceptualisation, Writing – Review and Editing, Supervision, Funding acquisition; NC: Conceptualisation, Investigation, Formal analysis, Writing – original draft, Supervision, Visualisation; PL: Investigation, Formal analysis, Writing – original draft, Writing – Review and Editing, Visualisation; PLG: Formal analysis, Writing – Review and Editing.

## Supplementary figures

**Table S1:** Individual chemicals and their concentrations within mixture G.

| Chemical Class | Chemical measured | Chemical name | Concentration 1 X Mix G (nM) |
| --- | --- | --- | --- |
| Phthalates | MEP | Monoethyl phthalate | 27,6 |
|  | MBP | Monobutyl phthalate | 23,1 |
|  | MBzP | Monobenzyl phthalate | 10,7 |
|  | MEHP | Mono-(2-ethylhexyl) phthalate | 14,6 |
|  | MINP | Di-isononyl phthalate | 21,1 |
| Antibacterial | Triclosan |  | 3,1 |
| PFAS | PFOA | Perfluorooctanoic acid | 3,4 |
|  | PFOS | Perfluorooctane sulfonate | 10,5 |
|  | PFHxS | Perfluorohexanesulphonic acid | 3,3 |

**Table S2:** Differentially expressed genes induced by mixture G in bmMSCs and iPSCs-MSCs. See excel file named Supplementary Table S2.

**Table S3:** Gene Ontology over-representation analyses on the differentially expressed genes. See excel file named Supplementary Table S3.

**Table S4:** Overlap of Mix G downregulated genes and risk genes associated with birth weight from the GWAS catalog (Sollis et al., 2023). See excel file named Supplementary Table S4.

**Figure S1:**
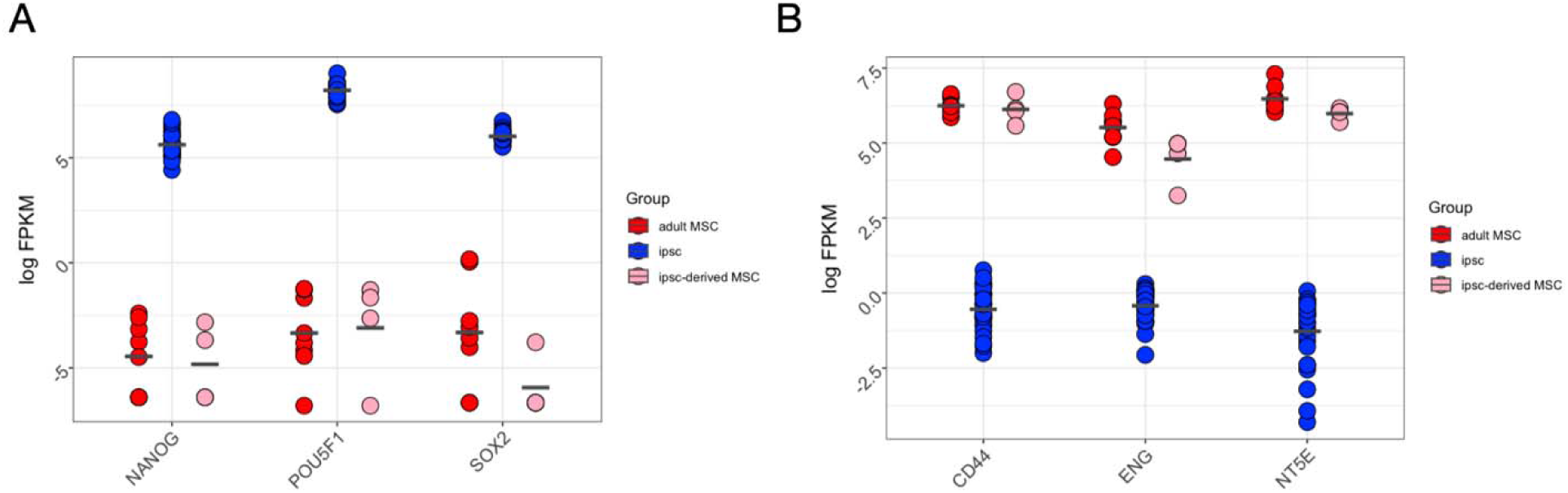
Validation of the MSC cellular systems. Gene expression analysis based on the log FPKM (fragments per kilobase of transcript per million mapped reads) values of MSCs models and iPSCs for markers of pluripotency (**A**), and markers of MSC (**B**).

**Figure S2:**
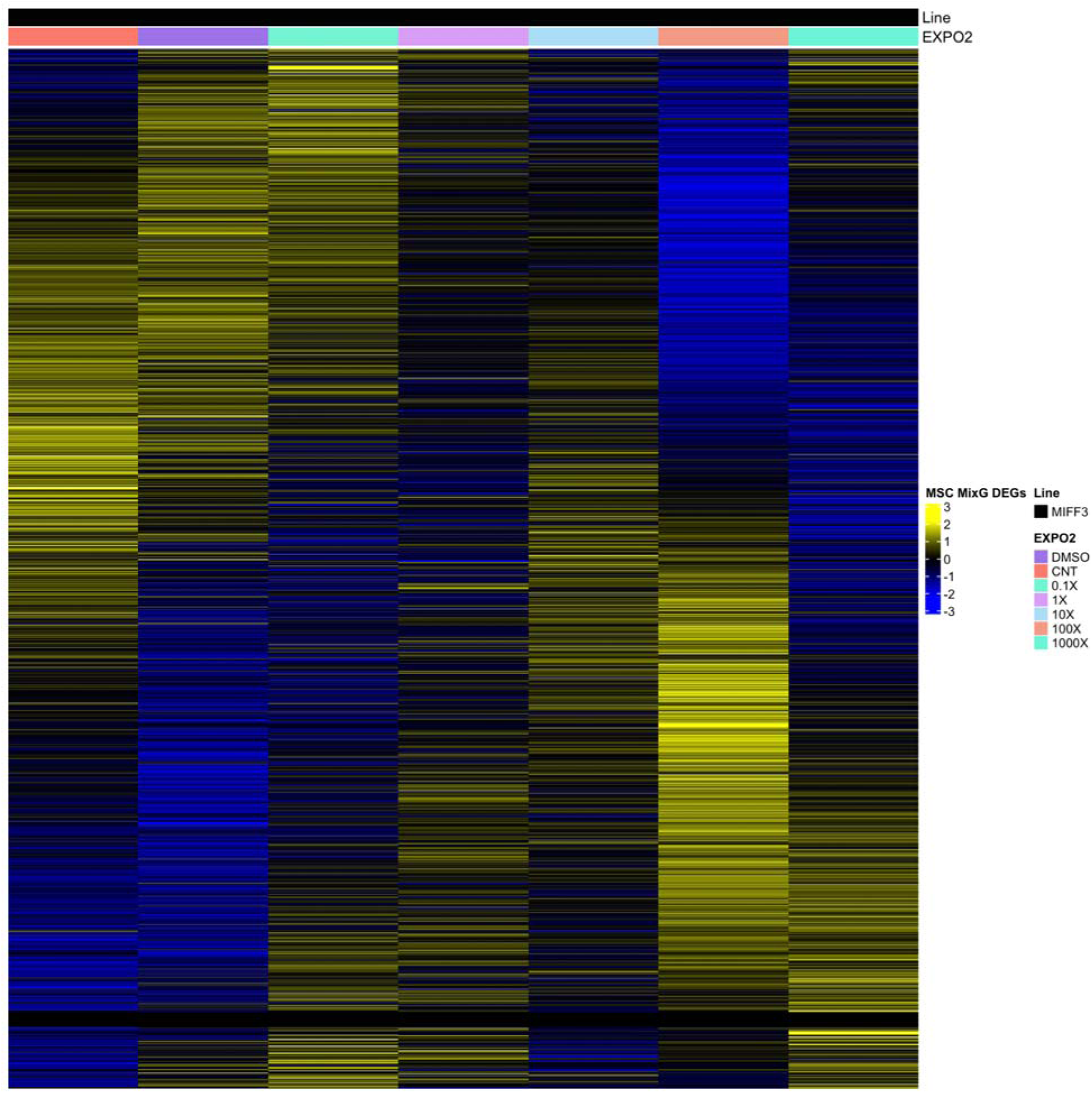
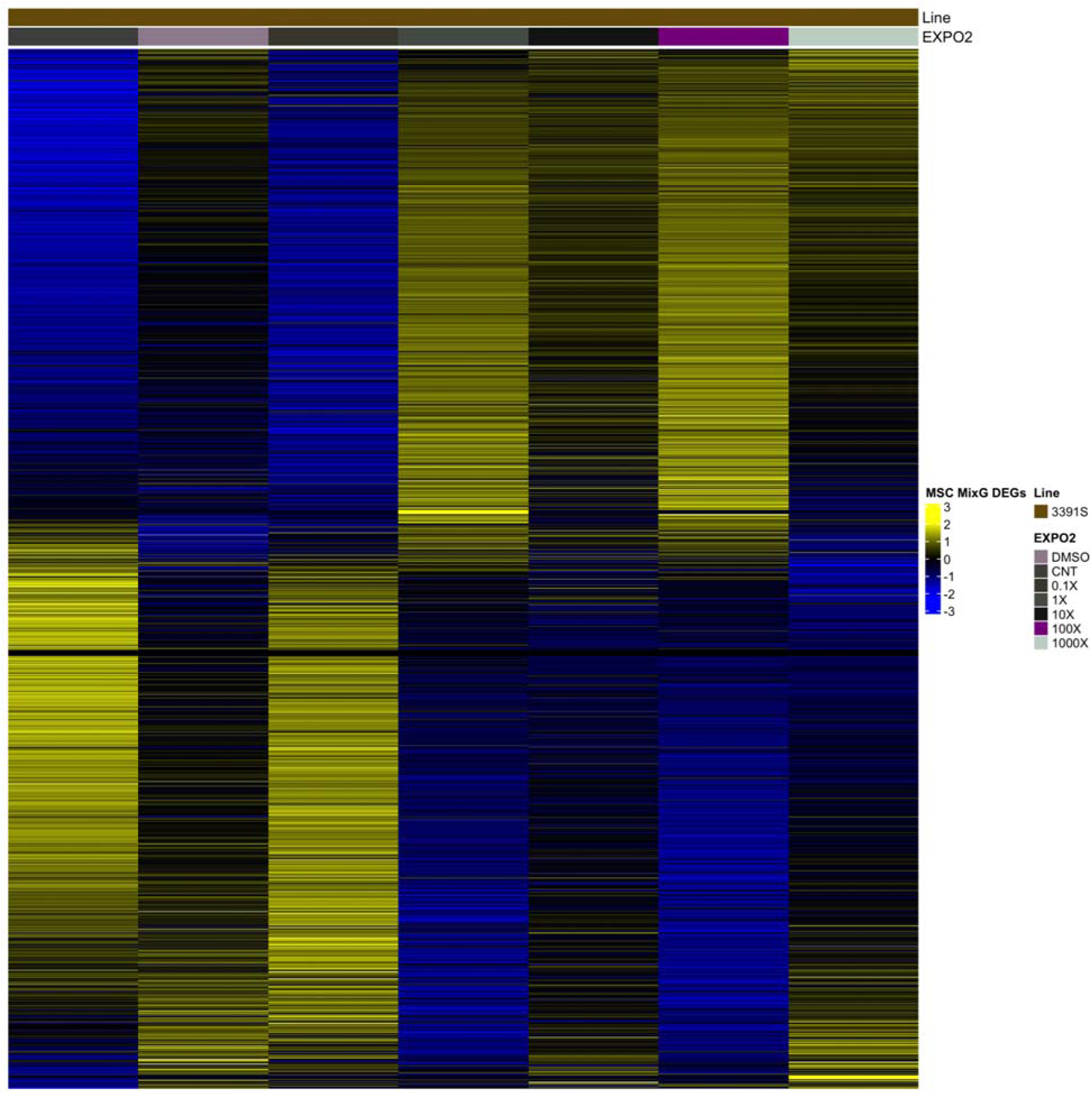
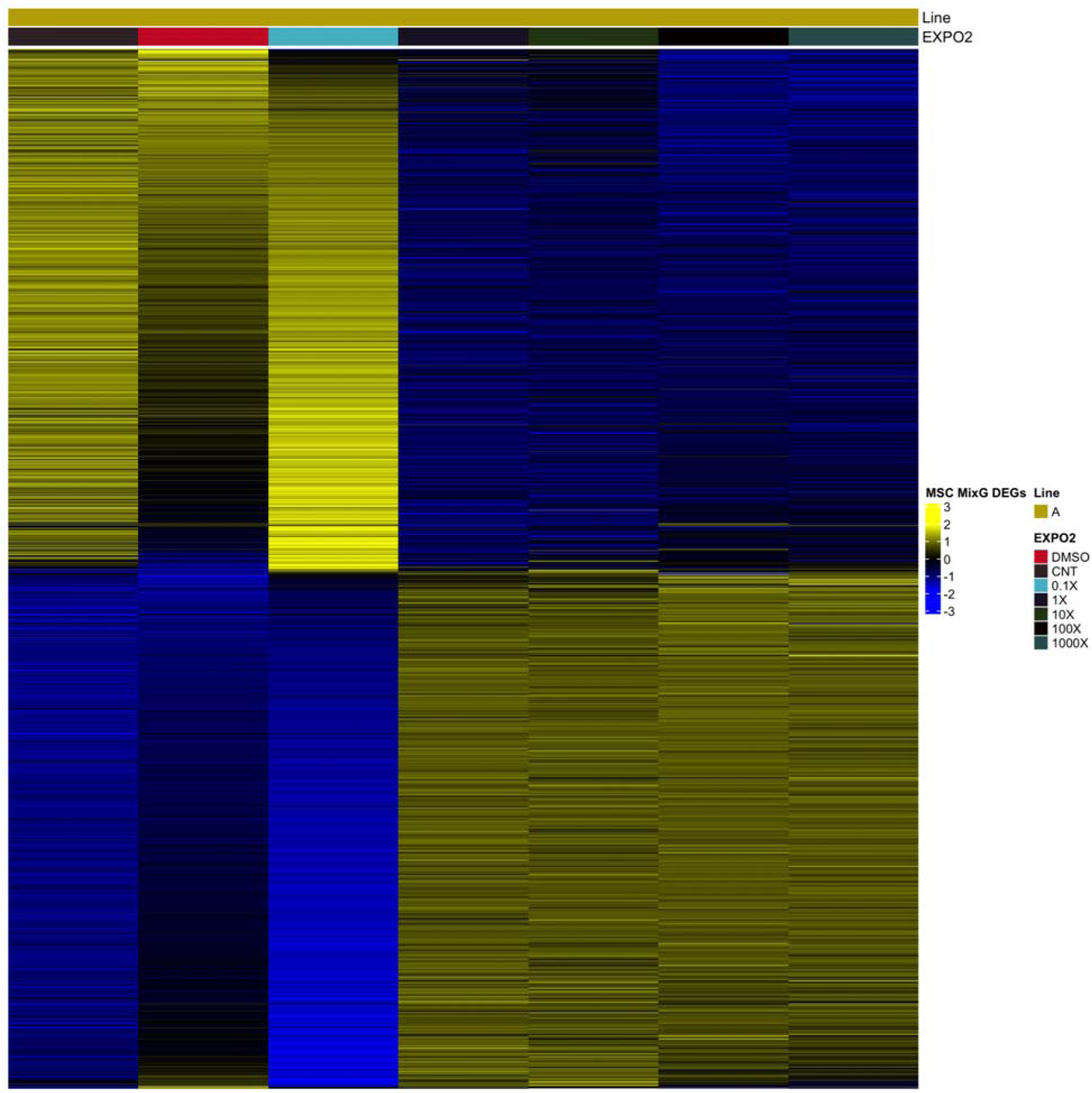
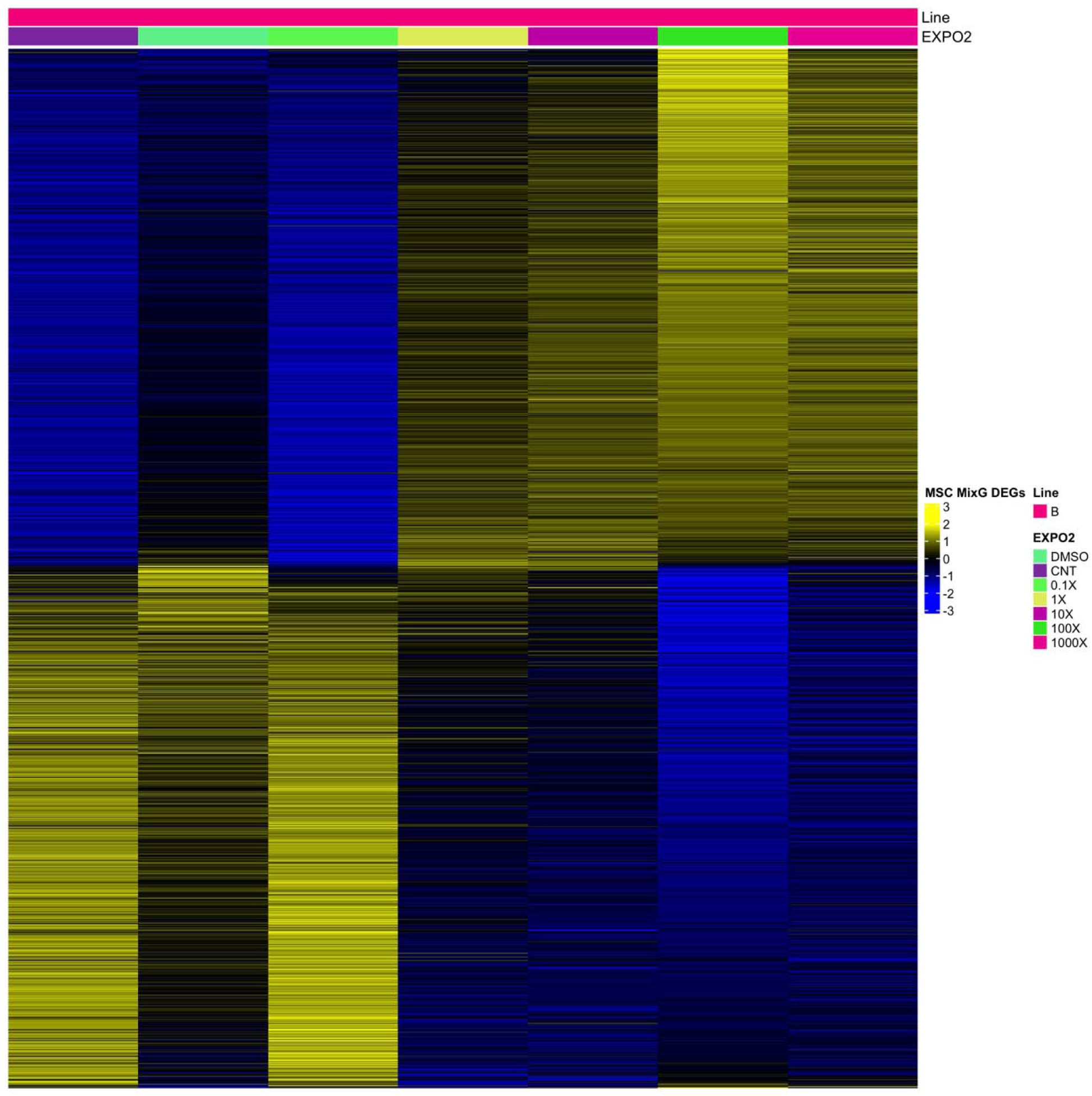
A heatmap showing the core set of 1604 differentially expressed genes (DEGs; FDR < 0.05, logFC > 0.2, logCPM > 0; both MSC models combined), showing different DEGs expression enrichment upon mixture G treatment in comparison to the control DMSO treatment. In order of appearance from the top: iPSC derived MSC Line1 Male; iPSC derived MSC Line2 Female; adult derived MSC Line1; adult derived MSC Line2.

**Figure S3:**
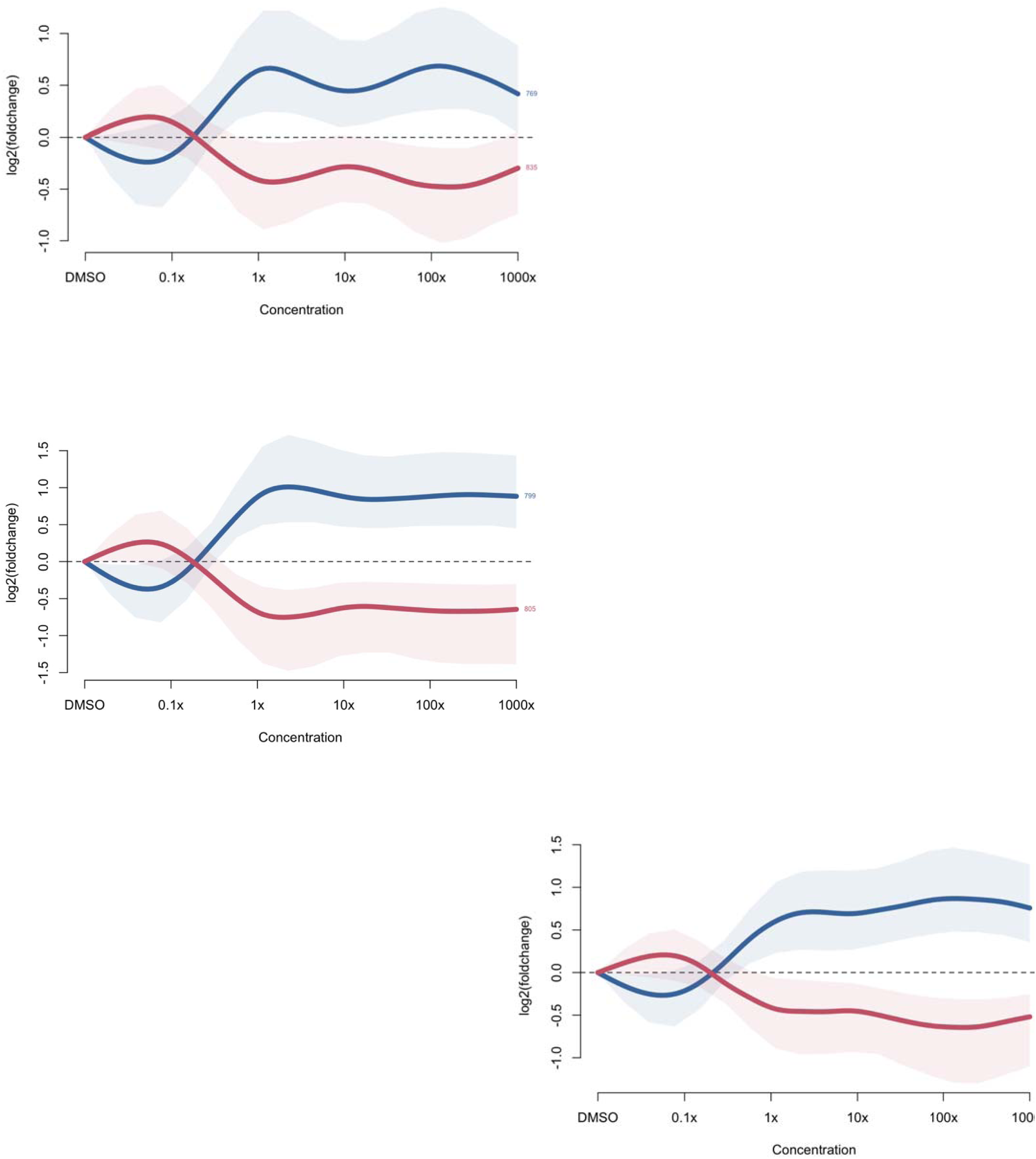
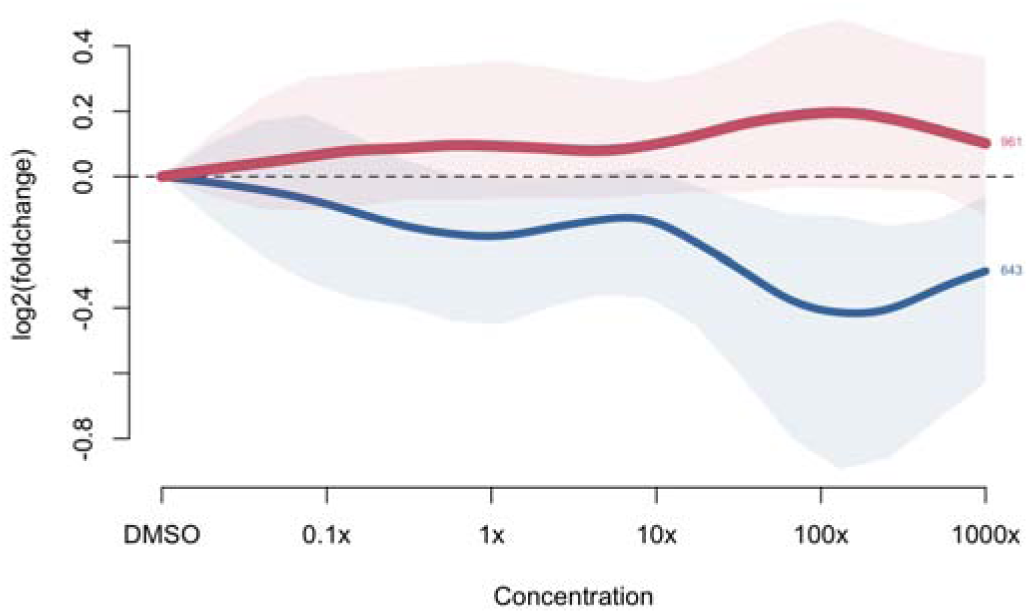
Dose response curve for each MSC cell line. In order of appearance from the top: DEGS dose-response patterns iPSC derived MSC Line1 Male; DEGS dose-response patterns iPSC derived MSC Line2 Female; DEGS dose-response patterns adult derived MSC Line1; DEGS dose-response patterns adult derived MSC Line2.

