## Supplementary Notebooks Bioinformatic Analysis for "A mixture of endocrine disrupting chemicals linked to lower birth weight induces adipogenesis and transcriptional changes related to birth weight alterations and diabetes": 01_MixG_MesenchimalStemCells.html

MixG Exposure on Mesenchymal Stem Cells


Code 

- Show All Code
- Hide All Code

### MixG Exposure on Mesenchymal Stem Cells

##### Data loading

```
suppressPackageStartupMessages({
  library(SummarizedExperiment)  
  library(SEtools)  
  library(edgeR)
  library(DT)
  library(pheatmap)
  library(plotly)
  library(dplyr)
  library(sva)
  library(overlapper)
  library(sechm)
})
source("Functions/EDC_Functions.R")
source("Functions/CriFormatted.R")
load("Data/AllSEcorrected_MSC.RData",verbose=T)
```

```
## Loading objects:
##   SEs
##   DEAs
```

```
load("Data/TotalSumExp.RData",verbose=T)
```

```
## Loading objects:
##   Total
```

```
load("Data/iPSC.RData", verbose = T)
```

```
## Loading objects:
##   e
```

```
load("Data/geneLengths.RData")
```

##### Validation of the MSC cellular systems

```
MSC <- as.data.frame(assays(Total[,which((Total$EXPO=="CNT"|Total$EXPO=="DMSO") & Total$type=="MSC")])$counts)
iPSC <- e
Tot <- merge(MSC,iPSC,by="row.names")
rownames(Tot) <- Tot$Row.names
Tot$Row.names <- NULL

TempDGE <- DGEList(counts=Tot,group=colnames(Tot))
TempDGE <- calcNormFactors(TempDGE, method='TMM')

fpkm <- edgeR::rpkm(TempDGE, gene.length=geneLengths[rownames(TempDGE)], normalized.lib.size=TRUE, prior.count=1, log=TRUE)

sampletypes <- rep(c("adult MSC","ipsc-derived MSC","adult MSC","ipsc"), c(4,4,4,ncol(iPSC)))
SampleColors <- c("adult MSC"="red", "ipsc-derived MSC"="pink", "ipsc"="blue")
GeneSet <- c("SOX2", "POU5F1", "NANOG")
for (i in 1:length(GeneSet))
  {
    Gene <- GeneSet[i]
    GeneCorrectedExp <- fpkm[Gene, ] 
    DataTemp <- data.frame(Sample=names(GeneCorrectedExp), Exp=as.numeric(GeneCorrectedExp), Group=sampletypes,        Gene=Gene)
    
    if(i==1){
      Data <- DataTemp
    }else{
      Data <- rbind(Data, DataTemp)
    }
  } 

    
plot <- ggplot(data=Data, aes(Gene,Exp, fill=Group))+
    geom_jitter(position=position_dodge(0.6), size=5, pch=21) + 
    stat_summary(fun.y=mean, fun.ymin=mean, fun.ymax=mean,
                 geom="crossbar", width=0.5, col='gray35', position=position_dodge(0.6)) + 
    scale_fill_manual(values=SampleColors) +
    theme_bw() + xlab('') + ylab("log FPKM") +
    theme(plot.title = element_text(face='bold', colour='darkred', size=18, hjust=0.5), 
          axis.title=element_text(size=14), axis.text=element_text(size=12.5, angle=45, hjust=1))
```

```
## Warning: The `fun.y` argument of `stat_summary()` is deprecated as of ggplot2 3.3.0.
## ℹ Please use the `fun` argument instead.
## This warning is displayed once every 8 hours.
## Call `lifecycle::last_lifecycle_warnings()` to see where this warning was
## generated.
```

```
## Warning: The `fun.ymin` argument of `stat_summary()` is deprecated as of ggplot2 3.3.0.
## ℹ Please use the `fun.min` argument instead.
## This warning is displayed once every 8 hours.
## Call `lifecycle::last_lifecycle_warnings()` to see where this warning was
## generated.
```

```
## Warning: The `fun.ymax` argument of `stat_summary()` is deprecated as of ggplot2 3.3.0.
## ℹ Please use the `fun.max` argument instead.
## This warning is displayed once every 8 hours.
## Call `lifecycle::last_lifecycle_warnings()` to see where this warning was
## generated.
```

```
plot
```

```
GeneSet <- c("ENG" , "NT5E" ,  "CD44")
for (i in 1:length(GeneSet))
  {
    Gene <- GeneSet[i]
    GeneCorrectedExp <- fpkm[Gene, ] 
    DataTemp <- data.frame(Sample=names(GeneCorrectedExp), Exp=as.numeric(GeneCorrectedExp), Group=sampletypes,        Gene=Gene)
    
    if(i==1){
      Data <- DataTemp
    }else{
      Data <- rbind(Data, DataTemp)
    }
  } 

    
plot <- ggplot(data=Data, aes(Gene,Exp, fill=Group))+
    geom_jitter(position=position_dodge(0.6), size=5, pch=21) + 
    stat_summary(fun.y=mean, fun.ymin=mean, fun.ymax=mean,
                 geom="crossbar", width=0.5, col='gray35', position=position_dodge(0.6)) + 
    scale_fill_manual(values=SampleColors) +
    theme_bw() + xlab('') + ylab("log FPKM") +
    theme(plot.title = element_text(face='bold', colour='darkred', size=18, hjust=0.5), 
          axis.title=element_text(size=14), axis.text=element_text(size=12.5, angle=45, hjust=1)) 
  
plot
```

#### DEGs heatmaps

```
msc.degs <- row.names(DEAs$acute.msc)[which(DEAs$acute.msc$FDR < 0.05 & ((abs(DEAs$acute.msc$logFC.EXPO0.1X) > 0.5 | abs(DEAs$acute.msc$logFC.EXPO1X) > 0.5 | abs(DEAs$acute.msc$logFC.EXPO10X) > 0.5 | abs(DEAs$acute.msc$logFC.EXPO100X) > 0.5 | abs(DEAs$acute.msc$logFC.EXPO1000X) > 0.5)) & DEAs$acute.msc$logCPM > 0)]
save(msc.degs,file = "Data/DEGsMSC.RData")

MixG <- SEs$acute.msc[,which(SEs$acute.msc$EXPO=="CNT"|SEs$acute.msc$EXPO=="0.1X"| SEs$acute.msc$EXPO=="1X"| SEs$acute.msc$EXPO=="10X"| SEs$acute.msc$EXPO=="100X"| SEs$acute.msc$EXPO=="1000X")]
```

###### All

```
MixG$Line2 <- as.character(MixG$Line)
MixG$Line2[which(MixG$Line2=="A")] <- "Adult MSC male"
MixG$Line2[which(MixG$Line2=="B")] <- "Adult MSC female"
MixG$Line2[which(MixG$Line2=="MIFF3")] <- "iPSC derived MSC male"
MixG$Line2[which(MixG$Line2=="3391S")] <- "iPSC derived MSC female"
MixG <- SEtools::log2FC(MixG, fromAssay = "logcpm",controls = which(MixG$EXPO=="CNT"), by = "Line", isLog = T)
sechm(MixG,assayName = "log2FC",msc.degs, do.scale = T, show_rownames = F, top_annotation =  c("Line2","EXPO2"), name = "MSC MixG DEGs")
```

###### iPSC derived MSC Line1 Male

```
sechm(MixG[,which(MixG$Line == "MIFF3")], msc.degs, do.scale = T, show_rownames = F, top_annotation =  c("EXPO2"), name = "MSC MixG DEGs",cluster_rows = FALSE)
```

```
## Using assay 'log2FC'
```

###### iPSC derived MSC Line2 Female

```
sechm(MixG[,which(MixG$Line == "3391S")], msc.degs, do.scale = T, show_rownames = F, top_annotation =  c("EXPO2"), name = "MSC MixG DEGs",cluster_rows = FALSE)
```

```
## Using assay 'log2FC'
```

###### adult derived MSC Line1

```
sechm(MixG[,which(MixG$Line == "A")], msc.degs, do.scale = T, show_rownames = F, top_annotation =  c("EXPO2"), name = "MSC MixG DEGs", cluster_rows = FALSE)
```

```
## Using assay 'log2FC'
```

###### adult derived MSC Line2

```
sechm(MixG[,which(MixG$Line == "B")], msc.degs, do.scale = T, show_rownames = F, top_annotation =  c("EXPO2"), name = "MSC MixG DEGs",cluster_rows = FALSE)
```

```
## Using assay 'log2FC'
```

#### DEGS dose-response patterns All

```
Genes<- intersect(row.names(DEAs$acute.msc), msc.degs)

design1 <- as.data.frame(colData(MixG))
fcmat1 <- getFoldchangeMatrix(assays(MixG)$logcpm[Genes,], design1, is.log = TRUE)

# dose-response clusters of DEGs
cc <- getConsClust(fcmat1,2)
autoLayout(1)
labs <- c("DMSO","0.1x","1x","10x","100x","1000x")
plotGenesClusters(fcmat1, design1, cc, labels=labs,  showNumber=T,spar = 0)
```

\_\_\_

###### DEGS dose-response patterns iPSC derived MSC Line1 Male

```
Genes<- intersect(row.names(DEAs$acute.msc), msc.degs)

design1 <- as.data.frame(colData(MixG[,which(MixG$Line == "MIFF3")]))
fcmat1 <- getFoldchangeMatrix(assays(MixG[,which(MixG$Line == "MIFF3")])$logcpm[Genes,], design1, is.log = TRUE)

# dose-response clusters of DEGs
cc <- getConsClust(fcmat1,2)
autoLayout(1)
labs <- c("DMSO","0.1x","1x","10x","100x","1000x")
plotGenesClusters(fcmat1, design1, cc, labels=labs,  showNumber=T,spar = 0)
```

\_\_\_

###### DEGS dose-response patterns iPSC derived MSC Line2 Female

```
Genes<- intersect(row.names(DEAs$acute.msc), msc.degs)

design1 <- as.data.frame(colData(MixG[,which(MixG$Line == "3391S")]))
fcmat1 <- getFoldchangeMatrix(assays(MixG[,which(MixG$Line == "3391S")])$logcpm[Genes,], design1, is.log = TRUE)

# dose-response clusters of DEGs
cc <- getConsClust(fcmat1,2)
autoLayout(1)
labs <- c("DMSO","0.1x","1x","10x","100x","1000x")
plotGenesClusters(fcmat1, design1, cc, labels=labs,  showNumber=T,spar = 0)
```

---

###### DEGS dose-response patterns adult derived MSC Line1

```
Genes<- intersect(row.names(DEAs$acute.msc), msc.degs)

design1 <- as.data.frame(colData(MixG[,which(MixG$Line == "A")]))
fcmat1 <- getFoldchangeMatrix(assays(MixG[,which(MixG$Line == "A")])$logcpm[Genes,], design1, is.log = TRUE)

# dose-response clusters of DEGs
cc <- getConsClust(fcmat1,2)
autoLayout(1)
labs <- c("DMSO","0.1x","1x","10x","100x","1000x")
plotGenesClusters(fcmat1, design1, cc, labels=labs,  showNumber=T,spar = 0)
```

---

###### DEGS dose-response patterns adult derived MSC Line2

```
Genes<- intersect(row.names(DEAs$acute.msc), msc.degs)

design1 <- as.data.frame(colData(MixG[,which(MixG$Line == "B")]))
fcmat1 <- getFoldchangeMatrix(assays(MixG[,which(MixG$Line == "B")])$logcpm[Genes,], design1, is.log = TRUE)

# dose-response clusters of DEGs
cc <- getConsClust(fcmat1,2)
autoLayout(1)
labs <- c("DMSO","0.1x","1x","10x","100x","1000x")
plotGenesClusters(fcmat1, design1, cc, labels=labs,  showNumber=T,spar = 0)
```

##### Altered birth weight genes

```
load("Data/MetabGenes.RData", verbose = T)
## Loading objects:
##   metabolism
Genes<- intersect(row.names(DEAs$acute.msc), metabolism$birthWeight)
design1 <- as.data.frame(colData(MixG))

fcmat1 <- getFoldchangeMatrix(assays(MixG)$logcpm[Genes,], design1, is.log = TRUE)

cc<- c(1:length(Genes))
names(cc) <- Genes
autoLayout(1)
labs <- c("CNT","0.1x","1x","10x","100x","1000x")
plotGenesClusters(fcmat1, design1, cc, labels=labs,  showNumber=T,spar = 0)
```

##### Altered obesity DEGs

```
load("Data/MetabGenes.RData", verbose = T)
## Loading objects:
##   metabolism
msc.degs_strict <- row.names(DEAs$acute.msc)[which((abs(DEAs$acute.msc$logFC.EXPO0.1X) > 1 | abs(DEAs$acute.msc$logFC.EXPO1X) > 1 | abs(DEAs$acute.msc$logFC.EXPO10X) > 1 | abs(DEAs$acute.msc$logFC.EXPO100X) > 1 | abs(DEAs$acute.msc$logFC.EXPO1000X) > 1) & DEAs$acute.msc$FDR <= 0.01 & DEAs$acute.msc$logCPM > 0)]
Genes<- intersect(msc.degs_strict, metabolism$obesity)
fcmat1 <- getFoldchangeMatrix(assays(MixG)$logcpm[Genes,], design1, is.log = TRUE)

cc<- c(1:length(Genes))
names(cc) <- Genes
autoLayout(1)
labs <- c("CNT","0.1x","1x","10x","100x","1000x")
plotGenesClusters(fcmat1, design1, cc, labels=labs,  showNumber=T,spar = 0)
```

##### Adipogenesis and osteogenesis genes

```
a <- c("GPLD1","CEBPB","CEBPD","AGT","APOE","CHST11","CXCL12","FOXC1")
b <- c("IFRD1","SEMA3D","IGF2","IGFBP7","NRP1","SEMA3A","JAZF1","SEMA3B")
c <- c("KLF2","PIN1","POSTN","PSAP","IGFBP2","SIRT6","SOX9","BSCL2")
d <- c("SPP1","TRPV2","WNT5A","TCF7L2","LMNA","CAV1","IGF2BP2","EZH2","INSR")
e <- c("NRP1", "SEMA3B")

scc <- function(x){
    cc <- 1:length(x)
    names(cc) <- x
    return(cc)
}

Genes<- union(a,union(b,union(c,d)))

fcmat <- getFoldchangeMatrix(assays(MixG)$logcpm[Genes,], design1, is.log = TRUE)
#fcmat <- getFoldchangeMatrix(donorm(MSC_MixG.counts),MSC_MixG.design)

labs <- c("DMSO","0.1x","1x","10x","100x","1000x")
plotGenesClusters(fcmat[a,], design1, scc(a), labels=labs, showNumber=T, q=c(0.2,0.8))
```

```
plotGenesClusters(fcmat[b,], design1, scc(b), labels=labs, showNumber=T, q=c(0.2,0.8))
```

```
plotGenesClusters(fcmat[c,], design1, scc(c), labels=labs, showNumber=T, q=c(0.2,0.8))
```

```
plotGenesClusters(fcmat[d,], design1, scc(d), labels=labs, showNumber=T, q=c(0.2,0.8))
```

```
plotGenesClusters(fcmat[e,], design1, scc(e), labels=labs, showNumber=T, q=c(0.2,0.8))
```

#### Overlap Analysis

##### Metabolic Genes

```
load("Data/MetabGenes.RData", verbose = T)
## Loading objects:
##   metabolism
adipo <- c(c("GPLD1","CEBPB","CEBPD","AGT","APOE","CHST11","CXCL12","FOXC1"), c("IFRD1","SEMA3D","IGF2","IGFBP7","NRP1","SEMA3A","JAZF1","SEMA3B"), c("KLF2","PIN1","POSTN","PSAP","IGFBP2","SIRT6","SOX9","BSCL2"),c("SPP1","TRPV2","WNT5A","TCF7L2","LMNA","CAV1","IGF2BP2","EZH2","INSR"), c("NRP1", "SEMA3B"))
metabolism$adipo <- adipo


affected <- row.names(DEAs$acute.msc)[which(DEAs$acute.msc$FDR <= 0.05 & DEAs$acute.msc$logCPM > 0)]

controlNeg <- row.names(DEAs$acute.msc)[-which(DEAs$acute.msc$FDR <= 0.05 & DEAs$acute.msc$logCPM > 0)]


msc.degs <- row.names(DEAs$acute.msc)[which(DEAs$acute.msc$FDR < 0.05 & ((abs(DEAs$acute.msc$logFC.EXPO0.1X) > 0.5 | abs(DEAs$acute.msc$logFC.EXPO1X) > 0.5 | abs(DEAs$acute.msc$logFC.EXPO10X) > 0.5 | abs(DEAs$acute.msc$logFC.EXPO100X) > 0.5 | abs(DEAs$acute.msc$logFC.EXPO1000X) > 0.5)) & DEAs$acute.msc$logCPM > 0)]

#msc.degs_strict <- row.names(DEAs$acute.msc)[which((abs(DEAs$acute.msc$logFC.EXPO0.1X) > 1 | abs(DEAs$acute.msc$logFC.EXPO1X) > 1 | abs(DEAs$acute.msc$logFC.EXPO10X) > 1 | abs(DEAs$acute.msc$logFC.EXPO100X) > 1 | abs(DEAs$acute.msc$logFC.EXPO1000X) > 1) & DEAs$acute.msc$FDR <= 0.01 & DEAs$acute.msc$logCPM > 0)]

msc.degs_DOWN <- row.names(DEAs$acute.msc)[which(DEAs$acute.msc$FDR < 0.05 & ((abs(DEAs$acute.msc$logFC.EXPO0.1X) > 0.5 | abs(DEAs$acute.msc$logFC.EXPO1X) > 0.5 | abs(DEAs$acute.msc$logFC.EXPO10X) > 0.5 | abs(DEAs$acute.msc$logFC.EXPO100X) > 0.5 | abs(DEAs$acute.msc$logFC.EXPO1000X) > 0.5)) & DEAs$acute.msc$logCPM > 0 & (DEAs$acute.msc$logFC.EXPO1000X + DEAs$acute.msc$logFC.EXPO100X + DEAs$acute.msc$logFC.EXPO10X + DEAs$acute.msc$logFC.EXPO1X)/4 < 0)]

msc.degs_UP <- row.names(DEAs$acute.msc)[which(DEAs$acute.msc$FDR < 0.05 & ((abs(DEAs$acute.msc$logFC.EXPO0.1X) > 0.5 | abs(DEAs$acute.msc$logFC.EXPO1X) > 0.5 | abs(DEAs$acute.msc$logFC.EXPO10X) > 0.5 | abs(DEAs$acute.msc$logFC.EXPO100X) > 0.5 | abs(DEAs$acute.msc$logFC.EXPO1000X) > 0.5)) & DEAs$acute.msc$logCPM > 0 &  (DEAs$acute.msc$logFC.EXPO1000X + DEAs$acute.msc$logFC.EXPO100X + DEAs$acute.msc$logFC.EXPO10X + DEAs$acute.msc$logFC.EXPO1X)/4 > 0)]


MixG_DEGS <- list(MSC_DEGs_DOWN=msc.degs_DOWN, MSC_DEGs_UP=msc.degs_UP, MSC_DEGs=msc.degs, NonAffectedGenes=controlNeg)


m <- overlapper::multintersect(ll = MixG_DEGS, ll2 = metabolism, universe = rownames(filterGenes(SEs$acute.msc)))
dotplot.multintersect(m)
```

##### NDD genes (as a negative control)

```
load("Data/ASD.RData", verbose = T)
## Loading objects:
##   ASD
##   SFARI
##   SFARIgenes
##   AutismKB
##   DecipherNeuro
##   MSSNG
##   ASDsevere
##   ID
##   NeuroOmim
##   iPSYCH
##   GDI

m <- overlapper::multintersect(ll = MixG_DEGS, ll2 = ASD, universe = rownames(filterGenes(SEs$acute.msc)))
dotplot.multintersect(m)
```

##### Overlap DEGs and hormonal genes

```
load("Data/HormonalGenes/HormonalGenes.RData",verbose = T)
## Loading objects:
##   Thyroid
##   Androgen
##   Estrogen
##   Corticoid
##   Progesterone
##   PPAR
##   Retinoic

uni <-rownames(filterGenes(SEs$acute.msc))


hormonal <- list(Thyroid=Thyroid, Androgen=Androgen, Estrogen=Estrogen, Corticoid=Corticoid,Progesterone=Progesterone,PPAR=PPAR, Retinoic=Retinoic, All= union(Thyroid,union(Androgen, union(Estrogen,union(Corticoid, union(Progesterone,union(PPAR,Retinoic)))))))

m <- overlapper::multintersect(MixG_DEGS, hormonal, universe = uni)
overlapper::dotplot.multintersect(m)
```

##### Extract overlap between MixG DEGs and corticoids genes

```
overlap <- list("MixG_DEGS"=MixG_DEGS$MSC_DEGs, "Corticoids"=hormonal$Corticoid,"intersection"= intersect(MixG_DEGS$MSC_DEGs,hormonal$Corticoid))
save(overlap, file = "Data/Overlap_MixG_Corticoids.RData")

library(stringi)
## Warning: package 'stringi' was built under R version 4.3.1
test <- stri_list2matrix(overlap, byrow=FALSE)
colnames(test) <- c("MixG_DEGS", "Corticoids","Intersection")
test <- as.data.frame(test)
library(openxlsx)
write.xlsx(test, "Data/Overlap_MixG_Corticoids.xlsx")
```

##### Overlap DEGs and adipogenesis genes from https://doi.org/10.1016/j.stemcr.2017.02.018

```
# library(readxl)
# 
# for (x in c("h0.5","h1","h2","h3","h6","h12","h24","h48","h72","h96")) {
# 
# x <- read_excel(paste0("Data/2023_Data/AdipogenesisDEGS/",x) ,sheet = "Sheet 1")
# paste0(x,"_UP") <- x$...1[which(x$adj.P.Val<0.001 & x$logFC>0)]
# paste0(x,"_DOWN") <- x$...1[which(x$adj.P.Val<0.001& x$logFC<0)]
#   
# }
```

```
library(readxl)


for (x in c("h0.5","h1","h2","h3","h6","h12","h24","h48","h72","h96")) {

  
tmp <- read_excel(paste0(paste0("Data/2023_Data/AdipogenesisDEGS/",x),".xlsx") ,sheet = "Sheet 1")

tmp2<-tmp$Gene[which(tmp$adj.P.Val<0.001)]
tmp3<-tmp$Gene[which(tmp$adj.P.Val<0.001 & tmp$logFC>0)]
tmp4<-tmp$Gene[which(tmp$adj.P.Val<0.001 & tmp$logFC<0)]
assign(paste0(x,"_ALL"), tmp2)
assign(paste0(x,"_UP"), tmp3)
assign(paste0(x,"_DOWN"), tmp4)

}

adipo <- list()
adipo <- list(
"h0.5_ALL"=h0.5_ALL, "h0.5_UP"=h0.5_UP, "h0.5_DOWN"=h0.5_DOWN, 
"h1_ALL"=h1_ALL, "h1_UP"=h1_UP, "h1_DOWN"=h1_DOWN,
"h2_ALL"=h2_ALL, "h2_UP"=h2_UP, "h2_DOWN"=h2_DOWN,
"h3_ALL"=h3_ALL, "h3_UP"=h3_UP, "h3_DOWN"=h3_DOWN,
"h6_ALL"=h6_ALL, "h6_UP"=h6_UP, "h6_DOWN"=h6_DOWN,
"h12_ALL"=h12_ALL, "h12_UP"=h12_UP, "h12_DOWN"=h12_DOWN,
"h24_ALL"=h24_ALL, "h24_UP"=h24_UP, "h24_DOWN"=h24_DOWN,
"h48_ALL"=h48_ALL, "h48_UP"=h48_UP, "h48_DOWN"=h48_DOWN,
"h72_ALL"=h72_ALL, "h72_UP"=h72_UP, "h72_DOWN"=h72_DOWN,
"h96_ALL"=h96_ALL, "h96_UP"=h96_UP, "h96_DOWN"=h96_DOWN
)
```

```
# library(readxl)
# h3 <- read_excel("Data/Adipogenesis/DEG_0hr_vs_3hr.xlsx",sheet = "Sheet1")
# h3UP <- h3$Gene[which(h3$adj.P.Val<0.001 & h3$logFC>0)]
# h3DOWN <- h3$Gene[which(h3$adj.P.Val<0.001& h3$logFC<0)]

# h6 <- read_excel("Data/Adipogenesis/DEG_0hr_vs_6hr.xlsx",sheet = "Sheet1")
# h6 <- h6$Gene[which(h6$adj.P.Val<0.001)]
# 
# h24 <- read_excel("Data/Adipogenesis/DEG_0hr_vs_24hr.xlsx",sheet = "Sheet1")
# h24 <- h24$Gene[which(h24$adj.P.Val<0.001)]
# 
# h48 <- read_excel("Data/Adipogenesis/DEG_0hr_vs_48hr.xlsx",sheet = "Sheet1")
# h48 <- h48$Gene[which(h48$adj.P.Val<0.001)]
# 
# h96 <- read_excel("Data/Adipogenesis/DEG_0hr_vs_96hr.xlsx",sheet = "Sheet1")
# h96 <- h96$Gene[which(h96$adj.P.Val<0.001)]
# 
# adipo <- list(h6=h6,h24=h24)
```

###### only adipogenesis DEGs vs themselves

```
uni <-rownames(filterGenes(SEs$acute.msc))

m <- overlapper::multintersect(adipo, universe = uni)
overlapper::dotplot.multintersect(m)
## Warning: Removed 378 rows containing missing values (`geom_point()`).
## Warning: Removed 378 rows containing missing values (`geom_text()`).
```

###### MixG\_DEGS vs adipogenesis DEGs

```
m <- overlapper::multintersect(MixG_DEGS, adipo, universe = uni)
overlapper::dotplot.multintersect(m)
```

```
write.table(m$of, file = "Data/2023_Data/overlaps_adipo.txt", sep = "\t")
#write.csv(m$of, file = "Data/2023_Data/overlaps.csv")


#save overlapping genes for GO analysis
adipo_DOWN <- intersect(MixG_DEGS$MSC_DEGs_UP, adipo$h48_DOWN)
adipo_UP <- intersect(MixG_DEGS$MSC_DEGs_DOWN, adipo$h48_UP)

save(adipo_DOWN,adipo_UP, file = "Data/adipo.RData")
```

##### Heatmaps to check UP and DOWN regulated DEGs across lines

upregulated DEGs are upregulated in both adult MSC lines

```
sechm(MixG[,which(MixG$Line == "A"|MixG$Line == "B" )],MixG_DEGS$MSC_DEGs_UP, do.scale = T, show_rownames = F, top_annotation =  c("Line","EXPO2"), name = "MSC MixG DEGs")
```

```
## Using assay 'log2FC'
```

downregulated DEGs are downregulated in both adult MSC lines

```
sechm(MixG[,which(MixG$Line == "A"|MixG$Line == "B" )],MixG_DEGS$MSC_DEGs_DOWN, do.scale = T, show_rownames = F, top_annotation =  c("Line","EXPO2"), name = "MSC MixG DEGs")
```

```
## Using assay 'log2FC'
```

upregulated DEGs are upregulated in the female iPSC-MSC line, while
they are less clear cut in the male iPSC-MSC line

```
sechm(MixG[,which(MixG$Line == "3391S" )],MixG_DEGS$MSC_DEGs_UP, do.scale = T, show_rownames = F, top_annotation =  c("Line","EXPO2"), name = "Adipo DEGs")
```

```
## Using assay 'log2FC'
```

```
sechm(MixG[,which(MixG$Line == "MIFF3")],MixG_DEGS$MSC_DEGs_UP, do.scale = T, show_rownames = F, top_annotation =  c("Line","EXPO2"), name = "Adipo DEGs")
```

```
## Using assay 'log2FC'
```

downregulated DEGs are downregulated in the female iPSC-MSC line,
while they are less clear cut in the male iPSC-MSC line

```
sechm(MixG[,which(MixG$Line == "3391S" )],MixG_DEGS$MSC_DEGs_DOWN, do.scale = T, show_rownames = F, top_annotation =  c("Line","EXPO2"), name = "Adipo DEGs")
```

```
## Using assay 'log2FC'
```

```
sechm(MixG[,which(MixG$Line == "MIFF3")],MixG_DEGS$MSC_DEGs_DOWN, do.scale = T, show_rownames = F, top_annotation =  c("Line","EXPO2"), name = "Adipo DEGs")
```

```
## Using assay 'log2FC'
```

###### heaatmap to check adipogenesis genes in our dataset

if we check how upregulated genes during adipogenesis behave in adult
MSC lines we see that the majority are downregulated, but some are also
upregulated

```
sechm(MixG[,which(MixG$Line == "A"|MixG$Line == "B" )],adipo$h96_UP, do.scale = T, show_rownames = F, top_annotation =  c("Line","EXPO2"), name = "Adipo DEGs")
```

```
## Using assay 'log2FC'
```

if we check how downregulated genes during adipogenesis behave in
adult MSC lines we see that the majority are upregulated, but some are
also downregulated

```
sechm(MixG[,which(MixG$Line == "A"|MixG$Line == "B" )],adipo$h96_DOWN, do.scale = T, show_rownames = F, top_annotation =  c("Line","EXPO2"), name = "Adipo DEGs")
```

```
## Using assay 'log2FC'
```

##### Overlap DEGs and ostegenesis genes from https://doi.org/10.1016/j.stemcr.2017.02.018

```
# library(readxl)
# 
# for (x in c("h0.5","h1","h2","h3","h6","h12","h24","h48","h72","h96")) {
# 
# x <- read_excel(paste0("Data/2023_Data/OsteogenesisDEGS/",x) ,sheet = "Sheet 1")
# paste0(x,"_UP") <- x$...1[which(x$adj.P.Val<0.001 & x$logFC>0)]
# paste0(x,"_DOWN") <- x$...1[which(x$adj.P.Val<0.001& x$logFC<0)]
#   
# }
```

```
library(readxl)


for (x in c("h0.5","h1","h2","h3","h6","h12","h24","h48","h72","h96")) {

  
tmp <- read_excel(paste0(paste0("Data/2023_Data/OsteogenesisDEGS/",x),".xlsx") ,sheet = "Sheet 1")

tmp2<-tmp$Gene[which(tmp$adj.P.Val<0.001)]
tmp3<-tmp$Gene[which(tmp$adj.P.Val<0.001 & tmp$logFC>0)]
tmp4<-tmp$Gene[which(tmp$adj.P.Val<0.001 & tmp$logFC<0)]
assign(paste0(x,"_ALL"), tmp2)
assign(paste0(x,"_UP"), tmp3)
assign(paste0(x,"_DOWN"), tmp4)

}

osteo <- list()
osteo <- list(
"h0.5_ALL"=h0.5_ALL, "h0.5_UP"=h0.5_UP, "h0.5_DOWN"=h0.5_DOWN, 
"h1_ALL"=h1_ALL, "h1_UP"=h1_UP, "h1_DOWN"=h1_DOWN,
"h2_ALL"=h2_ALL, "h2_UP"=h2_UP, "h2_DOWN"=h2_DOWN,
"h3_ALL"=h3_ALL, "h3_UP"=h3_UP, "h3_DOWN"=h3_DOWN,
"h6_ALL"=h6_ALL, "h6_UP"=h6_UP, "h6_DOWN"=h6_DOWN,
"h12_ALL"=h12_ALL, "h12_UP"=h12_UP, "h12_DOWN"=h12_DOWN,
"h24_ALL"=h24_ALL, "h24_UP"=h24_UP, "h24_DOWN"=h24_DOWN,
"h48_ALL"=h48_ALL, "h48_UP"=h48_UP, "h48_DOWN"=h48_DOWN,
"h72_ALL"=h72_ALL, "h72_UP"=h72_UP, "h72_DOWN"=h72_DOWN,
"h96_ALL"=h96_ALL, "h96_UP"=h96_UP, "h96_DOWN"=h96_DOWN
)
```

```
# library(readxl)
# h3 <- read_excel("Data/Adipogenesis/DEG_0hr_vs_3hr.xlsx",sheet = "Sheet1")
# h3UP <- h3$Gene[which(h3$adj.P.Val<0.001 & h3$logFC>0)]
# h3DOWN <- h3$Gene[which(h3$adj.P.Val<0.001& h3$logFC<0)]

# h6 <- read_excel("Data/Adipogenesis/DEG_0hr_vs_6hr.xlsx",sheet = "Sheet1")
# h6 <- h6$Gene[which(h6$adj.P.Val<0.001)]
# 
# h24 <- read_excel("Data/Adipogenesis/DEG_0hr_vs_24hr.xlsx",sheet = "Sheet1")
# h24 <- h24$Gene[which(h24$adj.P.Val<0.001)]
# 
# h48 <- read_excel("Data/Adipogenesis/DEG_0hr_vs_48hr.xlsx",sheet = "Sheet1")
# h48 <- h48$Gene[which(h48$adj.P.Val<0.001)]
# 
# h96 <- read_excel("Data/Adipogenesis/DEG_0hr_vs_96hr.xlsx",sheet = "Sheet1")
# h96 <- h96$Gene[which(h96$adj.P.Val<0.001)]
# 
# adipo <- list(h6=h6,h24=h24)
```

###### only osteogenesis DEGs vs themselves

```
uni <-rownames(filterGenes(SEs$acute.msc))

m <- overlapper::multintersect(osteo, universe = uni)
overlapper::dotplot.multintersect(m)
## Warning: Removed 378 rows containing missing values (`geom_point()`).
## Warning: Removed 378 rows containing missing values (`geom_text()`).
```

###### MixG\_DEGS vs osteogenesis DEGs

```
m <- overlapper::multintersect(MixG_DEGS, osteo, universe = uni)
overlapper::dotplot.multintersect(m)
```

```
write.table(m$of, file = "Data/2023_Data/overlaps_osteo.txt", sep = "\t")
#write.csv(m$of, file = "Data/2023_Data/overlaps.csv")

#save overlapping genes for GO analysis
osteo_DOWN <- intersect(MixG_DEGS$MSC_DEGs_UP, osteo$h96_DOWN)
osteo_UP <- intersect(MixG_DEGS$MSC_DEGs_DOWN, osteo$h96_UP)


save(osteo_DOWN,osteo_UP, file = "Data/osteo.RData")
```

##### Heatmaps

###### heaatmap to check osteogenesis genes in our dataset

if we check how upregulated genes during osteogenesis behave in adult
MSC lines we see that the majority are downregulated, but some are also
upregulated

```
sechm(MixG[,which(MixG$Line == "A"|MixG$Line == "B" )],osteo$h96_UP, do.scale = T, show_rownames = F, top_annotation =  c("Line","EXPO2"), name = "OSTEO DEGs UP")
```

```
## Using assay 'log2FC'
```

if we check how downregulated genes during osteogenesis behave in
adult MSC lines we see that the majority are upregulated, but some are
also downregulated

```
sechm(MixG[,which(MixG$Line == "A"|MixG$Line == "B" )],osteo$h96_DOWN, do.scale = T, show_rownames = F, top_annotation =  c("Line","EXPO2"), name = "OSTEO DEGs DOWN")
```

```
## Using assay 'log2FC'
```

###### osteogenesis vs adipogenesis DEGs

```
uni <-rownames(filterGenes(SEs$acute.msc))

m <- overlapper::multintersect(adipo,osteo, universe = uni)
overlapper::dotplot.multintersect(m)
## Warning: Removed 1 rows containing missing values (`geom_point()`).
```

##### Supervised exploration of interesting genes

from this article:https://doi.org/10.1186/s13287-019-1498-0

Osteogenic inhibition FOXO1 - Osteogenesis and inhibition of
adipogenesis - Downregulated FGF2 - Osteogenesis - Downregulated SOX4 -
Osteogenesis - Downregulated COL1A1 - Osteogenesis and inhibition of
adipogenesis - Downregulated COL1A2 - Osteogenesis and inhibition of
adipogenesis - Downregulated TGFB2 - Osteogenesis - Downregulated JAG1 -
Osteogenesis - Downregulated HEY2 - Osteogenesis - Downregulated ALPL -
Osteogenesis - Downregulated TGFBR2 - Inhibit of osteogenesis -
Upregulated SFRP1 - inhibitor of the Wnt pathway - Upregulated

Adipogenic PTGER2 - Inhibit oste, promote adipo - Upregulated WNT5A -
Adipogenesis/obesity - Upregulated

Osteogenic/inhibition of adipogenesis c-Cbl - Osteogenesis
-Upregulated CEBPD - Adipogenesis - Downregulated CEBPG - Adipogenesis -
Downregulated WWTR1 - Osteogenesis - Upregulated (TAZ)

```
OsteogenicInhibition <- c("FOXO1", "FGF2", "SOX4", "COL1A1", "COL1A2", "TGFB2", "JAG1", "HEY2", "ALPL", "TGFBR2","SFRP1")
Adipogenic<- c("PTGER2", "WNT5A")
AdipogenicInhibition <- c("CBL", "CEBPD", "CEBPG", "WWTR1")
```

OsteogenicInhibition in adult MSC lines

```
sechm(MixG[,which(MixG$Line == "A"|MixG$Line == "B" )], OsteogenicInhibition, do.scale = T, show_rownames = T, top_annotation =  c("Line","EXPO2") )
```

```
## Using assay 'log2FC'
```

OsteogenicInhibition in female iPSC MSC lines

```
sechm(MixG[,which(MixG$Line == "3391S" )],OsteogenicInhibition, do.scale = T, show_rownames = T, top_annotation =  c("Line","EXPO2"))
```

```
## Using assay 'log2FC'
```

OsteogenicInhibition in male iPSC MSC lines

```
sechm(MixG[,which(MixG$Line == "MIFF3")],OsteogenicInhibition, do.scale = T, show_rownames = T, top_annotation =  c("Line","EXPO2"))
```

```
## Using assay 'log2FC'
```

Adipogenic in adult MSC lines

```
sechm(MixG[,which(MixG$Line == "A"|MixG$Line == "B" )], Adipogenic, do.scale = T, show_rownames = T, top_annotation =  c("Line","EXPO2"))
```

```
## Using assay 'log2FC'
```

Adipogenic in female iPSC MSC lines

```
sechm(MixG[,which(MixG$Line == "3391S" )],Adipogenic, do.scale = T, show_rownames = T, top_annotation =  c("Line","EXPO2"))
```

```
## Using assay 'log2FC'
```

Adipogenic in male iPSC MSC lines

```
sechm(MixG[,which(MixG$Line == "MIFF3")],Adipogenic, do.scale = T, show_rownames = T, top_annotation =  c("Line","EXPO2"))
```

```
## Using assay 'log2FC'
```

AdipogenicInhibition in adult MSC lines

```
sechm(MixG[,which(MixG$Line == "A"|MixG$Line == "B" )], AdipogenicInhibition, do.scale = T, show_rownames = T, top_annotation =  c("Line","EXPO2"))
```

```
## Using assay 'log2FC'
```

AdipogenicInhibition in female iPSC MSC lines

```
sechm(MixG[,which(MixG$Line == "3391S" )],AdipogenicInhibition, do.scale = T, show_rownames = T, top_annotation =  c("Line","EXPO2"))
```

```
## Using assay 'log2FC'
```

AdipogenicInhibition in male iPSC MSC lines

```
sechm(MixG[,which(MixG$Line == "MIFF3")],AdipogenicInhibition, do.scale = T, show_rownames = T, top_annotation =  c("Line","EXPO2"))
```

```
## Using assay 'log2FC'
```

##### Data praparation

For details on data filtering, normalization, batch correction and
differential expression analysis, see here

---

To continue with the results you can go here

##### Index

01\_MixG\_MesenchimalStemCells

02\_FunctionalMSCMixG

##### Authors

Nicolò Caporale:

Cristina Cheroni:

Pierre-Luc Germain:

Giuseppe Testa:

Lab: http://www.testalab.eu/

‘Date: November 13, 2024’
