## Supplementary Notebooks Bioinformatic Analysis for "A mixture of endocrine disrupting chemicals linked to lower birth weight induces adipogenesis and transcriptional changes related to birth weight alterations and diabetes": 02_FunctionalMSCMixG.html

RNASeq Characterization of differential expression results: MixG acute treatment in Mesenchymal Stem Cells


Code 

- Show All Code
- Hide All Code

### RNASeq Characterization of differential expression results: MixG acute treatment in Mesenchymal Stem Cells

##### 1. Environment Set Up

###### 1.1 Parameters

```
for (i in 1:length(params))
  print(paste('Parameter:', names(params)[i], ' - Value:', params[[i]], '- Class:', class(params[[i]])))
## [1] "Parameter: Dataset  - Value: acute.msc - Class: character"
## [1] "Parameter: topTagFile  - Value: Data/AllSEcorrected_MSC.RData - Class: character"
## [1] "Parameter: FDRTr  - Value: 0.05 - Class: numeric"
## [1] "Parameter: LogFCTr  - Value: 0.5 - Class: numeric"
## [1] "Parameter: OutputFolder  - Value: Data/FunctionalAnalysis/02_MSCMixG/ - Class: character"
## [1] "Parameter: GOSeq  - Value: Yes - Class: character"
## [1] "Parameter: TopGO  - Value: Yes - Class: character"
## [1] "Parameter: Camera  - Value: Yes - Class: character"
## [1] "Parameter: Gsea  - Value: Yes - Class: character"
## [1] "Parameter: GseaPvalSel  - Value: 0.05 - Class: numeric"
```

```
Dataset <- params$Dataset
LogFCTh <- params$LogFCTr
FDRTh <- params$FDRTr
OutputFolder <- ifelse(is.null(params$OutputFolder), getwd(), params$OutputFolder) 

if (dir.exists(OutputFolder) == FALSE) {
  dir.create(OutputFolder, recursive=FALSE)
}

if (dir.exists(paste0(OutputFolder, 'TopGO')) == FALSE) {
  dir.create(paste0(OutputFolder, 'TopGO'), recursive=FALSE)
}

if (dir.exists(paste0(OutputFolder, 'Gsea')) == FALSE) {
  dir.create(paste0(OutputFolder, 'Gsea'), recursive=FALSE)
}
```

###### 1.2 Libraries and functions

```
library(stringr)
## Warning: package 'stringr' was built under R version 4.3.1
library(DT)
## Warning: package 'DT' was built under R version 4.3.1
library(ggplot2)
library(ggrepel)
library(AnnotationDbi)
## Warning: package 'AnnotationDbi' was built under R version 4.3.1
## Loading required package: stats4
## Loading required package: BiocGenerics
## 
## Attaching package: 'BiocGenerics'
## The following objects are masked from 'package:stats':
## 
##     IQR, mad, sd, var, xtabs
## The following objects are masked from 'package:base':
## 
##     anyDuplicated, aperm, append, as.data.frame, basename, cbind,
##     colnames, dirname, do.call, duplicated, eval, evalq, Filter, Find,
##     get, grep, grepl, intersect, is.unsorted, lapply, Map, mapply,
##     match, mget, order, paste, pmax, pmax.int, pmin, pmin.int,
##     Position, rank, rbind, Reduce, rownames, sapply, setdiff, sort,
##     table, tapply, union, unique, unsplit, which.max, which.min
## Loading required package: Biobase
## Welcome to Bioconductor
## 
##     Vignettes contain introductory material; view with
##     'browseVignettes()'. To cite Bioconductor, see
##     'citation("Biobase")', and for packages 'citation("pkgname")'.
## Loading required package: IRanges
## Warning: package 'IRanges' was built under R version 4.3.1
## Loading required package: S4Vectors
## Warning: package 'S4Vectors' was built under R version 4.3.1
## 
## Attaching package: 'S4Vectors'
## The following object is masked from 'package:utils':
## 
##     findMatches
## The following objects are masked from 'package:base':
## 
##     expand.grid, I, unname
library(org.Hs.eg.db)
## 
library(gridExtra)
## 
## Attaching package: 'gridExtra'
## The following object is masked from 'package:Biobase':
## 
##     combine
## The following object is masked from 'package:BiocGenerics':
## 
##     combine
library(RColorBrewer)
library(topGO)
## Loading required package: graph
## 
## Attaching package: 'graph'
## The following object is masked from 'package:stringr':
## 
##     boundary
## Loading required package: GO.db
## 
## Loading required package: SparseM
## 
## Attaching package: 'SparseM'
## The following object is masked from 'package:base':
## 
##     backsolve
## 
## groupGOTerms:    GOBPTerm, GOMFTerm, GOCCTerm environments built.
## 
## Attaching package: 'topGO'
## The following object is masked from 'package:IRanges':
## 
##     members
library(fgsea)
library(tidyr)
## 
## Attaching package: 'tidyr'
## The following object is masked from 'package:S4Vectors':
## 
##     expand
library(dplyr)
## 
## Attaching package: 'dplyr'
## The following object is masked from 'package:graph':
## 
##     union
## The following object is masked from 'package:gridExtra':
## 
##     combine
## The following object is masked from 'package:AnnotationDbi':
## 
##     select
## The following objects are masked from 'package:IRanges':
## 
##     collapse, desc, intersect, setdiff, slice, union
## The following objects are masked from 'package:S4Vectors':
## 
##     first, intersect, rename, setdiff, setequal, union
## The following object is masked from 'package:Biobase':
## 
##     combine
## The following objects are masked from 'package:BiocGenerics':
## 
##     combine, intersect, setdiff, union
## The following objects are masked from 'package:stats':
## 
##     filter, lag
## The following objects are masked from 'package:base':
## 
##     intersect, setdiff, setequal, union
```

```
options(stringsAsFactors=FALSE)
```

```
source('Functions/EDC_Functions.R')
source('Functions/CriFormatted.R')
source('Functions/goseq.R')
```

###### 1.3 Gene sets

- Molecular Signature Database: H1 (hallmark gene sets); KEGG gene
  sets
- ASD-related: list containing most relevant ASD or NDD-related
  genes
- Metabolism-related

```
H1 <- gmtPathways('Data/MolecularSignatureDatabase/H/h.all.v7.0.symbols.gmt')
Kegg <- gmtPathways('Data/MolecularSignatureDatabase/C2/KEGG/c2.cp.kegg.v7.0.symbols.gmt')

load('Data/ASD.RData', verbose=TRUE)
## Loading objects:
##   ASD
##   SFARI
##   SFARIgenes
##   AutismKB
##   DecipherNeuro
##   MSSNG
##   ASDsevere
##   ID
##   NeuroOmim
##   iPSYCH
##   GDI
# remove not useful ones. I keep the gene sets in ASD list
rm(SFARI, SFARIgenes, AutismKB, DecipherNeuro, 
   MSSNG, ASDsevere, ID, NeuroOmim, iPSYCH, GDI)

load('Data/MetabGenes.RData', verbose=TRUE)
## Loading objects:
##   metabolism
MetabGeneSet <- metabolism
```

---

##### 2. Data Upload: differential expression results

- Upload the result of differential expression analysis.
- \_\_Select DEGs as having an FDR < 0.05 (anova-like FDR) and a
  log2FC > 0.5 in each MixG expo condition. Calculate the mean
  fold-change in the different treatments.

```
load(params$topTagFile, verbose=TRUE)
## Loading objects:
##   SEs
##   DEAs

Res <- list()
Res$tableRaw <- DEAs$acute.msc
Res$table <- DEAs$acute.msc
Res$table$genes <- row.names(Res$table)
Res$table$logFC <- (Res$table$logFC.EXPO0.1X + Res$table$logFC.EXPO1X + Res$table$logFC.EXPO10X + Res$table$logFC.EXPO100X + Res$table$logFC.EXPO1000X)/5
Res$table$GeneName <- row.names(Res$table) # I generated this column to make the dataframe compatible with some functions
Res$degs <- dplyr::filter(Res$table, FDR < FDRTh & (abs(logFC.EXPO0.1X) > LogFCTh | abs(logFC.EXPO1X) > LogFCTh | abs(logFC.EXPO10X) > LogFCTh | abs(logFC.EXPO100X) > LogFCTh | abs(logFC.EXPO1000X) > LogFCTh) & logCPM >0)

dim(Res$degs) 
## [1] 1604   12

### subset overlap MixG DEGs and adipogenesis genes from https://doi.org/10.1016/j.stemcr.2017.02.018
load("Data/adipo.RData")
length(adipo_UP)
## [1] 149
length(adipo_DOWN)
## [1] 282
Res$Adipo_UP <- Res$degs[adipo_UP,]
Res$Adipo_DOWN <- Res$degs[adipo_DOWN,]

### subset overlap MixG DEGs and osteogenesis genes from https://doi.org/10.1016/j.stemcr.2017.02.018
load("Data/osteo.RData")
length(osteo_UP)
## [1] 236
length(osteo_DOWN)
## [1] 283

Res$Osteo_UP <- Res$degs[osteo_UP,]
Res$Osteo_DOWN <- Res$degs[osteo_DOWN,]

### subset overlap MixG DEGs and Corticoid genes from https://www.gsea-msigdb.org/gsea/msigdb
load("Data/Overlap_MixG_Corticoids.RData", verbose = T)
## Loading objects:
##   overlap
length(overlap$MixG_DEGS)
## [1] 1604
length(overlap$Corticoids)
## [1] 2391
length(overlap$intersection)
## [1] 396
Res$MixG_Corticoid <- Res$degs[overlap$intersection,]
```

**12827** genes have been testes for differential
expression.

Imposing a threshold of 1.4142136 on the FC and 0.05 on the FDR (as
specified in parameters), **1604** genes are selected.

---

##### 3. RESULTS NAVIGATION: Interactive Table

From topTag I generate an interactive table for result interrogation
with link to the Gene Cards. **The table reports all the genes
having a FDR < 0.05 and a Log2FC > 0.5** for at least one
of the two treatments as absolute value, according to the threshold
settings.

```
searchURL <- 'https://www.genecards.org/cgi-bin/carddisp.pl?gene='
# First part of the URL that will be used to generate the link

Res$table %>% 
  dplyr::filter(FDR < FDRTh & (abs(logFC.EXPO0.1X) > LogFCTh | abs(logFC.EXPO1X) > LogFCTh | abs(logFC.EXPO10X) > LogFCTh | abs(logFC.EXPO100X) > LogFCTh | abs(logFC.EXPO1000X) > LogFCTh) & logCPM >0) %>%
  dplyr::mutate(GeneLink=paste0('<a href="', searchURL, genes, '">', genes, '</a>')) %>% 
  # generation of the link
  dplyr::select(12, 13, 11, 1:5, 9) %>% # selection of columns to be shown
  DT::datatable(class = 'hover', rownames=FALSE, caption='Differential expression results', filter='top', 
            extensions='Buttons', options = list(pageLength=10, autoWidth=TRUE, dom='Bfrtip', 
                                                 buttons=c('csv', 'excel')), escape=FALSE) %>%
  formatRound(c(3:6), c(rep(2,3), 4))
```

##### 3.1 RESULTS NAVIGATION: Interactive Table

From topTag I generate an interactive table for result interrogation
with link to the Gene Cards. \_\_The table reports all the genes
overlapping between MIXG DEGs Down and adipogenesis UP genes from https://doi.org/10.1016/j.stemcr.2017.02.018`\_\_

```
searchURL <- 'https://www.genecards.org/cgi-bin/carddisp.pl?gene='
# First part of the URL that will be used to generate the link

Res$Adipo_UP %>% 

  dplyr::mutate(GeneLink=paste0('<a href="', searchURL, genes, '">', genes, '</a>')) %>% 
  # generation of the link
  dplyr::select(12, 13, 11, 1:5, 9) %>% # selection of columns to be shown
  
  DT::datatable(class = 'hover', rownames=FALSE, caption='Differential expression results', filter='top', 
            extensions='Buttons', options = list(pageLength=10, autoWidth=TRUE, dom='Bfrtip', 
                                                 buttons=c('csv', 'excel')), escape=FALSE) %>%
  formatRound(c(3:6), c(rep(2,3), 4))
```

From topTag I generate an interactive table for result interrogation
with link to the Gene Cards. \_\_The table reports all the genes
overlapping between MIXG DEGs UP and adipogenesis DOWN genes from https://doi.org/10.1016/j.stemcr.2017.02.018`\_\_

```
searchURL <- 'https://www.genecards.org/cgi-bin/carddisp.pl?gene='
# First part of the URL that will be used to generate the link

Res$Adipo_DOWN %>% 

  dplyr::mutate(GeneLink=paste0('<a href="', searchURL, genes, '">', genes, '</a>')) %>% 
  # generation of the link
  dplyr::select(12, 13, 11, 1:5, 9) %>% # selection of columns to be shown
  
  DT::datatable(class = 'hover', rownames=FALSE, caption='Differential expression results', filter='top', 
            extensions='Buttons', options = list(pageLength=10, autoWidth=TRUE, dom='Bfrtip', 
                                                 buttons=c('csv', 'excel')), escape=FALSE) %>%
  formatRound(c(3:6), c(rep(2,3), 4))
```

From topTag I generate an interactive table for result interrogation
with link to the Gene Cards. \_\_The table reports all the genes
overlapping between MIXG DEGs Down and osteogenesis UP genes from https://doi.org/10.1016/j.stemcr.2017.02.018`\_\_

```
searchURL <- 'https://www.genecards.org/cgi-bin/carddisp.pl?gene='
# First part of the URL that will be used to generate the link

Res$Osteo_DOWN %>% 

  dplyr::mutate(GeneLink=paste0('<a href="', searchURL, genes, '">', genes, '</a>')) %>% 
  # generation of the link
  dplyr::select(12, 13, 11, 1:5, 9) %>% # selection of columns to be shown
  
  DT::datatable(class = 'hover', rownames=FALSE, caption='Differential expression results', filter='top', 
            extensions='Buttons', options = list(pageLength=10, autoWidth=TRUE, dom='Bfrtip', 
                                                 buttons=c('csv', 'excel')), escape=FALSE) %>%
  formatRound(c(3:6), c(rep(2,3), 4))
```

From topTag I generate an interactive table for result interrogation
with link to the Gene Cards. \_\_The table reports all the genes
overlapping between MIXG DEGs UP and osteogenesis DOWN genes from https://doi.org/10.1016/j.stemcr.2017.02.018`\_\_

```
searchURL <- 'https://www.genecards.org/cgi-bin/carddisp.pl?gene='
# First part of the URL that will be used to generate the link

Res$Osteo_DOWN %>% 

  dplyr::mutate(GeneLink=paste0('<a href="', searchURL, genes, '">', genes, '</a>')) %>% 
  # generation of the link
  dplyr::select(12, 13, 11, 1:5, 9) %>% # selection of columns to be shown
  
  DT::datatable(class = 'hover', rownames=FALSE, caption='Differential expression results', filter='top', 
            extensions='Buttons', options = list(pageLength=10, autoWidth=TRUE, dom='Bfrtip', 
                                                 buttons=c('csv', 'excel')), escape=FALSE) %>%
  formatRound(c(3:6), c(rep(2,3), 4))
```

From topTag I generate an interactive table for result interrogation
with link to the Gene Cards. **The table reports all the genes
overlapping between MIXG DEGs Corticoid Pathway-related genes from https://www.gsea-msigdb.org/gsea/msigdb**

```
searchURL <- 'https://www.genecards.org/cgi-bin/carddisp.pl?gene='
# First part of the URL that will be used to generate the link

Res$MixG_Corticoid %>% 

  dplyr::mutate(GeneLink=paste0('<a href="', searchURL, genes, '">', genes, '</a>')) %>% 
  # generation of the link
  dplyr::select(12, 13, 11, 1:5, 9) %>% # selection of columns to be shown
  
  DT::datatable(class = 'hover', rownames=FALSE, caption='Differential expression results', filter='top', 
            extensions='Buttons', options = list(pageLength=10, autoWidth=TRUE, dom='Bfrtip', 
                                                 buttons=c('csv', 'excel')), escape=FALSE) %>%
  formatRound(c(3:6), c(rep(2,3), 4))
```

---

##### 4. TOPGO for Gene Ontology Enrichment analysis

Gene ontology enrichment analysis is performed on the set of 1604
genes using TopGO with Fisher statistics and weight01 algorithm. The
division between up-regulated and down-regulated genes is done on the
mean FC between the two treatments.

###### 4.1 Selection of modulated genes and generation of gene vectors

I generate vectors for the gene universe, all modulated genes,
up-regulated genes and down-regulated genes in the format required by
TopGO.

```
UniverseGenes <- Res$table %>% dplyr::pull(GeneName)
DEG <- Res$degs %>% dplyr::pull(GeneName)
DEGUp <- Res$degs %>% dplyr::filter(logFC > 0) %>% dplyr::pull(GeneName) 
DEGDown <- Res$degs %>% dplyr::filter(logFC < 0) %>% dplyr::pull(GeneName) 
  
# generation of named vectors in the format required by TopGO
GeneVectors <- list()
GeneVectors$DEGenes <- factor(as.integer(UniverseGenes%in%DEG))
names(GeneVectors$DEGenes) <- UniverseGenes
GeneVectors$DEGenesDown <- factor(as.integer(UniverseGenes%in%DEGDown))
names(GeneVectors$DEGenesDown) <- UniverseGenes
GeneVectors$DEGenesUp <- factor(as.integer(UniverseGenes%in%DEGUp))
names(GeneVectors$DEGenesUp) <- UniverseGenes
```

Therefore:

- universe genes: **12827** genes
- modulated genes: **1604** genes
- down-regulated genes: **809** genes of interest
- up-regulated genes: **795** genes of interest

```
BpEval <- ifelse(params$TopGO=='Yes', TRUE, FALSE)
# the analysis is not done if the number of DEGs (all, down-reg or up-reg) is lower than 2.
BpEval <- ifelse(table(GeneVectors$DEGenes)['1'] < 2 | is.na(table(GeneVectors$DEGenes)['1']) | table(GeneVectors$DEGenesDown)['1'] < 2 | is.na(table(GeneVectors$DEGenesDown)['1']) | table(GeneVectors$DEGenesUp)['1'] < 2 | is.na(table(GeneVectors$DEGenesUp)['1']), FALSE, BpEval)
```

On the basis of the analysis settings or the number of differentially
expressed genes, TopGO analysis **IS performed**.

###### 4.2 Biological Process for ALL modulated: 1604 genes

```
# I generate a list that contains the association between each gene and the GO terms that are associated to it
BPann <- topGO::annFUN.org(whichOnto="BP", feasibleGenes=names(GeneVectors$DEGenes), mapping="org.Hs.eg.db", ID="symbol") %>% inverseList()

# Wrapper function for topGO analysis (see helper file)
ResBPAll <- topGOResults(Genes=GeneVectors$DEGenes, gene2GO=BPann, ontology='BP', description=NULL, nodeSize=10, algorithm='weight01', statistic='fisher', EnTh=2, PvalTh=0.01, minTerms=10)
## 
## Building most specific GOs .....
##  ( 10828 GO terms found. )
## 
## Build GO DAG topology ..........
##  ( 14356 GO terms and 32221 relations. )
## 
## Annotating nodes ...............
##  ( 10648 genes annotated to the GO terms. )
## 
##           -- Weight01 Algorithm -- 
## 
##       the algorithm is scoring 5251 nontrivial nodes
##       parameters: 
##           test statistic: fisher
## 
##   Level 17:  1 nodes to be scored    (0 eliminated genes)
## 
##   Level 16:  11 nodes to be scored   (0 eliminated genes)
## 
##   Level 15:  39 nodes to be scored   (20 eliminated genes)
## 
##   Level 14:  61 nodes to be scored   (153 eliminated genes)
## 
##   Level 13:  94 nodes to be scored   (458 eliminated genes)
## 
##   Level 12:  184 nodes to be scored  (1165 eliminated genes)
## 
##   Level 11:  377 nodes to be scored  (2818 eliminated genes)
## 
##   Level 10:  600 nodes to be scored  (4378 eliminated genes)
## 
##   Level 9:   760 nodes to be scored  (5630 eliminated genes)
## 
##   Level 8:   806 nodes to be scored  (7249 eliminated genes)
## 
##   Level 7:   819 nodes to be scored  (8578 eliminated genes)
## 
##   Level 6:   706 nodes to be scored  (9408 eliminated genes)
## 
##   Level 5:   432 nodes to be scored  (9927 eliminated genes)
## 
##   Level 4:   248 nodes to be scored  (10312 eliminated genes)
## 
##   Level 3:   94 nodes to be scored   (10463 eliminated genes)
## 
##   Level 2:   18 nodes to be scored   (10517 eliminated genes)
## 
##   Level 1:   1 nodes to be scored    (10543 eliminated genes)
write.table(ResBPAll$ResAll, file=paste0(OutputFolder, 'TopGO/BPAllResults.txt'), sep='\t', row.names=FALSE)
```

###### 4.3 Biological Process Analysis for DOWN-REGULATED: 809 genes

```
# Wrapper function for topGO analysis (see helper file)
ResBPDown <- topGOResults(Genes=GeneVectors$DEGenesDown, gene2GO=BPann, ontology='BP', description=NULL, nodeSize=10, algorithm='weight01', statistic='fisher', EnTh=2, PvalTh=0.01, minTerms=10)
## 
## Building most specific GOs .....
##  ( 10828 GO terms found. )
## 
## Build GO DAG topology ..........
##  ( 14356 GO terms and 32221 relations. )
## 
## Annotating nodes ...............
##  ( 10648 genes annotated to the GO terms. )
## 
##           -- Weight01 Algorithm -- 
## 
##       the algorithm is scoring 4484 nontrivial nodes
##       parameters: 
##           test statistic: fisher
## 
##   Level 17:  1 nodes to be scored    (0 eliminated genes)
## 
##   Level 16:  6 nodes to be scored    (0 eliminated genes)
## 
##   Level 15:  26 nodes to be scored   (20 eliminated genes)
## 
##   Level 14:  43 nodes to be scored   (92 eliminated genes)
## 
##   Level 13:  66 nodes to be scored   (343 eliminated genes)
## 
##   Level 12:  144 nodes to be scored  (932 eliminated genes)
## 
##   Level 11:  277 nodes to be scored  (2545 eliminated genes)
## 
##   Level 10:  470 nodes to be scored  (4117 eliminated genes)
## 
##   Level 9:   625 nodes to be scored  (5222 eliminated genes)
## 
##   Level 8:   694 nodes to be scored  (6746 eliminated genes)
## 
##   Level 7:   727 nodes to be scored  (8355 eliminated genes)
## 
##   Level 6:   651 nodes to be scored  (9306 eliminated genes)
## 
##   Level 5:   405 nodes to be scored  (9900 eliminated genes)
## 
##   Level 4:   240 nodes to be scored  (10307 eliminated genes)
## 
##   Level 3:   90 nodes to be scored   (10460 eliminated genes)
## 
##   Level 2:   18 nodes to be scored   (10517 eliminated genes)
## 
##   Level 1:   1 nodes to be scored    (10543 eliminated genes)
# Selection on enrichment of at least 2 is implemented (also to avoid depleted categories). Then categories are ranked by PVal and all the ones with Pval < th are selected. If the number is < minTerms, othter terms are included to reach the minimum number. 

# Results are shown in an interactive table
searchURL <- 'http://amigo.geneontology.org/amigo/term/'

ResBPDown$ResSel %>% 
  dplyr::mutate(GOLink=paste0('<a href="', searchURL, GO.ID, '">', GO.ID, '</a>')) %>% 
  dplyr::select(8, 2, 4, 5, 7, 6) %>% # selection of columns to be shown
  #dplyr::filter(as.numeric(Statistics) < 0.01) %>% # only significant terms are shown in the table
  DT::datatable(class = 'hover', rownames = FALSE, caption='Down-regulated genes: Biological Process Gene Ontology Enrichment',  filter='top', extensions='Buttons', options = list(pageLength = 5, autoWidth = TRUE, dom='Bfrtip', buttons=c('csv', 'excel')), escape=FALSE)
```

###### 4.4 Biological Process Analysis for UP-REGULATED: 795 genes

```
# Wrapper function for topGO analysis (see helper file)
ResBPUp <- topGOResults(Genes=GeneVectors$DEGenesUp, gene2GO=BPann, ontology='BP', description=NULL, nodeSize=10, algorithm='weight01', statistic='fisher', EnTh=2, PvalTh=0.01,  minTerms=10)
## 
## Building most specific GOs .....
##  ( 10828 GO terms found. )
## 
## Build GO DAG topology ..........
##  ( 14356 GO terms and 32221 relations. )
## 
## Annotating nodes ...............
##  ( 10648 genes annotated to the GO terms. )
## 
##           -- Weight01 Algorithm -- 
## 
##       the algorithm is scoring 4704 nontrivial nodes
##       parameters: 
##           test statistic: fisher
## 
##   Level 17:  1 nodes to be scored    (0 eliminated genes)
## 
##   Level 16:  10 nodes to be scored   (0 eliminated genes)
## 
##   Level 15:  33 nodes to be scored   (20 eliminated genes)
## 
##   Level 14:  52 nodes to be scored   (140 eliminated genes)
## 
##   Level 13:  86 nodes to be scored   (407 eliminated genes)
## 
##   Level 12:  156 nodes to be scored  (1022 eliminated genes)
## 
##   Level 11:  324 nodes to be scored  (2768 eliminated genes)
## 
##   Level 10:  505 nodes to be scored  (4190 eliminated genes)
## 
##   Level 9:   668 nodes to be scored  (5447 eliminated genes)
## 
##   Level 8:   731 nodes to be scored  (7022 eliminated genes)
## 
##   Level 7:   745 nodes to be scored  (8447 eliminated genes)
## 
##   Level 6:   646 nodes to be scored  (9349 eliminated genes)
## 
##   Level 5:   409 nodes to be scored  (9893 eliminated genes)
## 
##   Level 4:   229 nodes to be scored  (10308 eliminated genes)
## 
##   Level 3:   90 nodes to be scored   (10462 eliminated genes)
## 
##   Level 2:   18 nodes to be scored   (10517 eliminated genes)
## 
##   Level 1:   1 nodes to be scored    (10543 eliminated genes)
## Warning: There was 1 warning in `dplyr::filter()`.
## ℹ In argument: `as.numeric(Statistics) <= PvalTh`.
## Caused by warning:
## ! NAs introduced by coercion

ResBPUp$ResSel %>% 
  dplyr::mutate(GOLink=paste0('<a href="', searchURL, GO.ID, '">', GO.ID, '</a>')) %>% 
  dplyr::select(8, 2, 4, 5, 7, 6) %>% # selection of columns to be shown
  #dplyr::filter(as.numeric(Statistics) < 0.01) %>% # only significant terms are shown in the table
  DT::datatable(class = 'hover', rownames = FALSE, caption='Up-regulated genes: Biological Process Gene Ontology Enrichment',  filter='top', extensions='Buttons', options = list(pageLength = 5, autoWidth = TRUE, dom='Bfrtip', buttons=c('csv', 'excel')), escape=FALSE)
```

###### 4.5 Result visualization: Barplot

```
# the if clauses avoids an error in case there is one empty category
if(dim(ResBPUp$ResSel)[1] > 0 & dim(ResBPDown$ResSel)[1] > 0 & dim(ResBPAll$ResSel)[1] > 0){

TopGOBar <- topGOBarplotAll_new(TopGOResAll=ResBPAll$ResSel, TopGOResDown=ResBPDown$ResSel, TopGOResUp=ResBPUp$ResSel, terms=8, pvalTh=0.01)

TopGOBar

ggsave(filename=paste0(OutputFolder, '/TopGO/Barplot.pdf'), TopGOBar, 
       width=13, height=8)
  }
## Warning in topGOBarplotAll_new(TopGOResAll = ResBPAll$ResSel, TopGOResDown = ResBPDown$ResSel, : If you did specify the 'cols' argument please make sure to corectly
##                 set names. See `?topGOBarplotAll`
```

##### TOPGO for Gene Ontology Enrichment analysis for overlapping genes between MIXG DEGs DOWN and adipogenesis UP genes

###### Selection of modulated genes and generation of gene vectors

I generate vectors for the gene universe, all modulated genes,
up-regulated genes and down-regulated genes in the format required by
TopGO.

```
UniverseGenes <- Res$table %>% dplyr::pull(GeneName)
DEG <- Res$Adipo_UP %>% dplyr::pull(GeneName)
#DEGUp <- Res$Adipo %>% dplyr::filter(logFC > 0) %>% dplyr::pull(GeneName) 
#DEGDown <- Res$Adipo %>% dplyr::filter(logFC < 0) %>% dplyr::pull(GeneName) 
  
# generation of named vectors in the format required by TopGO
GeneVectors <- list()
GeneVectors$DEGenes <- factor(as.integer(UniverseGenes%in%DEG))
names(GeneVectors$DEGenes) <- UniverseGenes
# GeneVectors$DEGenesDown <- factor(as.integer(UniverseGenes%in%DEGDown))
# names(GeneVectors$DEGenesDown) <- UniverseGenes
# GeneVectors$DEGenesUp <- factor(as.integer(UniverseGenes%in%DEGUp))
# names(GeneVectors$DEGenesUp) <- UniverseGenes
```

Therefore:

- universe genes: **12827** genes
- modulated genes: **149** genes

```
BpEval <- ifelse(params$TopGO=='Yes', TRUE, FALSE)
# the analysis is not done if the number of DEGs (all, down-reg or up-reg) is lower than 2.
BpEval <- ifelse(table(GeneVectors$DEGenes)['1'] < 2 | is.na(table(GeneVectors$DEGenes)['1']) | table(GeneVectors$DEGenesDown)['1'] < 2 | is.na(table(GeneVectors$DEGenesDown)['1']) | table(GeneVectors$DEGenesUp)['1'] < 2 | is.na(table(GeneVectors$DEGenesUp)['1']), FALSE, BpEval)
```

On the basis of the analysis settings or the number of differentially
expressed genes, TopGO analysis **IS NOT performed**.

###### Biological Process for ALL modulated: 149 genes

```
# I generate a list that contains the association between each gene and the GO terms that are associated to it
BPann <- topGO::annFUN.org(whichOnto="BP", feasibleGenes=names(GeneVectors$DEGenes), mapping="org.Hs.eg.db", ID="symbol") %>% inverseList()

# Wrapper function for topGO analysis (see helper file)
ResBPAll <- topGOResults(Genes=GeneVectors$DEGenes, gene2GO=BPann, ontology='BP', description=NULL, nodeSize=10, algorithm='weight01', statistic='fisher', EnTh=2, PvalTh=0.01, minTerms=10)
## 
## Building most specific GOs .....
##  ( 10828 GO terms found. )
## 
## Build GO DAG topology ..........
##  ( 14356 GO terms and 32221 relations. )
## 
## Annotating nodes ...............
##  ( 10648 genes annotated to the GO terms. )
## 
##           -- Weight01 Algorithm -- 
## 
##       the algorithm is scoring 2908 nontrivial nodes
##       parameters: 
##           test statistic: fisher
## 
##   Level 16:  1 nodes to be scored    (0 eliminated genes)
## 
##   Level 15:  10 nodes to be scored   (0 eliminated genes)
## 
##   Level 14:  18 nodes to be scored   (13 eliminated genes)
## 
##   Level 13:  33 nodes to be scored   (197 eliminated genes)
## 
##   Level 12:  77 nodes to be scored   (666 eliminated genes)
## 
##   Level 11:  149 nodes to be scored  (2240 eliminated genes)
## 
##   Level 10:  256 nodes to be scored  (3508 eliminated genes)
## 
##   Level 9:   350 nodes to be scored  (4440 eliminated genes)
## 
##   Level 8:   424 nodes to be scored  (5650 eliminated genes)
## 
##   Level 7:   496 nodes to be scored  (7404 eliminated genes)
## 
##   Level 6:   478 nodes to be scored  (8969 eliminated genes)
## 
##   Level 5:   326 nodes to be scored  (9744 eliminated genes)
## 
##   Level 4:   193 nodes to be scored  (10254 eliminated genes)
## 
##   Level 3:   79 nodes to be scored   (10444 eliminated genes)
## 
##   Level 2:   17 nodes to be scored   (10508 eliminated genes)
## 
##   Level 1:   1 nodes to be scored    (10543 eliminated genes)
write.table(ResBPAll$ResAll, file=paste0(OutputFolder, 'TopGO/BPAllResults.txt'), sep='\t', row.names=FALSE)
```

###### Result visualization: Barplot

```
# the if clauses avoids an error in case there is one empty category
#if(dim(ResBPUp$ResSel)[1] > 0 & dim(ResBPDown$ResSel)[1] > 0 & dim(ResBPAll$ResSel)[1] > 0){

#TopGOBar <- topGOBarplotAll(TopGOResAll=ResBPAll$ResSel, TopGOResDown=ResBPDown$ResSel, TopGOResUp=ResBPUp$ResSel, terms=8, pvalTh=0.01, title=NULL)

TopGOBar <- topGOBarplot_new(TopGORes = ResBPAll$ResSel, terms=8, pvalTh=0.01)

TopGOBar
```

##### TOPGO for Gene Ontology Enrichment analysis for overlapping genes between MIXG DEGs UP and adipogenesis DOWN genes

###### Selection of modulated genes and generation of gene vectors

I generate vectors for the gene universe, all modulated genes,
up-regulated genes and down-regulated genes in the format required by
TopGO.

```
UniverseGenes <- Res$table %>% dplyr::pull(GeneName)
DEG <- Res$Adipo_DOWN %>% dplyr::pull(GeneName)
#DEGUp <- Res$Adipo %>% dplyr::filter(logFC > 0) %>% dplyr::pull(GeneName) 
#DEGDown <- Res$Adipo %>% dplyr::filter(logFC < 0) %>% dplyr::pull(GeneName) 
  
# generation of named vectors in the format required by TopGO
GeneVectors <- list()
GeneVectors$DEGenes <- factor(as.integer(UniverseGenes%in%DEG))
names(GeneVectors$DEGenes) <- UniverseGenes
# GeneVectors$DEGenesDown <- factor(as.integer(UniverseGenes%in%DEGDown))
# names(GeneVectors$DEGenesDown) <- UniverseGenes
# GeneVectors$DEGenesUp <- factor(as.integer(UniverseGenes%in%DEGUp))
# names(GeneVectors$DEGenesUp) <- UniverseGenes
```

Therefore:

- universe genes: **12827** genes
- modulated genes: **282** genes

```
BpEval <- ifelse(params$TopGO=='Yes', TRUE, FALSE)
# the analysis is not done if the number of DEGs (all, down-reg or up-reg) is lower than 2.
BpEval <- ifelse(table(GeneVectors$DEGenes)['1'] < 2 | is.na(table(GeneVectors$DEGenes)['1']) | table(GeneVectors$DEGenesDown)['1'] < 2 | is.na(table(GeneVectors$DEGenesDown)['1']) | table(GeneVectors$DEGenesUp)['1'] < 2 | is.na(table(GeneVectors$DEGenesUp)['1']), FALSE, BpEval)
```

On the basis of the analysis settings or the number of differentially
expressed genes, TopGO analysis **IS NOT performed**.

###### Biological Process for ALL modulated: 282 genes

```
# I generate a list that contains the association between each gene and the GO terms that are associated to it
BPann <- topGO::annFUN.org(whichOnto="BP", feasibleGenes=names(GeneVectors$DEGenes), mapping="org.Hs.eg.db", ID="symbol") %>% inverseList()

# Wrapper function for topGO analysis (see helper file)
ResBPAll <- topGOResults(Genes=GeneVectors$DEGenes, gene2GO=BPann, ontology='BP', description=NULL, nodeSize=10, algorithm='weight01', statistic='fisher', EnTh=2, PvalTh=0.01, minTerms=10)
## 
## Building most specific GOs .....
##  ( 10828 GO terms found. )
## 
## Build GO DAG topology ..........
##  ( 14356 GO terms and 32221 relations. )
## 
## Annotating nodes ...............
##  ( 10648 genes annotated to the GO terms. )
## 
##           -- Weight01 Algorithm -- 
## 
##       the algorithm is scoring 3339 nontrivial nodes
##       parameters: 
##           test statistic: fisher
## 
##   Level 17:  1 nodes to be scored    (0 eliminated genes)
## 
##   Level 16:  9 nodes to be scored    (0 eliminated genes)
## 
##   Level 15:  23 nodes to be scored   (20 eliminated genes)
## 
##   Level 14:  34 nodes to be scored   (131 eliminated genes)
## 
##   Level 13:  59 nodes to be scored   (350 eliminated genes)
## 
##   Level 12:  102 nodes to be scored  (827 eliminated genes)
## 
##   Level 11:  204 nodes to be scored  (2442 eliminated genes)
## 
##   Level 10:  336 nodes to be scored  (3707 eliminated genes)
## 
##   Level 9:   446 nodes to be scored  (4831 eliminated genes)
## 
##   Level 8:   482 nodes to be scored  (6330 eliminated genes)
## 
##   Level 7:   537 nodes to be scored  (7778 eliminated genes)
## 
##   Level 6:   491 nodes to be scored  (9083 eliminated genes)
## 
##   Level 5:   330 nodes to be scored  (9736 eliminated genes)
## 
##   Level 4:   185 nodes to be scored  (10243 eliminated genes)
## 
##   Level 3:   82 nodes to be scored   (10451 eliminated genes)
## 
##   Level 2:   17 nodes to be scored   (10513 eliminated genes)
## 
##   Level 1:   1 nodes to be scored    (10534 eliminated genes)
## Warning: There was 1 warning in `dplyr::filter()`.
## ℹ In argument: `as.numeric(Statistics) <= PvalTh`.
## Caused by warning:
## ! NAs introduced by coercion
write.table(ResBPAll$ResAll, file=paste0(OutputFolder, 'TopGO/BPAllResults.txt'), sep='\t', row.names=FALSE)
```

###### Result visualization: Barplot

```
# the if clauses avoids an error in case there is one empty category
#if(dim(ResBPUp$ResSel)[1] > 0 & dim(ResBPDown$ResSel)[1] > 0 & dim(ResBPAll$ResSel)[1] > 0){

#TopGOBar <- topGOBarplotAll(TopGOResAll=ResBPAll$ResSel, TopGOResDown=ResBPDown$ResSel, TopGOResUp=ResBPUp$ResSel, terms=8, pvalTh=0.01, title=NULL)

TopGOBar <- topGOBarplot_new(TopGORes = ResBPAll$ResSel, terms=8, pvalTh=0.01)

TopGOBar
```

##### TOPGO for Gene Ontology Enrichment analysis for overlapping genes between MIXG DEGs DOWN and osteogenesis UP genes

###### Selection of modulated genes and generation of gene vectors

I generate vectors for the gene universe, all modulated genes,
up-regulated genes and down-regulated genes in the format required by
TopGO.

```
UniverseGenes <- Res$table %>% dplyr::pull(GeneName)
DEG <- Res$Osteo_UP %>% dplyr::pull(GeneName)
#DEGUp <- Res$Adipo %>% dplyr::filter(logFC > 0) %>% dplyr::pull(GeneName) 
#DEGDown <- Res$Adipo %>% dplyr::filter(logFC < 0) %>% dplyr::pull(GeneName) 
  
# generation of named vectors in the format required by TopGO
GeneVectors <- list()
GeneVectors$DEGenes <- factor(as.integer(UniverseGenes%in%DEG))
names(GeneVectors$DEGenes) <- UniverseGenes
# GeneVectors$DEGenesDown <- factor(as.integer(UniverseGenes%in%DEGDown))
# names(GeneVectors$DEGenesDown) <- UniverseGenes
# GeneVectors$DEGenesUp <- factor(as.integer(UniverseGenes%in%DEGUp))
# names(GeneVectors$DEGenesUp) <- UniverseGenes
```

Therefore:

- universe genes: **12827** genes
- modulated genes: **236** genes

```
BpEval <- ifelse(params$TopGO=='Yes', TRUE, FALSE)
# the analysis is not done if the number of DEGs (all, down-reg or up-reg) is lower than 2.
BpEval <- ifelse(table(GeneVectors$DEGenes)['1'] < 2 | is.na(table(GeneVectors$DEGenes)['1']) | table(GeneVectors$DEGenesDown)['1'] < 2 | is.na(table(GeneVectors$DEGenesDown)['1']) | table(GeneVectors$DEGenesUp)['1'] < 2 | is.na(table(GeneVectors$DEGenesUp)['1']), FALSE, BpEval)
```

On the basis of the analysis settings or the number of differentially
expressed genes, TopGO analysis **IS NOT performed**.

###### Biological Process for ALL modulated: 236 genes

```
# I generate a list that contains the association between each gene and the GO terms that are associated to it
BPann <- topGO::annFUN.org(whichOnto="BP", feasibleGenes=names(GeneVectors$DEGenes), mapping="org.Hs.eg.db", ID="symbol") %>% inverseList()

# Wrapper function for topGO analysis (see helper file)
ResBPAll <- topGOResults(Genes=GeneVectors$DEGenes, gene2GO=BPann, ontology='BP', description=NULL, nodeSize=10, algorithm='weight01', statistic='fisher', EnTh=2, PvalTh=0.01, minTerms=10)
## 
## Building most specific GOs .....
##  ( 10828 GO terms found. )
## 
## Build GO DAG topology ..........
##  ( 14356 GO terms and 32221 relations. )
## 
## Annotating nodes ...............
##  ( 10648 genes annotated to the GO terms. )
## 
##           -- Weight01 Algorithm -- 
## 
##       the algorithm is scoring 3301 nontrivial nodes
##       parameters: 
##           test statistic: fisher
## 
##   Level 16:  1 nodes to be scored    (0 eliminated genes)
## 
##   Level 15:  11 nodes to be scored   (0 eliminated genes)
## 
##   Level 14:  24 nodes to be scored   (13 eliminated genes)
## 
##   Level 13:  37 nodes to be scored   (191 eliminated genes)
## 
##   Level 12:  86 nodes to be scored   (742 eliminated genes)
## 
##   Level 11:  178 nodes to be scored  (2280 eliminated genes)
## 
##   Level 10:  306 nodes to be scored  (3574 eliminated genes)
## 
##   Level 9:   418 nodes to be scored  (4678 eliminated genes)
## 
##   Level 8:   487 nodes to be scored  (6106 eliminated genes)
## 
##   Level 7:   566 nodes to be scored  (7813 eliminated genes)
## 
##   Level 6:   527 nodes to be scored  (9080 eliminated genes)
## 
##   Level 5:   354 nodes to be scored  (9820 eliminated genes)
## 
##   Level 4:   208 nodes to be scored  (10279 eliminated genes)
## 
##   Level 3:   81 nodes to be scored   (10449 eliminated genes)
## 
##   Level 2:   16 nodes to be scored   (10517 eliminated genes)
## 
##   Level 1:   1 nodes to be scored    (10543 eliminated genes)
write.table(ResBPAll$ResAll, file=paste0(OutputFolder, 'TopGO/BPAllResults.txt'), sep='\t', row.names=FALSE)
```

###### Result visualization: Barplot

```
# the if clauses avoids an error in case there is one empty category
#if(dim(ResBPUp$ResSel)[1] > 0 & dim(ResBPDown$ResSel)[1] > 0 & dim(ResBPAll$ResSel)[1] > 0){

#TopGOBar <- topGOBarplotAll(TopGOResAll=ResBPAll$ResSel, TopGOResDown=ResBPDown$ResSel, TopGOResUp=ResBPUp$ResSel, terms=8, pvalTh=0.01, title=NULL)

TopGOBar <- topGOBarplot_new(TopGORes = ResBPAll$ResSel, terms=8, pvalTh=0.01)

TopGOBar
```

##### TOPGO for Gene Ontology Enrichment analysis for overlapping genes between MIXG DEGs UP and osteogenesis DOWN genes

###### Selection of modulated genes and generation of gene vectors

I generate vectors for the gene universe, all modulated genes,
up-regulated genes and down-regulated genes in the format required by
TopGO.

```
UniverseGenes <- Res$table %>% dplyr::pull(GeneName)
DEG <- Res$Osteo_DOWN %>% dplyr::pull(GeneName)
#DEGUp <- Res$Adipo %>% dplyr::filter(logFC > 0) %>% dplyr::pull(GeneName) 
#DEGDown <- Res$Adipo %>% dplyr::filter(logFC < 0) %>% dplyr::pull(GeneName) 
  
# generation of named vectors in the format required by TopGO
GeneVectors <- list()
GeneVectors$DEGenes <- factor(as.integer(UniverseGenes%in%DEG))
names(GeneVectors$DEGenes) <- UniverseGenes
# GeneVectors$DEGenesDown <- factor(as.integer(UniverseGenes%in%DEGDown))
# names(GeneVectors$DEGenesDown) <- UniverseGenes
# GeneVectors$DEGenesUp <- factor(as.integer(UniverseGenes%in%DEGUp))
# names(GeneVectors$DEGenesUp) <- UniverseGenes
```

Therefore:

- universe genes: **12827** genes
- modulated genes: **283** genes

```
BpEval <- ifelse(params$TopGO=='Yes', TRUE, FALSE)
# the analysis is not done if the number of DEGs (all, down-reg or up-reg) is lower than 2.
BpEval <- ifelse(table(GeneVectors$DEGenes)['1'] < 2 | is.na(table(GeneVectors$DEGenes)['1']) | table(GeneVectors$DEGenesDown)['1'] < 2 | is.na(table(GeneVectors$DEGenesDown)['1']) | table(GeneVectors$DEGenesUp)['1'] < 2 | is.na(table(GeneVectors$DEGenesUp)['1']), FALSE, BpEval)
```

On the basis of the analysis settings or the number of differentially
expressed genes, TopGO analysis **IS NOT performed**.

###### Biological Process for ALL modulated: 283 genes

```
# I generate a list that contains the association between each gene and the GO terms that are associated to it
BPann <- topGO::annFUN.org(whichOnto="BP", feasibleGenes=names(GeneVectors$DEGenes), mapping="org.Hs.eg.db", ID="symbol") %>% inverseList()

# Wrapper function for topGO analysis (see helper file)
ResBPAll <- topGOResults(Genes=GeneVectors$DEGenes, gene2GO=BPann, ontology='BP', description=NULL, nodeSize=10, algorithm='weight01', statistic='fisher', EnTh=2, PvalTh=0.01, minTerms=10)
## 
## Building most specific GOs .....
##  ( 10828 GO terms found. )
## 
## Build GO DAG topology ..........
##  ( 14356 GO terms and 32221 relations. )
## 
## Annotating nodes ...............
##  ( 10648 genes annotated to the GO terms. )
## 
##           -- Weight01 Algorithm -- 
## 
##       the algorithm is scoring 3543 nontrivial nodes
##       parameters: 
##           test statistic: fisher
## 
##   Level 17:  1 nodes to be scored    (0 eliminated genes)
## 
##   Level 16:  9 nodes to be scored    (0 eliminated genes)
## 
##   Level 15:  27 nodes to be scored   (20 eliminated genes)
## 
##   Level 14:  40 nodes to be scored   (131 eliminated genes)
## 
##   Level 13:  58 nodes to be scored   (368 eliminated genes)
## 
##   Level 12:  109 nodes to be scored  (893 eliminated genes)
## 
##   Level 11:  214 nodes to be scored  (2498 eliminated genes)
## 
##   Level 10:  364 nodes to be scored  (3807 eliminated genes)
## 
##   Level 9:   481 nodes to be scored  (4961 eliminated genes)
## 
##   Level 8:   524 nodes to be scored  (6445 eliminated genes)
## 
##   Level 7:   566 nodes to be scored  (7852 eliminated genes)
## 
##   Level 6:   503 nodes to be scored  (9092 eliminated genes)
## 
##   Level 5:   345 nodes to be scored  (9777 eliminated genes)
## 
##   Level 4:   199 nodes to be scored  (10264 eliminated genes)
## 
##   Level 3:   85 nodes to be scored   (10454 eliminated genes)
## 
##   Level 2:   17 nodes to be scored   (10516 eliminated genes)
## 
##   Level 1:   1 nodes to be scored    (10542 eliminated genes)
write.table(ResBPAll$ResAll, file=paste0(OutputFolder, 'TopGO/BPAllResults.txt'), sep='\t', row.names=FALSE)
```

###### Result visualization: Barplot

```
# the if clauses avoids an error in case there is one empty category
#if(dim(ResBPUp$ResSel)[1] > 0 & dim(ResBPDown$ResSel)[1] > 0 & dim(ResBPAll$ResSel)[1] > 0){

#TopGOBar <- topGOBarplotAll(TopGOResAll=ResBPAll$ResSel, TopGOResDown=ResBPDown$ResSel, TopGOResUp=ResBPUp$ResSel, terms=8, pvalTh=0.01, title=NULL)

TopGOBar <- topGOBarplot_new(TopGORes = ResBPAll$ResSel, terms=8, pvalTh=0.01)

TopGOBar
```

##### TOPGO for Gene Ontology Enrichment analysis for overlapping genes between MIXG DEGs and Corticoid genes

###### Selection of modulated genes and generation of gene vectors

I generate vectors for the gene universe, all modulated genes,
up-regulated genes and down-regulated genes in the format required by
TopGO.

```
UniverseGenes <- Res$table %>% dplyr::pull(GeneName)
DEG <- Res$MixG_Corticoid %>% dplyr::pull(GeneName)
#DEGUp <- Res$Adipo %>% dplyr::filter(logFC > 0) %>% dplyr::pull(GeneName) 
#DEGDown <- Res$Adipo %>% dplyr::filter(logFC < 0) %>% dplyr::pull(GeneName) 
  
# generation of named vectors in the format required by TopGO
GeneVectors <- list()
GeneVectors$DEGenes <- factor(as.integer(UniverseGenes%in%DEG))
names(GeneVectors$DEGenes) <- UniverseGenes
# GeneVectors$DEGenesDown <- factor(as.integer(UniverseGenes%in%DEGDown))
# names(GeneVectors$DEGenesDown) <- UniverseGenes
# GeneVectors$DEGenesUp <- factor(as.integer(UniverseGenes%in%DEGUp))
# names(GeneVectors$DEGenesUp) <- UniverseGenes
```

Therefore:

- universe genes: **12827** genes
- modulated genes: **396** genes

```
BpEval <- ifelse(params$TopGO=='Yes', TRUE, FALSE)
# the analysis is not done if the number of DEGs (all, down-reg or up-reg) is lower than 2.
BpEval <- ifelse(table(GeneVectors$DEGenes)['1'] < 2 | is.na(table(GeneVectors$DEGenes)['1']) | table(GeneVectors$DEGenesDown)['1'] < 2 | is.na(table(GeneVectors$DEGenesDown)['1']) | table(GeneVectors$DEGenesUp)['1'] < 2 | is.na(table(GeneVectors$DEGenesUp)['1']), FALSE, BpEval)
```

On the basis of the analysis settings or the number of differentially
expressed genes, TopGO analysis **IS NOT performed**.

###### Biological Process for ALL modulated: 396 genes

```
# I generate a list that contains the association between each gene and the GO terms that are associated to it
BPann <- topGO::annFUN.org(whichOnto="BP", feasibleGenes=names(GeneVectors$DEGenes), mapping="org.Hs.eg.db", ID="symbol") %>% inverseList()

# Wrapper function for topGO analysis (see helper file)
ResBPAll <- topGOResults(Genes=GeneVectors$DEGenes, gene2GO=BPann, ontology='BP', description=NULL, nodeSize=10, algorithm='weight01', statistic='fisher', EnTh=2, PvalTh=0.01, minTerms=10)
## 
## Building most specific GOs .....
##  ( 10828 GO terms found. )
## 
## Build GO DAG topology ..........
##  ( 14356 GO terms and 32221 relations. )
## 
## Annotating nodes ...............
##  ( 10648 genes annotated to the GO terms. )
## 
##           -- Weight01 Algorithm -- 
## 
##       the algorithm is scoring 4132 nontrivial nodes
##       parameters: 
##           test statistic: fisher
## 
##   Level 17:  1 nodes to be scored    (0 eliminated genes)
## 
##   Level 16:  7 nodes to be scored    (0 eliminated genes)
## 
##   Level 15:  25 nodes to be scored   (20 eliminated genes)
## 
##   Level 14:  41 nodes to be scored   (104 eliminated genes)
## 
##   Level 13:  66 nodes to be scored   (328 eliminated genes)
## 
##   Level 12:  133 nodes to be scored  (945 eliminated genes)
## 
##   Level 11:  264 nodes to be scored  (2621 eliminated genes)
## 
##   Level 10:  429 nodes to be scored  (4126 eliminated genes)
## 
##   Level 9:   571 nodes to be scored  (5216 eliminated genes)
## 
##   Level 8:   626 nodes to be scored  (6644 eliminated genes)
## 
##   Level 7:   674 nodes to be scored  (8285 eliminated genes)
## 
##   Level 6:   603 nodes to be scored  (9261 eliminated genes)
## 
##   Level 5:   376 nodes to be scored  (9875 eliminated genes)
## 
##   Level 4:   211 nodes to be scored  (10299 eliminated genes)
## 
##   Level 3:   88 nodes to be scored   (10462 eliminated genes)
## 
##   Level 2:   16 nodes to be scored   (10516 eliminated genes)
## 
##   Level 1:   1 nodes to be scored    (10543 eliminated genes)
write.table(ResBPAll$ResAll, file=paste0(OutputFolder, 'TopGO/BPAllResults.txt'), sep='\t', row.names=FALSE)
```

###### Result visualization: Barplot

```
# the if clauses avoids an error in case there is one empty category
#if(dim(ResBPUp$ResSel)[1] > 0 & dim(ResBPDown$ResSel)[1] > 0 & dim(ResBPAll$ResSel)[1] > 0){

#TopGOBar <- topGOBarplotAll(TopGOResAll=ResBPAll$ResSel, TopGOResDown=ResBPDown$ResSel, TopGOResUp=ResBPUp$ResSel, terms=8, pvalTh=0.01, title=NULL)

TopGOBar <- topGOBarplot_new(TopGORes = ResBPAll$ResSel, terms=8, pvalTh=0.01)

TopGOBar
```

##### 5. CAMERA for Gene Set analysis

```
CameraEval <- ifelse(params$Camera=='Yes', TRUE, FALSE)
```

On the basis of the analysis settings, Camera analysis **IS
performed**.

```
printTable <- function(x){
  x$genes <- NULL
  datatable(x, filter="top", extensions = 'Buttons', 
            options = list( dom = 'Bfrtip', buttons = 'csv')
            )
}

ResCamera <- Res$table %>% dplyr::mutate(StatSign=F*sign(rowMeans(Res$tableRaw[,grep('logFC\\.',colnames(Res$tableRaw))])))
row.names(ResCamera) <- ResCamera$genes

printTable(cameraWrapper(dea=ResCamera, gsets=NULL, addSets=c(ASD, MetabGeneSet), addDEgenes=TRUE, dea.thres=0.05, minG=5, reportMax=500))
## 
## Attaching package: 'limma'
## The following object is masked from 'package:BiocGenerics':
## 
##     plotMA
```

```
#printTable(cameraWrapper(dea=ResCamera, gsets=c(H1, Kegg, ASD, MetabGeneSet), addSets=NULL, addDEgenes=TRUE, dea.thres=0.05, minG=5, reportMax=500))
```

##### 7. GSEA for Gene Set Enrichment Analysis

```
GseaEval <- ifelse(params$Gsea=='Yes', TRUE, FALSE)
```

On the basis of the analysis settings, GSEA analysis **IS
performed**.

###### 7.1 Ranked Lists for GSEA

**Rank for stat after selecting only genes with PValues <
0.05**. StatSign has been calculated as for camera analysis: F
value multiplied for the sign of the mean FC. **N.B. fgsea sort
the vector in decreasing order.**

```
FStat <- rankGeneVector(ResCamera, Order='StatSign', PValSel=params$GseaPvalSel)
length(FStat)
## [1] 3899
head(FStat)
##  HIST1H3C  HIST1H1A HIST1H2BB  HIST1H4L     PRSS3    PTPN22 
##  31.82806  28.84865  28.75754  26.70545  25.61997  22.25280
tail(FStat)
##      PODN        C3    CLEC3B      IL26     SFRP4       FLG 
## -23.51346 -24.20487 -25.18160 -27.43640 -35.42229 -36.91266
print(paste('Fgsea analysis is performed on', length(FStat), 'genes having PValue <', params$GseaPvalSel, 'and ranked according to signed F statistics'))
## [1] "Fgsea analysis is performed on 3899 genes having PValue < 0.05 and ranked according to signed F statistics"
```

###### 7.2 GSEA analysis

**fgsea analysis: H1 pathways**

```
set.seed(42)
GseaSelH1 <- fgsea(FStat, pathways=H1, nperm=75000, maxSize=500, minSize=10, nproc=1)
## Warning in fgsea(FStat, pathways = H1, nperm = 75000, maxSize = 500, minSize =
## 10, : You are trying to run fgseaSimple. It is recommended to use
## fgseaMultilevel. To run fgseaMultilevel, you need to remove the nperm argument
## in the fgsea function call.
## Warning in preparePathwaysAndStats(pathways, stats, minSize, maxSize, gseaParam, : There are ties in the preranked stats (0.08% of the list).
## The order of those tied genes will be arbitrary, which may produce unexpected results.
##   |                                                                              |                                                                      |   0%  |                                                                              |=                                                                     |   1%  |                                                                              |==                                                                    |   3%  |                                                                              |===                                                                   |   4%  |                                                                              |====                                                                  |   5%  |                                                                              |=====                                                                 |   7%  |                                                                              |======                                                                |   8%  |                                                                              |=======                                                               |   9%  |                                                                              |=======                                                               |  11%  |                                                                              |========                                                              |  12%  |                                                                              |=========                                                             |  13%  |                                                                              |==========                                                            |  15%  |                                                                              |===========                                                           |  16%  |                                                                              |============                                                          |  17%  |                                                                              |=============                                                         |  19%  |                                                                              |==============                                                        |  20%  |                                                                              |===============                                                       |  21%  |                                                                              |================                                                      |  23%  |                                                                              |=================                                                     |  24%  |                                                                              |==================                                                    |  25%  |                                                                              |===================                                                   |  27%  |                                                                              |====================                                                  |  28%  |                                                                              |=====================                                                 |  29%  |                                                                              |=====================                                                 |  31%  |                                                                              |======================                                                |  32%  |                                                                              |=======================                                               |  33%  |                                                                              |========================                                              |  35%  |                                                                              |=========================                                             |  36%  |                                                                              |==========================                                            |  37%  |                                                                              |===========================                                           |  39%  |                                                                              |============================                                          |  40%  |                                                                              |=============================                                         |  41%  |                                                                              |==============================                                        |  43%  |                                                                              |===============================                                       |  44%  |                                                                              |================================                                      |  45%  |                                                                              |=================================                                     |  47%  |                                                                              |==================================                                    |  48%  |                                                                              |===================================                                   |  49%  |                                                                              |===================================                                   |  51%  |                                                                              |====================================                                  |  52%  |                                                                              |=====================================                                 |  53%  |                                                                              |======================================                                |  55%  |                                                                              |=======================================                               |  56%  |                                                                              |========================================                              |  57%  |                                                                              |=========================================                             |  59%  |                                                                              |==========================================                            |  60%  |                                                                              |===========================================                           |  61%  |                                                                              |============================================                          |  63%  |                                                                              |=============================================                         |  64%  |                                                                              |==============================================                        |  65%  |                                                                              |===============================================                       |  67%  |                                                                              |================================================                      |  68%  |                                                                              |=================================================                     |  69%  |                                                                              |=================================================                     |  71%  |                                                                              |==================================================                    |  72%  |                                                                              |===================================================                   |  73%  |                                                                              |====================================================                  |  75%  |                                                                              |=====================================================                 |  76%  |                                                                              |======================================================                |  77%  |                                                                              |=======================================================               |  79%  |                                                                              |========================================================              |  80%  |                                                                              |=========================================================             |  81%  |                                                                              |==========================================================            |  83%  |                                                                              |===========================================================           |  84%  |                                                                              |============================================================          |  85%  |                                                                              |=============================================================         |  87%  |                                                                              |==============================================================        |  88%  |                                                                              |===============================================================       |  89%  |                                                                              |===============================================================       |  91%  |                                                                              |================================================================      |  92%  |                                                                              |=================================================================     |  93%  |                                                                              |==================================================================    |  95%  |                                                                              |===================================================================   |  96%  |                                                                              |====================================================================  |  97%  |                                                                              |===================================================================== |  99%  |                                                                              |======================================================================| 100%
data.table::fwrite(GseaSelH1, file=paste0(OutputFolder, '/Gsea/GseaSelH1.txt'), sep="\t", sep2=c("", " ", ""))

GseaResH1 <- dplyr::filter(GseaSelH1, padj < 0.1 & abs(NES) > 2) %>% 
  dplyr::mutate(LEGenes=stringr::str_count(leadingEdge, ",")+1) %>% 
  dplyr::mutate(GeneSet='H1') %>% 
  dplyr::select(1, 2, 3, 5, 9, 10)
## Warning: There was 1 warning in `dplyr::mutate()`.
## ℹ In argument: `LEGenes = stringr::str_count(leadingEdge, ",") + 1`.
## Caused by warning in `stri_count_regex()`:
## ! argument is not an atomic vector; coercing
GseaResH1[,1]
##                                        pathway
##  1:                   HALLMARK_MITOTIC_SPINDLE
##  2:                        HALLMARK_DNA_REPAIR
##  3:                    HALLMARK_G2M_CHECKPOINT
##  4:                        HALLMARK_MYOGENESIS
##  5:         HALLMARK_INTERFERON_GAMMA_RESPONSE
##  6:                       HALLMARK_E2F_TARGETS
##  7:                    HALLMARK_MYC_TARGETS_V1
##  8:                    HALLMARK_MYC_TARGETS_V2
##  9: HALLMARK_EPITHELIAL_MESENCHYMAL_TRANSITION
## 10:                   HALLMARK_SPERMATOGENESIS
## 11:                 HALLMARK_KRAS_SIGNALING_DN

if(dim(GseaResH1)[1] > 0){
plotGseaTable(H1[GseaResH1 %>% dplyr::arrange(NES) %>% dplyr::pull(pathway)], 
              FStat, GseaSelH1, gseaParam=0.5)
}
```

**fgsea analysis: Kegg pathways**

```
set.seed(42)
GseaSelKegg <- fgsea(FStat, pathways=Kegg, nperm=75000, maxSize=500, minSize=10, nproc=1)
## Warning in fgsea(FStat, pathways = Kegg, nperm = 75000, maxSize = 500, minSize
## = 10, : You are trying to run fgseaSimple. It is recommended to use
## fgseaMultilevel. To run fgseaMultilevel, you need to remove the nperm argument
## in the fgsea function call.
## Warning in preparePathwaysAndStats(pathways, stats, minSize, maxSize, gseaParam, : There are ties in the preranked stats (0.08% of the list).
## The order of those tied genes will be arbitrary, which may produce unexpected results.
##   |                                                                              |                                                                      |   0%  |                                                                              |=                                                                     |   1%  |                                                                              |==                                                                    |   3%  |                                                                              |===                                                                   |   4%  |                                                                              |====                                                                  |   5%  |                                                                              |=====                                                                 |   7%  |                                                                              |======                                                                |   8%  |                                                                              |=======                                                               |   9%  |                                                                              |=======                                                               |  11%  |                                                                              |========                                                              |  12%  |                                                                              |=========                                                             |  13%  |                                                                              |==========                                                            |  15%  |                                                                              |===========                                                           |  16%  |                                                                              |============                                                          |  17%  |                                                                              |=============                                                         |  19%  |                                                                              |==============                                                        |  20%  |                                                                              |===============                                                       |  21%  |                                                                              |================                                                      |  23%  |                                                                              |=================                                                     |  24%  |                                                                              |==================                                                    |  25%  |                                                                              |===================                                                   |  27%  |                                                                              |====================                                                  |  28%  |                                                                              |=====================                                                 |  29%  |                                                                              |=====================                                                 |  31%  |                                                                              |======================                                                |  32%  |                                                                              |=======================                                               |  33%  |                                                                              |========================                                              |  35%  |                                                                              |=========================                                             |  36%  |                                                                              |==========================                                            |  37%  |                                                                              |===========================                                           |  39%  |                                                                              |============================                                          |  40%  |                                                                              |=============================                                         |  41%  |                                                                              |==============================                                        |  43%  |                                                                              |===============================                                       |  44%  |                                                                              |================================                                      |  45%  |                                                                              |=================================                                     |  47%  |                                                                              |==================================                                    |  48%  |                                                                              |===================================                                   |  49%  |                                                                              |===================================                                   |  51%  |                                                                              |====================================                                  |  52%  |                                                                              |=====================================                                 |  53%  |                                                                              |======================================                                |  55%  |                                                                              |=======================================                               |  56%  |                                                                              |========================================                              |  57%  |                                                                              |=========================================                             |  59%  |                                                                              |==========================================                            |  60%  |                                                                              |===========================================                           |  61%  |                                                                              |============================================                          |  63%  |                                                                              |=============================================                         |  64%  |                                                                              |==============================================                        |  65%  |                                                                              |===============================================                       |  67%  |                                                                              |================================================                      |  68%  |                                                                              |=================================================                     |  69%  |                                                                              |=================================================                     |  71%  |                                                                              |==================================================                    |  72%  |                                                                              |===================================================                   |  73%  |                                                                              |====================================================                  |  75%  |                                                                              |=====================================================                 |  76%  |                                                                              |======================================================                |  77%  |                                                                              |=======================================================               |  79%  |                                                                              |========================================================              |  80%  |                                                                              |=========================================================             |  81%  |                                                                              |==========================================================            |  83%  |                                                                              |===========================================================           |  84%  |                                                                              |============================================================          |  85%  |                                                                              |=============================================================         |  87%  |                                                                              |==============================================================        |  88%  |                                                                              |===============================================================       |  89%  |                                                                              |===============================================================       |  91%  |                                                                              |================================================================      |  92%  |                                                                              |=================================================================     |  93%  |                                                                              |==================================================================    |  95%  |                                                                              |===================================================================   |  96%  |                                                                              |====================================================================  |  97%  |                                                                              |===================================================================== |  99%  |                                                                              |======================================================================| 100%
data.table::fwrite(GseaSelKegg, file=paste0(OutputFolder, '/Gsea/GseaSelKegg.txt'), sep="\t", sep2=c("", " ", ""))

GseaResKegg <- dplyr::filter(GseaSelKegg, padj < 0.1 & abs(NES) > 2) %>% 
  dplyr::mutate(LEGenes=stringr::str_count(leadingEdge, ",")+1) %>% 
  dplyr::mutate(GeneSet='Kegg') %>% 
  dplyr::select(1, 2, 3, 5, 9, 10)
## Warning: There was 1 warning in `dplyr::mutate()`.
## ℹ In argument: `LEGenes = stringr::str_count(leadingEdge, ",") + 1`.
## Caused by warning in `stri_count_regex()`:
## ! argument is not an atomic vector; coercing
GseaResKegg[,1]
##                                          pathway
##  1:                   KEGG_PYRIMIDINE_METABOLISM
##  2:                         KEGG_DNA_REPLICATION
##  3:                             KEGG_SPLICEOSOME
##  4:                    KEGG_BASE_EXCISION_REPAIR
##  5:              KEGG_NUCLEOTIDE_EXCISION_REPAIR
##  6:                         KEGG_MISMATCH_REPAIR
##  7:                              KEGG_CELL_CYCLE
##  8:                          KEGG_OOCYTE_MEIOSIS
##  9:                KEGG_ECM_RECEPTOR_INTERACTION
## 10:            KEGG_CELL_ADHESION_MOLECULES_CAMS
## 11: KEGG_PROGESTERONE_MEDIATED_OOCYTE_MATURATION
## 12:            KEGG_SYSTEMIC_LUPUS_ERYTHEMATOSUS
## 13:         KEGG_HYPERTROPHIC_CARDIOMYOPATHY_HCM
## 14:                  KEGG_DILATED_CARDIOMYOPATHY

if(dim(GseaResKegg)[1] > 0){
plotGseaTable(Kegg[GseaResKegg %>% dplyr::arrange(NES) %>% dplyr::pull(pathway)], 
              FStat, GseaSelKegg, gseaParam=0.5)
}
```

###### 7.3 GSEA results visualization

```
GseaRes <- rbind(GseaResH1, GseaResKegg)
Cols <- c(H1='#e76bf3', Kegg="#39b600")

BubbleGsea <- ggplot(GseaRes, aes(x=NES, y=-log10(padj), fill=GeneSet, size=LEGenes)) +
  geom_point(alpha=0.65, shape=21, color='black') + 
  #geom_text_repel(label=GseaRes$pathway, cex=1.8, 
                  #segment.color="grey50", segment.size=0.3,
                  #force=20, nudge_y=-0.15) +
  geom_vline(xintercept = 0, col='blue') +
  xlab('NES') + ylab('-log10 Adjusted PVal') +
  scale_fill_manual(values=Cols) +
  scale_size(range=c(3,15), name='Leading Edge \n Genes') +
  theme_bw()
BubbleGsea
```

```
BubbleGsea <- ggplot(GseaRes, aes(x=NES, y=-log10(padj), fill=GeneSet, size=LEGenes)) +
  geom_point(alpha=0.65, shape=21, color='black') + 
  geom_text_repel(label=GseaRes$pathway, cex=1.8, 
                  segment.color="grey50", segment.size=0.3,
                  force=20, nudge_y=-0.15) +
  geom_vline(xintercept = 0, col='blue') +
  xlab('NES') + ylab('-log10 Adjusted PVal') +
  scale_fill_manual(values=Cols) +
  scale_size(range=c(3,15), name='Leading Edge \n Genes') +
  theme_bw()
BubbleGsea
```

```
ggsave(filename=paste0(OutputFolder, '/Gsea/Bubble.pdf'), BubbleGsea, 
       width=6.5, height=5.5)
```

---

##### 8. Savings

Most of the useful information has been saved during the analysis.
Here I save the workspace and information about the session.

```
SessionInfo <- sessionInfo()
Date <- date()
#save.image(paste0(OutputFolder, 'FunctionalAnalysisWorkspace.RData'))
```

---

##### Index

01\_MixG\_MesenchimalStemCells

02\_FunctionalMSCMixG

##### Authors

Nicolò Caporale:

Cristina Cheroni:

Pierre-Luc Germain:

Giuseppe Testa:

Lab: http://www.testalab.eu/

‘Date: November 13, 2024’
