## Supplementary Notebooks Bioinformatic Analysis for "A mixture of endocrine disrupting chemicals linked to lower birth weight induces adipogenesis and transcriptional changes related to birth weight alterations and diabetes": 03_GWAS.html

GWAS Gene Sets


Code 

- Show All Code
- Hide All Code

### GWAS Gene Sets

###### Purpose

Purpose of this script is to organize the gene sets that have been
donwloaded from GWAS catalogue.

##### Environment Set Up

```
library(tidyr)
library(dplyr)
## 
## Attaching package: 'dplyr'
## The following objects are masked from 'package:stats':
## 
##     filter, lag
## The following objects are masked from 'package:base':
## 
##     intersect, setdiff, setequal, union
library(stringr)
## Warning: package 'stringr' was built under R version 4.3.1
```

```
OutputFolder <- 'DataPreparation/GWAS/'
```

```
GWASCat <- list()
```

---

##### ADHD risk genes

```
GWASCat$ADHD <- list()
GWASCat$ADHD$Info <- 'GWAS Catalogue, Attention Deficit Hyperactivity Disorder (EFO_0003888). Downloaded 2024.01.16 from https://www.ebi.ac.uk/gwas/efotraits/EFO_0003888'


GWASCat$ADHD$Table <- read.table('DataPreparation/GWAS/EFO_0003888_associations_export.tsv', sep = '\t', header = T) %>%
  dplyr::rename(Alleles=riskAllele, PVal=pValue, PValAnnotation=pValueAnnotation, 
                MappedGene=mappedGenes, Trait=traitName, BgTrait=bgTraits, 
                Study=accessionId)

#GWASCat$ADHD$Table$PVal <- GWASCat$ADHD$Table$PVal %>% stringr::str_replace_all(' ', '') %>% 
#  stringr::str_replace('x10', 'E') %>% as.numeric() 

head(GWASCat$ADHD$Table)
##         Alleles  PVal PValAnnotation riskFrequency orValue beta          ci
## 1   rs1979398-? 3e-06              -            NR    1.23    -        [NR]
## 2 rs183882582-T 2e-08              -          0.98    1.43    - [1.26-1.60]
## 3   rs3958046-T 2e-08              -           0.4    1.09    - [1.06-1.10]
## 4 rs200721207-T 4e-08              -          0.66     1.1    - [1.06-1.13]
## 5   rs1920644-T 5e-08              -          0.52    1.09    - [1.05-1.12]
## 6   rs4858241-T 5e-07              -            NR    1.08    - [1.05-1.12]
##        MappedGene                                    Trait
## 1           ITGA1 Attention deficit hyperactivity disorder
## 2 RNU6-293P,RSPH3 Attention deficit hyperactivity disorder
## 3    CADPS2,FEZF1 Attention deficit hyperactivity disorder
## 4 LINC02497,PCDH7 Attention deficit hyperactivity disorder
## 5    RPL6P8,ARL14 Attention deficit hyperactivity disorder
## 6        SGO1-AS1 Attention deficit hyperactivity disorder
##                                  efoTraits BgTrait      Study   locations
## 1 attention deficit hyperactivity disorder       - GCST004778  5:52898497
## 2 attention deficit hyperactivity disorder       - GCST010291 6:158963192
## 3 attention deficit hyperactivity disorder       - GCST010291 7:122315274
## 4 attention deficit hyperactivity disorder       - GCST010291  4:31149844
## 5 attention deficit hyperactivity disorder       - GCST010291 3:160595566
## 6 attention deficit hyperactivity disorder       - GCST010291  3:20627579
##   pubmedId   author
## 1 28809852    Liu L
## 2 32279069 Rovira P
## 3 32279069 Rovira P
## 4 32279069 Rovira P
## 5 32279069 Rovira P
## 6 32279069 Rovira P
```

```
#obtain the genes in a vector
GWASCat$ADHD$Gene <-  GWASCat$ADHD$Table$MappedGene %>% stringr::str_split(', ') %>% unlist() %>% unique()
# discard empty records and sort
GWASCat$ADHD$Gene <- GWASCat$ADHD$Gene[! GWASCat$ADHD$Gene %in% "-"] %>% stringr::str_sort()

head(GWASCat$ADHD$Gene)
## [1] "ABCB9"         "ABHD17C"       "ABI1"          "ACTG1P17"     
## [5] "ACTG1P22"      "ACTG1P22,VRK2"
```

ADHD: table with **1838** entries and
**1288** associated genes.

---

##### Diabetes Mellitus

```
GWASCat$DM <- list()
GWASCat$DM$Info <- 'GWAS Catalogue, Diabetes Mellitus (EFO_0000400). Downloaded 2024.01.16 from https://www.ebi.ac.uk/gwas/efotraits/EFO_0000400'


GWASCat$DM$Table <- read.table('DataPreparation/GWAS/EFO_0000400_associations_export.tsv', sep = '\t', header = T) %>%
  dplyr::rename(Alleles=riskAllele, PVal=pValue, PValAnnotation=pValueAnnotation, 
                MappedGene=mappedGenes, Trait=traitName, BgTrait=bgTraits, 
                Study=accessionId)

#GWASCat$DM$Table$PVal <- GWASCat$DM$Table$PVal %>% stringr::str_replace_all(' ', '') %>% 
#  stringr::str_replace('x10', 'E') %>% as.numeric()

head(GWASCat$DM$Table)
##       Alleles  PVal PValAnnotation riskFrequency   orValue beta
## 1 rs9273368-G 8e-40              -          0.35      2.99    -
## 2 rs9273368-? 3e-78              -         0.301     2.439    -
## 3     rs689-? 1e-18              -         0.715     1.473    -
## 4 rs2476601-? 5e-16              -          0.14     1.529    -
## 5 rs3184504-? 2e-08              -          0.52      1.24    -
## 6 rs1983890-C 3e-08              -         0.642 1.2302363    -
##                                    ci        MappedGene
## 1                         [2.60-3.91] HLA-DQB1,HLA-DQA1
## 2                       [2.222-2.676] HLA-DQB1,HLA-DQA1
## 3                       [1.352-1.605]      INS-IGF2,INS
## 4                        [1.38-1.693]            PTPN22
## 5                       [1.151-1.336]       SH2B3,ATXN2
## 6 [1.14396450049362-1.32301382549448]      RBM17,PFKFB3
##                                            Trait
## 1 Latent autoimmune diabetes vs. type 1 diabetes
## 2 Latent autoimmune diabetes vs. type 2 diabetes
## 3 Latent autoimmune diabetes vs. type 2 diabetes
## 4 Latent autoimmune diabetes vs. type 2 diabetes
## 5 Latent autoimmune diabetes vs. type 2 diabetes
## 6                     Latent autoimmune diabetes
##                                                       efoTraits BgTrait
## 1 type 1 diabetes mellitus,latent autoimmune diabetes in adults       -
## 2 latent autoimmune diabetes in adults,type 2 diabetes mellitus       -
## 3 latent autoimmune diabetes in adults,type 2 diabetes mellitus       -
## 4 latent autoimmune diabetes in adults,type 2 diabetes mellitus       -
## 5 latent autoimmune diabetes in adults,type 2 diabetes mellitus       -
## 6                          latent autoimmune diabetes in adults       -
##        Study    locations pubmedId       author
## 1 GCST007247   6:32658698 30254083 Cousminer DL
## 2 GCST007246   6:32658698 30254083 Cousminer DL
## 3 GCST007246   11:2160994 30254083 Cousminer DL
## 4 GCST007246  1:113834946 30254083 Cousminer DL
## 5 GCST007246 12:111446804 30254083 Cousminer DL
## 6 GCST007245   10:6136651 30254083 Cousminer DL
```

```
#obtain the genes in a vector
GWASCat$DM$Gene <-  GWASCat$DM$Table$MappedGene %>% stringr::str_split(', ') %>% unlist() %>% unique()
# discard empty records and sort
GWASCat$DM$Gene <- GWASCat$DM$Gene[! GWASCat$DM$Gene %in% "-"] %>% stringr::str_sort()

head(GWASCat$DM$Gene)
## [1] "ABCA1"          "ABCA12"         "ABCB10"         "ABCB9"         
## [5] "ABCC1"          "ABCC5,EEF1A1P8"
```

Diabetes Mellitus: table with **6538** entries and
**2184** associated genes.

---

##### Birth Weight

```
GWASCat$BW <- list()
GWASCat$BW$Info <- 'GWAS Catalogue, Birth Weight (EFO_0004344). Downloaded 2024.01.16 from https://www.ebi.ac.uk/gwas/efotraits/EFO_0004344'


GWASCat$BW$Table <- read.table('DataPreparation/GWAS/EFO_0004344_associations_export.tsv', sep = '\t', header = T) %>%
  dplyr::rename(Alleles=riskAllele, PVal=pValue, PValAnnotation=pValueAnnotation, 
                MappedGene=mappedGenes, Trait=traitName, BgTrait=bgTraits, 
                Study=accessionId)

#GWASCat$BW$Table$PVal <- GWASCat$BW$Table$PVal %>% stringr::str_replace_all(' ', '') %>% 
#  stringr::str_replace('x10', 'E') %>% as.numeric()

head(GWASCat$BW$Table)
##         Alleles  PVal PValAnnotation riskFrequency orValue                 beta
## 1   rs7402982-A 1e-09              -          0.42       - 0.0231 unit increase
## 2   rs1351394-T 2e-33              -          0.48       -  0.043 unit increase
## 3 rs144843919-G 2e-09              -          0.96       - 0.0685 unit increase
## 4  rs35261542-C 1e-28              -          0.73       - 0.0444 unit increase
## 5   rs1819436-C 2e-09              -          0.87       - 0.0329 unit increase
## 6 rs138715366-C 1e-26              -          0.99       - 0.2441 unit increase
##              ci         MappedGene        Trait    efoTraits BgTrait      Study
## 1 [0.016-0.031]              IGF1R Birth weight birth weight       - GCST005146
## 2  [0.036-0.05]              HMGA2 Birth weight birth weight       - GCST005146
## 3 [0.046-0.091]  SUZ12P1,RN7SL316P Birth weight birth weight       - GCST005146
## 4 [0.037-0.052]             CDKAL1 Birth weight birth weight       - GCST005146
## 5 [0.022-0.044] LINC00446,OBI1-AS1 Birth weight birth weight       - GCST005146
## 6    [0.2-0.29]               YKT6 Birth weight birth weight       - GCST005146
##     locations pubmedId      author
## 1 15:98650040 27680694 Horikoshi M
## 2 12:65958046 27680694 Horikoshi M
## 3 17:30710321 27680694 Horikoshi M
## 4  6:20675561 27680694 Horikoshi M
## 5 13:78006148 27680694 Horikoshi M
## 6  7:44206672 27680694 Horikoshi M
```

```
#obtain the genes in a vector
GWASCat$BW$Gene <-  GWASCat$BW$Table$MappedGene %>% stringr::str_split(', ') %>% unlist() %>% unique()
# discard empty records and sort
GWASCat$BW$Gene <- GWASCat$BW$Gene[! GWASCat$BW$Gene %in% "-"] %>% stringr::str_sort()

head(GWASCat$BW$Gene)
## [1] "ABCA3"       "ABCC9"       "ABCC9,KCNJ8" "ACVR1C"      "ADAM17"     
## [6] "ADCY5"
```

Birth Weight: table with **601** entries and
**390** associated genes.

---

##### Obesity

```
GWASCat$OB <- list()
GWASCat$OB$Info <- 'GWAS Catalogue, Obesity (EFO_0001073). Downloaded 2024.01.16 from https://www.ebi.ac.uk/gwas/efotraits/EFO_0001073'


GWASCat$OB$Table <- read.table('DataPreparation/GWAS/EFO_0001073_associations_export.tsv', sep = '\t', header = T) %>%
  dplyr::rename(Alleles=riskAllele, PVal=pValue, PValAnnotation=pValueAnnotation, 
                MappedGene=mappedGenes, Trait=traitName, BgTrait=bgTraits, 
                Study=accessionId)

#GWASCat$OB$Table$PVal <- GWASCat$OB$Table$PVal %>% stringr::str_replace_all(' ', '') %>% 
#  stringr::str_replace('x10', 'E') %>% as.numeric()

head(GWASCat$OB$Table)
##        Alleles  PVal PValAnnotation riskFrequency   orValue beta ci
## 1  rs4142322-? 5e-06              -            NR         -    -  -
## 2 rs17573102-? 9e-06              -            NR  8.061831    -  -
## 3     rs9028-? 3e-07              -            NR  5.048128    -  -
## 4  rs7149926-? 1e-06              -            NR  4.771962    -  -
## 5 rs11753543-? 9e-06              -            NR 3.3840287    -  -
## 6  rs9736016-? 3e-07              -            NR 2.4456832    -  -
##          MappedGene                             Trait
## 1 SLC10A2,METTL21EP Obesity without metabolic disease
## 2  ZCCHC10P2,UBL5P1 Obesity without metabolic disease
## 3              RTP4 Obesity without metabolic disease
## 4    BLZF2P,MAGOH3P Obesity without metabolic disease
## 5 SOCS5P5,LINC02518 Obesity without metabolic disease
## 6       CXCR5,Y_RNA Obesity without metabolic disease
##                       efoTraits BgTrait      Study    locations pubmedId
## 1 metabolically healthy obesity       - GCST006492 13:102964708 30120429
## 2 metabolically healthy obesity       - GCST006492   5:29635057 30120429
## 3 metabolically healthy obesity       - GCST006492  3:187371509 30120429
## 4 metabolically healthy obesity       - GCST006492  14:68864011 30120429
## 5 metabolically healthy obesity       - GCST006492  6:113317153 30120429
## 6 metabolically healthy obesity       - GCST006492 11:118854185 30120429
##        author
## 1 Schlauch KA
## 2 Schlauch KA
## 3 Schlauch KA
## 4 Schlauch KA
## 5 Schlauch KA
## 6 Schlauch KA
```

```
#obtain the genes in a vector
GWASCat$OB$Gene <-  GWASCat$OB$Table$MappedGene %>% stringr::str_split(', ') %>% unlist() %>% unique()
# discard empty records and sort
GWASCat$OB$Gene <- GWASCat$OB$Gene[! GWASCat$OB$Gene %in% "-"] %>% stringr::str_sort()

head(GWASCat$OB$Gene)
## [1] "ADCY3"         "ADCY3,DNAJC27" "ADCY9"         "ALPK1"        
## [5] "ANO3"          "ARG1"
```

Obesity: table with **295** entries and
**196** associated genes.

---

##### Saving

```
GWASCat$SessionInfo <- sessionInfo()
saveRDS(GWASCat, paste0(OutputFolder, 'GWASCatalogueGenes.rds'))
```

```
sessionInfo()
```

```
## R version 4.3.0 (2023-04-21)
## Platform: aarch64-apple-darwin20 (64-bit)
## Running under: macOS 14.4
## 
## Matrix products: default
## BLAS:   /Library/Frameworks/R.framework/Versions/4.3-arm64/Resources/lib/libRblas.0.dylib 
## LAPACK: /Library/Frameworks/R.framework/Versions/4.3-arm64/Resources/lib/libRlapack.dylib;  LAPACK version 3.11.0
## 
## locale:
## [1] en_US.UTF-8/en_US.UTF-8/en_US.UTF-8/C/en_US.UTF-8/en_US.UTF-8
## 
## time zone: Europe/Rome
## tzcode source: internal
## 
## attached base packages:
## [1] stats     graphics  grDevices utils     datasets  methods   base     
## 
## other attached packages:
## [1] stringr_1.5.1 dplyr_1.1.2   tidyr_1.3.0  
## 
## loaded via a namespace (and not attached):
##  [1] vctrs_0.6.5       cli_3.6.2         knitr_1.45        rlang_1.1.3      
##  [5] xfun_0.42         stringi_1.8.3     purrr_1.0.2       generics_0.1.3   
##  [9] jsonlite_1.8.8    glue_1.7.0        htmltools_0.5.7   sass_0.4.8       
## [13] fansi_1.0.4       rmarkdown_2.26    evaluate_0.21     jquerylib_0.1.4  
## [17] tibble_3.2.1      fastmap_1.1.1     yaml_2.3.7        lifecycle_1.0.4  
## [21] compiler_4.3.0    pkgconfig_2.0.3   rstudioapi_0.15.0 digest_0.6.34    
## [25] R6_2.5.1          tidyselect_1.2.0  utf8_1.2.3        pillar_1.9.0     
## [29] magrittr_2.0.3    bslib_0.6.1       withr_2.5.0       tools_4.3.0      
## [33] cachem_1.0.8
```

```
date()
```

```
## [1] "Sun Mar 24 15:02:37 2024"
```
