## Supplementary Notebooks Bioinformatic Analysis for "A mixture of endocrine disrupting chemicals linked to lower birth weight induces adipogenesis and transcriptional changes related to birth weight alterations and diabetes": 03b_GWAS_Overlap.html

Overlap MIXG genes and risk genes from GWAS Catalogue


Code 

- Show All Code
- Hide All Code

### Overlap MIXG genes and risk genes from GWAS Catalogue

###### Nicolò Caporale, Cristina Cheroni

###### Date: November 13, 2024

Purpose: overlap between GWAS risk genes and Cortical Brain Organoid
WGCNA modules.

#### 1. SetUp

```
library(GeneOverlap)
library(tidyr)
library(dplyr)
## 
## Attaching package: 'dplyr'
## The following objects are masked from 'package:stats':
## 
##     filter, lag
## The following objects are masked from 'package:base':
## 
##     intersect, setdiff, setequal, union
library(ggplot2)
```

---

#### 3. Data load: gene-phenotype knowledge bases

```
GWAS <- readRDS('DataPreparation/GWAS/GWASCatalogueGenes.rds')
```

---

#### 4. Definition of Gene Universe and Modules

Gene Universe is defined as the genes that have been used for the
generation of gene modules.

##### 4.1 Gene Universe

Define the universe of shared genes between the two networks that
will be used for the overlap analysis.

```
source("Functions/EDC_Functions.R")
source("Functions/CriFormatted.R")
library(overlapper)
## 
## Attaching package: 'overlapper'
## The following object is masked _by_ '.GlobalEnv':
## 
##     overlap.prob
load("Data/geneLengths.RData")
Universe <- rownames(filterGenes(SEs$acute.msc))
## Loading required package: SummarizedExperiment
## Loading required package: MatrixGenerics
## Warning: package 'MatrixGenerics' was built under R version 4.3.1
## Loading required package: matrixStats
## Warning: package 'matrixStats' was built under R version 4.3.1
## 
## Attaching package: 'matrixStats'
## The following object is masked from 'package:dplyr':
## 
##     count
## 
## Attaching package: 'MatrixGenerics'
## The following objects are masked from 'package:matrixStats':
## 
##     colAlls, colAnyNAs, colAnys, colAvgsPerRowSet, colCollapse,
##     colCounts, colCummaxs, colCummins, colCumprods, colCumsums,
##     colDiffs, colIQRDiffs, colIQRs, colLogSumExps, colMadDiffs,
##     colMads, colMaxs, colMeans2, colMedians, colMins, colOrderStats,
##     colProds, colQuantiles, colRanges, colRanks, colSdDiffs, colSds,
##     colSums2, colTabulates, colVarDiffs, colVars, colWeightedMads,
##     colWeightedMeans, colWeightedMedians, colWeightedSds,
##     colWeightedVars, rowAlls, rowAnyNAs, rowAnys, rowAvgsPerColSet,
##     rowCollapse, rowCounts, rowCummaxs, rowCummins, rowCumprods,
##     rowCumsums, rowDiffs, rowIQRDiffs, rowIQRs, rowLogSumExps,
##     rowMadDiffs, rowMads, rowMaxs, rowMeans2, rowMedians, rowMins,
##     rowOrderStats, rowProds, rowQuantiles, rowRanges, rowRanks,
##     rowSdDiffs, rowSds, rowSums2, rowTabulates, rowVarDiffs, rowVars,
##     rowWeightedMads, rowWeightedMeans, rowWeightedMedians,
##     rowWeightedSds, rowWeightedVars
## Loading required package: GenomicRanges
## Loading required package: stats4
## Loading required package: BiocGenerics
## 
## Attaching package: 'BiocGenerics'
## The following objects are masked from 'package:dplyr':
## 
##     combine, intersect, setdiff, union
## The following objects are masked from 'package:stats':
## 
##     IQR, mad, sd, var, xtabs
## The following objects are masked from 'package:base':
## 
##     anyDuplicated, aperm, append, as.data.frame, basename, cbind,
##     colnames, dirname, do.call, duplicated, eval, evalq, Filter, Find,
##     get, grep, grepl, intersect, is.unsorted, lapply, Map, mapply,
##     match, mget, order, paste, pmax, pmax.int, pmin, pmin.int,
##     Position, rank, rbind, Reduce, rownames, sapply, setdiff, sort,
##     table, tapply, union, unique, unsplit, which.max, which.min
## Loading required package: S4Vectors
## Warning: package 'S4Vectors' was built under R version 4.3.1
## 
## Attaching package: 'S4Vectors'
## The following objects are masked from 'package:dplyr':
## 
##     first, rename
## The following object is masked from 'package:tidyr':
## 
##     expand
## The following object is masked from 'package:utils':
## 
##     findMatches
## The following objects are masked from 'package:base':
## 
##     expand.grid, I, unname
## Loading required package: IRanges
## Warning: package 'IRanges' was built under R version 4.3.1
## 
## Attaching package: 'IRanges'
## The following objects are masked from 'package:dplyr':
## 
##     collapse, desc, slice
## Loading required package: GenomeInfoDb
## Loading required package: Biobase
## Welcome to Bioconductor
## 
##     Vignettes contain introductory material; view with
##     'browseVignettes()'. To cite Bioconductor, see
##     'citation("Biobase")', and for packages 'citation("pkgname")'.
## 
## Attaching package: 'Biobase'
## The following object is masked from 'package:MatrixGenerics':
## 
##     rowMedians
## The following objects are masked from 'package:matrixStats':
## 
##     anyMissing, rowMedians
## 0 rows out of 12827 were removed.
length(Universe)
## [1] 12827
```

12827 genes have module assignment and will be used for the overlap
analysis.

#### 5. Test for overlap enrichment

##### 5.1 GWAS Risk Genes

For each disease, I select the genes that are found in the
universe.

```
GeneVect <- list()

GeneVect$ADHD <- unique(GWAS$ADHD$Gene)[unique(GWAS$ADHD$Gene) %in% Universe]
GeneVect$DM <- unique(GWAS$DM$Gene)[unique(GWAS$DM$Gene) %in% Universe]
GeneVect$BW <- unique(GWAS$BW$Gene)[unique(GWAS$BW$Gene) %in% Universe]
GeneVect$OB <- unique(GWAS$OB$Gene)[unique(GWAS$OB$Gene) %in% Universe]

GWASgenes <- list(NegControlADHD=GeneVect$ADHD, Diabetes=GeneVect$DM, BirthWeight=GeneVect$BW)
```

##### 5.2 Overlaps

```
m <- overlapper::multintersect(ll = MixG_DEGS, ll2 = GWASgenes, universe = rownames(filterGenes(SEs$acute.msc)))
## 0 rows out of 12827 were removed.
dotplot.multintersect(m)
```

##### 5.3 Printing genes and GWAS data

```
GWAS$BW$Table[which(GWAS$BW$Table$MappedGene %in%  c(intersect(MixG_DEGS$MSC_DEGs_DOWN,GWASgenes$BirthWeight) )),] %>% 

DT::datatable(class='hover', rownames=FALSE, caption='MIX G downregulated DEGs associated with Birth weight', filter='top', escape=TRUE, extension='Buttons', options=list(pageLength=30, dom='Bfrtip',columnDefs=list(list(className='dt-center', targets=14, visible=FALSE)),  #as if indexing starts from 0
buttons=list(I('colvis'), c('csv', 'excel')))
                )
```

```
GWAS$DM$Table[which(GWAS$DM$Table$MappedGene %in%  c(intersect(MixG_DEGS$MSC_DEGs_DOWN,GWASgenes$Diabetes) )),] %>% 

DT::datatable(class='hover', rownames=FALSE, caption='MIX G downregulated DEGs associated with Diabetes', filter='top', escape=TRUE, extension='Buttons', options=list(pageLength=30, dom='Bfrtip',columnDefs=list(list(className='dt-center', targets=14, visible=FALSE)),  #as if indexing starts from 0
buttons=list(I('colvis'), c('csv', 'excel')))
                )
```

```
GWAS$BW$Table[which(GWAS$BW$Table$MappedGene %in%  c(intersect(MixG_DEGS$MSC_DEGs_DOWN,intersect(GWASgenes$BirthWeight,GWASgenes$Diabetes)) )),] %>% 

DT::datatable(class='hover', rownames=FALSE, caption='MIX G downregulated DEGs associated with Birth weight and Diabetes (Birth weight GWAS info)', filter='top', escape=TRUE, extension='Buttons', options=list(pageLength=30, dom='Bfrtip',columnDefs=list(list(className='dt-center', targets=14, visible=FALSE)),  #as if indexing starts from 0
buttons=list(I('colvis'), c('csv', 'excel'))))
```

```
GWAS$DM$Table[which(GWAS$DM$Table$MappedGene %in%  c(intersect(MixG_DEGS$MSC_DEGs_DOWN,intersect(GWASgenes$BirthWeight,GWASgenes$Diabetes)) )),]%>% 

DT::datatable(class='hover', rownames=FALSE, caption='MIX G downregulated DEGs associated with Birth weight and Diabetes (Diabetes GWAS info)', filter='top', escape=TRUE, extension='Buttons', options=list(pageLength=30, dom='Bfrtip',columnDefs=list(list(className='dt-center', targets=14, visible=FALSE)),  #as if indexing starts from 0
buttons=list(I('colvis'), c('csv', 'excel'))))
```

#### Savings

```
SessionInfo <- sessionInfo()
Date <- date()
```

```
SessionInfo
```

```
## R version 4.3.0 (2023-04-21)
## Platform: aarch64-apple-darwin20 (64-bit)
## Running under: macOS 15.1
## 
## Matrix products: default
## BLAS:   /Library/Frameworks/R.framework/Versions/4.3-arm64/Resources/lib/libRblas.0.dylib 
## LAPACK: /Library/Frameworks/R.framework/Versions/4.3-arm64/Resources/lib/libRlapack.dylib;  LAPACK version 3.11.0
## 
## locale:
## [1] en_US.UTF-8/en_US.UTF-8/en_US.UTF-8/C/en_US.UTF-8/en_US.UTF-8
## 
## time zone: Europe/Rome
## tzcode source: internal
## 
## attached base packages:
## [1] stats4    stats     graphics  grDevices utils     datasets  methods  
## [8] base     
## 
## other attached packages:
##  [1] SummarizedExperiment_1.30.2 Biobase_2.60.0             
##  [3] GenomicRanges_1.52.1        GenomeInfoDb_1.36.4        
##  [5] IRanges_2.34.1              S4Vectors_0.38.2           
##  [7] BiocGenerics_0.46.0         MatrixGenerics_1.12.3      
##  [9] matrixStats_1.2.0           overlapper_0.99.1          
## [11] ggplot2_3.4.2               dplyr_1.1.2                
## [13] tidyr_1.3.0                 GeneOverlap_1.36.0         
## 
## loaded via a namespace (and not attached):
##  [1] tidyselect_1.2.0        viridisLite_0.4.2       farver_2.1.1           
##  [4] bitops_1.0-7            fastmap_1.1.1           lazyeval_0.2.2         
##  [7] RCurl_1.98-1.14         promises_1.2.0.1        digest_0.6.34          
## [10] mime_0.12               lifecycle_1.0.4         ellipsis_0.3.2         
## [13] VennDiagram_1.7.3       magrittr_2.0.3          compiler_4.3.0         
## [16] rlang_1.1.3             sass_0.4.8              tools_4.3.0            
## [19] utf8_1.2.3              yaml_2.3.7              data.table_1.14.8      
## [22] knitr_1.45              lambda.r_1.2.4          labeling_0.4.3         
## [25] S4Arrays_1.0.6          htmlwidgets_1.6.2       DelayedArray_0.26.7    
## [28] plyr_1.8.9              RColorBrewer_1.1-3      abind_1.4-5            
## [31] KernSmooth_2.23-21      withr_2.5.0             purrr_1.0.2            
## [34] grid_4.3.0              fansi_1.0.4             caTools_1.18.2         
## [37] xtable_1.8-4            colorspace_2.1-0        scales_1.3.0           
## [40] gtools_3.9.5            cli_3.6.2               crayon_1.5.2           
## [43] UpSetR_1.4.0            rmarkdown_2.26          generics_0.1.3         
## [46] rstudioapi_0.15.0       httr_1.4.6              reshape2_1.4.4         
## [49] cachem_1.0.8            stringr_1.5.1           zlibbioc_1.46.0        
## [52] formatR_1.14            XVector_0.40.0          vctrs_0.6.5            
## [55] Matrix_1.5-4            jsonlite_1.8.8          crosstalk_1.2.0        
## [58] plotly_4.10.4           jquerylib_0.1.4         glue_1.7.0             
## [61] DT_0.32                 cowplot_1.1.1           stringi_1.8.3          
## [64] gtable_0.3.3            futile.logger_1.4.3     later_1.3.1            
## [67] shinycssloaders_1.0.0   shinydashboard_0.7.2    munsell_0.5.0          
## [70] tibble_3.2.1            pillar_1.9.0            htmltools_0.5.7        
## [73] gplots_3.1.3.1          GenomeInfoDbData_1.2.10 R6_2.5.1               
## [76] lattice_0.21-8          evaluate_0.21           shiny_1.7.4            
## [79] highr_0.10              futile.options_1.0.1    ggsci_3.0.1            
## [82] httpuv_1.6.11           bslib_0.6.1             Rcpp_1.0.12            
## [85] gridExtra_2.3           xfun_0.42               pkgconfig_2.0.3
```

```
date()
```

```
## [1] "Wed Nov 13 16:52:42 2024"
```
